# RNA structure landscape of *S. cerevisiae* introns

**DOI:** 10.1101/2022.07.22.501175

**Authors:** Ramya Rangan, Rui Huang, Oarteze Hunter, Phillip Pham, Manuel Ares, Rhiju Das

## Abstract

Pre-mRNA secondary structures are hypothesized to play widespread roles in regulating RNA processing pathways, but these structures have been difficult to visualize *in vivo*. Here, we characterize *S. cerevisiae* pre-mRNA structures through transcriptome-wide dimethyl sulfate (DMS) probing, enriching for low-abundance pre-mRNA through splicing inhibition. We cross-validate structures found from phylogenetic and mutational studies and identify new structures within the majority of probed introns (102 of 161). We find widespread formation of “zipper stems” between the 5’ splice site and branch point, “downstream stems” between the branch point and the 3’ splice site, and previously uncharacterized long stems that distinguish pre-mRNA from spliced mRNA. Multi-dimensional chemical mapping reveals examples where intron structures can form *in vitro* without the presence of binding partners, and structure ensemble prediction suggests that such structures appear in introns across the *Saccharomyces* genus. We develop a high-throughput functional assay to characterize variants of RNA structure (VARS-seq) and we apply the method on 135 sets of stems across 7 introns, identifying structured elements that alter retained intron levels at a distance from canonical splice sites. This transcriptome-wide inference of intron RNA structures suggests new ideas and model systems for understanding how pre-mRNA folding influences gene expression.

## Introduction

Introns are widely prevalent features of eukaryotic genomes. Many genes contain long stretches of these non-coding RNA sequences, which are ultimately excised from mRNA precursors through RNA splicing. In the splicing reaction, a large and dynamic macromolecular machine, the spliceosome, precisely recognizes and positions three key intronic sequences termed the 5’ splice site, the branch point, and the 3’ splice site, carrying out the two catalytic steps required for removing introns (Fig. 1A).^1^ Despite their prevalence, the functional roles for many introns remain underexplored. In some cases, intron sequences beyond splice sites regulate gene expression by controlling splicing rates and promoting alternative splicing.^2,3^ In addition, introns can contain functional non-coding RNAs, alter pre-mRNA decay rates, and facilitate the evolution of new genes.^4–10^

**Figure 1:**
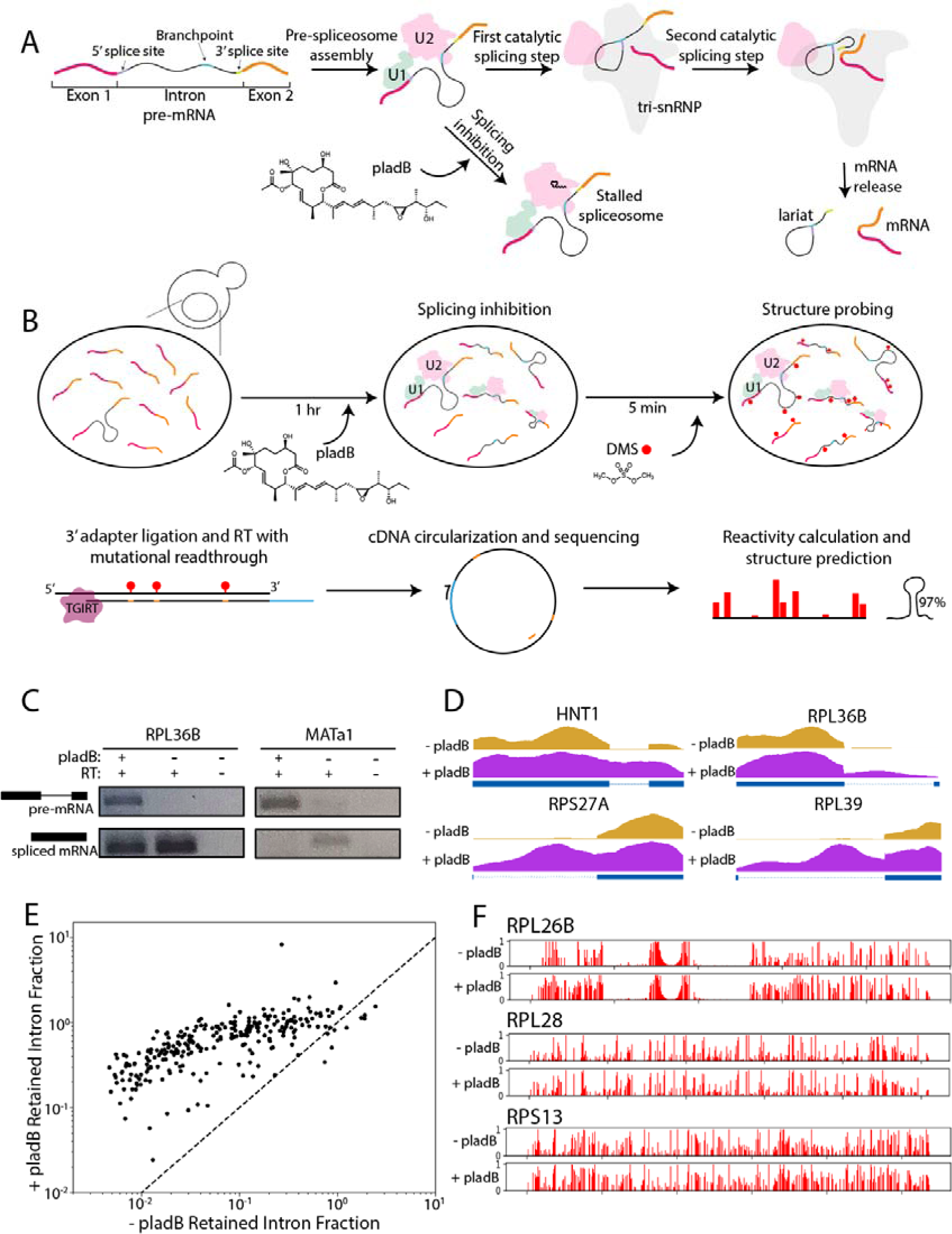
Splicing inhibition by pladB allows for accumulation of pre-mRNA. A) Schematic of RNA splicing. B) Schematic for splicing inhibition followed by DMS-MaPseq experiment. C) Accumulation of pre-mRNA shown for RPL36B and MATa1 by RT-PCR. D) Read coverage across intron-containing pre-mRNA with (purple) and without (orange) pladB treatment. E) Retained intron fraction with and without pladB treatment. Points are plotted on a log scale, and equal retained fraction from both conditions is indicated by a dotted line. F) Comparison of reactivity values for three introns with and without pladB treatment.

Intron RNA sequences are complex macromolecules that occupy an ensemble of secondary structures including single-stranded and double-stranded regions. Intron secondary structure can regulate splice site selection and splicing efficiency, and structure in pre-mRNA can function in numerous other nuclear processes such as RNA editing and RNA end-processing.^11,12^ *S. cerevisiae* provides a useful model system for studying the role of pre-mRNA secondary structures, with the catalytic steps of splicing, the spliceosomal machinery, processes of RNA modification, and RNA decay pathways highly conserved across eukaryotes from *S. cerevisiae* to humans.^1,13,14^ Studies in *S. cerevisiae* have revealed that intron stems called “zipper stems” can link the 5’ splice site to the branch point,^15–18^ and hairpins can lower the effective distance between the branch point and 3’ splice site to enable efficient splicing.^19^ In addition, pre-mRNA structures can bind to protein products translated from their corresponding mRNA to regulate autogenous gene expression control at the level of splicing.^7,20–22^

These studies exemplify different functions for pre-mRNA structures, but how generally these findings extend across the transcriptome is unknown. We are missing a broad experimental survey of structures in introns across the transcriptome of any organism, and functional data that test the role for native intron structures is limited. Though scans for covariation have identified some potentially functional structures,^23^ other structures found by functional studies have not been identified by these scans.^20–22,24^ Moreover, while sequence variants have been used to test some intron structures,^3,25^ the depth of mutagenesis has been limited.^26^ At this stage, the structural landscape and functional roles for intron secondary structures across any transcriptome are only sparsely determined.

*In vivo* chemical probing approaches provide an avenue for obtaining deep structural data on RNA across the transcriptome.^27^ However due to their low abundance, introns typically escape accurate structure detection and quantification by these methods. Here, we use splicing inhibition to enrich for unspliced pre-mRNA in transcriptome-wide structure probing experiments, identifying patterns in accessibility and folding that distinguish structural motifs in *S. cerevisiae* introns from coding regions *in vivo*. We combine this structural information with phylogenetic analysis across a broad set of yeast genomes, and we develop a strategy for high-throughput analysis of variants of RNA structure (VARS-seq) to evaluate levels of unspliced and spliced RNA for 135 sets of stems in 7 introns. Our combined structural, functional, and evolutionary analysis provides an atlas of pre-mRNA structural elements that serves as a foundation for understanding the previously hidden roles that introns play in gene regulation and genome evolution.

## Results

### Transcriptome-wide structure probing with splicing inhibition in *S. cerevisiae*

Structure probing experiments like DMS-MaPseq^27^ can evaluate the formation of RNA secondary structures across the transcriptome. However, since splicing proceeds rapidly in yeast,^28^ the coverage of pre-mRNA is limited in existing DMS-MaPseq datasets for *S. cerevisiae*, preventing the analysis of introns. Recently, *S. cerevisiae* strains with mutations in U2 snRNP component HSH155 (SF3B1 in human) have been developed that sensitize these yeast to splicing inhibition by Pladeinolide B (pladB).^29,30^ Typically, splicing proceeds in two catalytic steps, with sequences in the intron termed the 5’ splice site and the branch point interacting in the first catalytic step, and with the 5’ splice site and another sequence, the 3’ splice site, participating in the second catalytic step (Fig. 1A).^1^ Using a pladB-sensitive yeast strain, we stall the assembling pre-spliceosome (A-complex) before the first catalytic step of splicing has occurred, chemically inhibiting splicing for 1 hour prior to DMS treatment, library preparation, and sequencing (Fig. 1B).

Treatment with pladB led to an accumulation of pre-mRNA, increasing the proportion of unspliced mRNA for RPL36B and MATa1 (Fig. 1C). The data from DMS-MaPseq demonstrated increased intron retention in pladB treated cells for many genes, with read coverage in introns increasing from negligible levels in untreated cells to levels approaching coding regions (Fig. 1D). Most intron-containing genes showed increased intron retention upon pladB treatment, with a median ratio of retained intron fraction (RI fraction) in pladB versus control of 10.5 (Fig. 1E). Only 1 intron had an RI fraction less than 0.05, and 180 introns had an RI fraction over 0.5. Without pladB treatment, 143 out of 288 annotated introns had an RI fraction less than 0.05, and only 28 introns had an RI fraction over 0.5. Longer introns (over 200 nucleotides) and introns in ribosomal protein genes (RPGs) exhibited increased RI fractions upon pladB treatment compared to other introns (Fig. S1A-B). We noted no coverage bias between the ends of introns (Fig. S1C), suggesting that sequenced introns are not partial degradation byproducts of nonsense mediated decay, which predominantly involves XRN1-mediated 5’ to 3’ decay.^31,32^ RI fractions for introns detected with and without pladB treatment are shown in Table S1.

To assess the quality of our DMS-MaPseq reactivity data for introns, we examined control RNAs with known structure. DMS modifies A and C residues, leading to mutational frequencies of 2.7% and 2.3% respectively, whereas unreactive G and U residues exhibited mutation frequency rates closer to background (0.3 - 0.4%) (Fig. S2A). Data for ribosomal RNA (rRNA) aligned closely with prior experiments, with mutational frequency values for the 18S and 25S rRNA highly correlated with values from an earlier DMS-MaPseq study (Pearson correlation’s coefficient r=0.91, Fig. S2B).^27^ Moreover, DMS reactivity values permitted successful classification of accessible (solvent accessible and unpaired) from inaccessible (base-paired) rRNA residues, with an AUC of 0.91 (Fig. S2B). Finally, the reactivity profiles generated for stem-loops in the HAC1 and ASH1 mRNAs align with the well characterized secondary structures for these regions,^27^ with base-paired residues less reactive than positions in loops (Fig. S2C).

### Assessing structure prediction from DMS-MaPseq after splicing inhibition

With data generated from these DMS-MaPseq experiments, we predicted secondary structures using RNAstructure.^33,34^ We additionally assigned confidence estimates for individual helices by performing nonparametric bootstrapping on reactivity values, using an approach that has empirically improved the quality of stem predictions.^35^ To calibrate helix confidence estimate cutoffs, we made DMS-guided structure predictions for structured RNAs in our DMS-MaPseq dataset, including rRNAs, snRNAs, tRNAs, and mRNA segments. Helix confidence estimate thresholds improved the positive predictive value (PPV) and F1 score for stem predictions (Fig. S3A, Fig. S3C). When using reactivity data to make DMS-guided structure predictions with a helix confidence estimate threshold of 70% and minimum stem length cutoff of 5 base-pairs, predicted stems had a PPV of 82.3% (Table S2). DMS-guided structure prediction with helix confidence estimation was able to recover stems with this accuracy even for larger control RNAs like the U1 snRNA (Fig. S3D-E, Table S2). We therefore designated stems with helix confidence estimates > 70% and at least 5 base-pairs as “high confidence stems” and focused subsequent analyses on these stems, aiming to prioritize PPV while maintaining reasonable sensitivity (66.2%, Fig. S3B).

We next assessed the reproducibility of high confidence stems by comparing replicate DMS-MaPseq experiments. Across all 259 introns, we identified 425 high confidence stems, with 331 (77.6%) agreeing between replicates. We noted that introns with higher sequencing coverage had better replicability between experiments, with higher correlation values between their replicate DMS reactivity values (Fig. S4A) and better reproducibility for high confidence stems (Fig. S4B). Indeed, in the 98 introns with higher reactivity correlation between replicates (Pearson’s correlation’s coefficient r > 0.75), 88.1% of high confidence stems agreed between replicates. In addition, isolating high confidence stems enabled reproducibly analyzing portions of intron structures even in cases with lower global r between replicate reactivity profiles. In fact, even in introns with between replicate reactivity correlations ranging from 0.5 to 0.75, 70.0% of high confidence stems matched between replicates. We focused further analyses on the 161 introns with sufficient coverage to obtain between-replicate reactivity r > 0.5, limiting analysis to high confidence stems in these introns (84.5% agreement between replicates). Additionally, we note that highlighted intron structures from DMS-MaPseq in subsequent figures have between-replicate reactivity r > 0.75 and 88.1% agreement of high confidence stems. We include each intron’s replicate correlation and a list of nucleotides in high confidence stems in Table S1.

We identified 366 high confidence stems across the 161 introns with sufficient coverage. The majority (102 of 161) had at least one such high confidence stem. We explored additional structure prediction approaches to identify potential pseudoknots and alternative conformations in these introns using the DMS data, finding that no introns included high confidence predicted pseudoknots, and that all introns tested for multiple conformations were best explained by a single structure (see Supplemental Text).

To test whether the use of PladB resulted in alterations in intron folding, we examined the three introns that met our DMS-MaPseq coverage thresholds both with and without pladB treatment. For these introns, reactivity profiles are broadly similar, with highly reactive positions often shared between conditions (Fig. 1F). However, we noted intervals in RPL26B and RPS13 with elevated differences between reactivity profiles with and without pladB treatment, with some differences beyond replicate variation (Fig. S5). When comparing conditions with and without pladB treatment, the reactivities at A and C residues had correlations of 0.76, 0.93, and 0.84 for the introns in RPL26B, RPL28, and RPS13 respectively. To evaluate whether these differences affected structure prediction, we identified high confidence stems in secondary structures predicted for these three introns with and without pladB treatment. We found that these structures agreed well, with all high confidence stems shared between conditions (Fig. S6), suggesting that the high confidence stems adopted by introns after pladB treatment can be informative for evaluating and discovering intron structures in untreated *S. cerevisiae*.

### Coherence of DMS-guided predictions with previous reports of intron structures in *S. cerevisiae*

Structures have been proposed for some introns in *S. cerevisiae* through functional experiments,^7,15,16,19–22,24^ solved spliceosome structures including intron structures,^25,36^ covariation scans that pinpoint functional base-pairs,^23^ and *de novo* prediction methods based on conservation in sequence alignments.^37^ Using our DMS data, we were able to evaluate the support for the presence of these proposed structures *in vivo*.

We first focused on seven introns that included regulatory structures identified in functional experiments: introns in RPS17B,^15,16^ RPS23B,^19^ RPL32,^20^ RPS9A,^7^ RPS14B,^21,22^ RPL18A^24^ and RPS22B.^24^ Most structures in this class received medium or high support from our DMS data (Fig. 2A-E) with only two exceptions (Fig. S7A-B), in general validating structures found in these experiments that assessed mutants and compensatory mutants (Fig. 2G, see Supplemental Text for details). In contrast, structures identified through computational predictions in Hooks, et al. (2016)^37^ using CMfinder,^38^ RNAz,^39^ and Evofold^40^ showed low to medium support from DMS data (Fig. S8A-G), suggesting that these approaches less reliably identify structures that form *in vivo* compared to functional experiments (Fig. 2G, see Supplemental Text for details.) We note that our DMS-guided stem predictions include some false negatives (33.8% false negative rate on controls, Fig. S3A-B). Additionally, in cases where DMS data do not lead to confident stem prediction, the proposed structure may still be a functional conformation that represents a minor portion of the intron’s structural ensemble or appears only in a particular cellular state.

**Figure 2:**
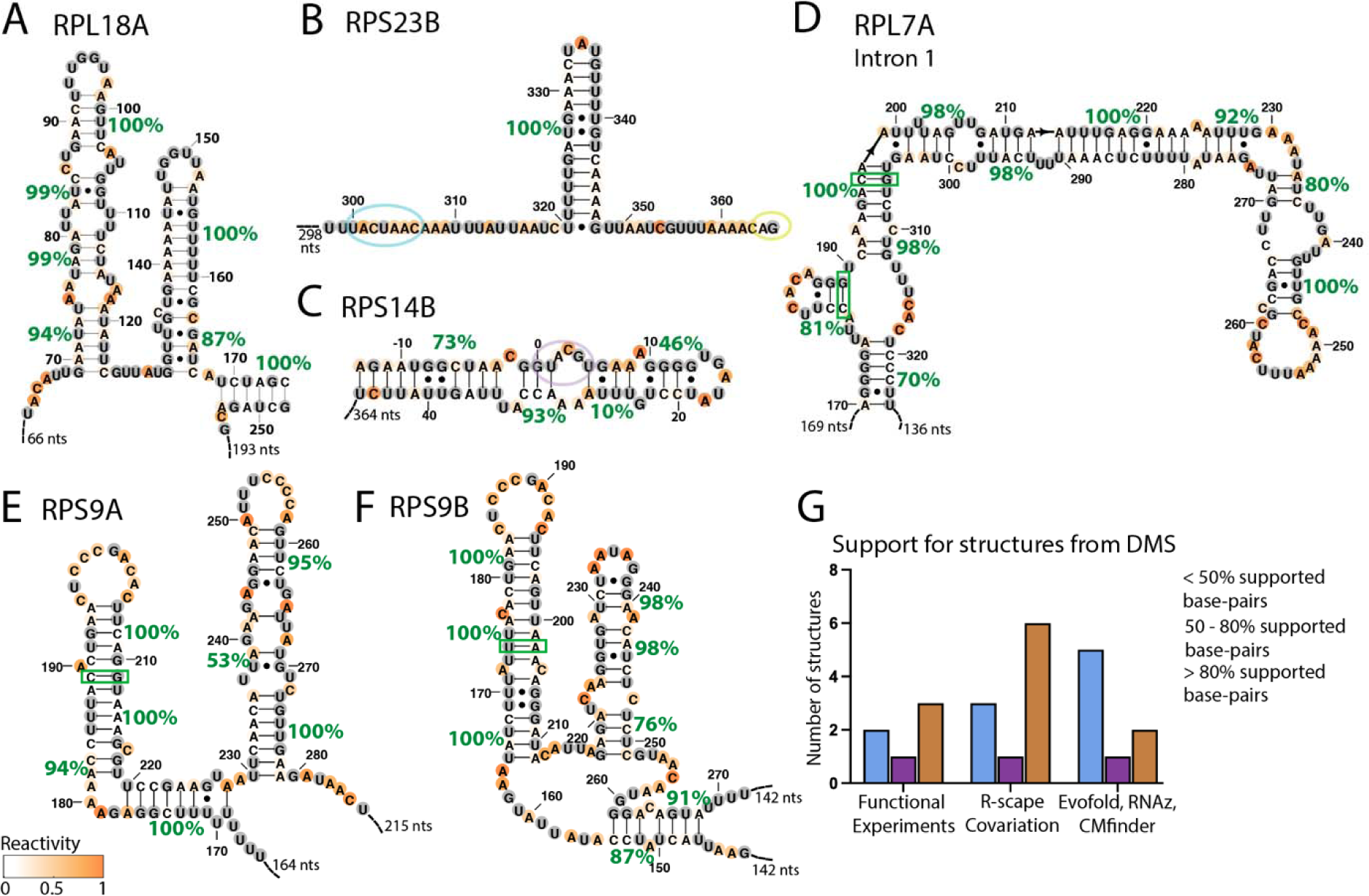
Support from DMS reactivity for *in vivo* formation of control structures and proposed functional structures. Helix confidence estimates and covariation for intron structures reported in A) RPL18A^24^, B) RPS23B^19^, C) RPS14B^21^, D) the first intron in RPL7A, E) RPS9A, and F) RPS9B^7^. Secondary structures are colored by DMS reactivity and helix confidence estimates are depicted as green percentages. 5’ splice site, branch point, and 3’ splice site sequences are circled in purple, blue, and yellow respectively. Covarying base pairs in RPS9A, RPS9B, and RPL7A are indicated as green boxes. G) Summary of the percent of supported base-pairs in structures proposed in prior functional studies, R-scape scans for covariation in multiple sequence alignments (MSAs), and in other approaches using sequence alignments to pinpoint structures (Evofold, RNAz, and cMfinder). A base-pair is supported if included in a stem with over 70% helix confidence estimate, and base-pair support statistics are computed based on all base-pairs in proposed structures (functional experiments, Evofold, RNAz, cMfinder) or significantly covarying base-pairs (R-scape covariation).

We next evaluated structures detected by R-scape, which identifies pairs of residues with significant covariation compared to phylogenetic sequence backgrounds. In particular, we evaluated DMS support for covarying residues identified by R-scape in Gao, et al. (2021),^23^ and we additionally evaluated 11 introns with significantly covarying residues from our own R-scape scan that used less stringent cutoffs (Fig. S9, see Methods). As a positive control for functional secondary structures in introns, we noted that seven of the eight *S. cerevisiae* introns that encode snoRNAs^41^ included significantly covarying residues, suggesting that the remaining introns with covariation may also encode functional structural elements. Since most intron structures with covarying residues were supported by DMS data (Fig. S10, Fig. 2D-F) with one exception (Fig. S7C), DMS data largely validated covariation both in snoRNAs as well as other intron structures (see Supplemental Text for details). The prevalence of DMS-validated covarying residues suggests that covariation from R-scape^42^ can reliably identify structures that form *in vivo* (Fig. 2G). Thus, overall, we found that compared to structures predicted by CMfinder,^38^ RNAz,^39^ and Evofold^40^, structures that were identified through functional experiments and R-scape^42^ covariation were more consistently supported by DMS data.

We also used our DMS data to identify high confidence stems in tRNA introns in *S. cerevisiae*. As these tRNA introns are not processed by the standard spliceosome machinery, we did not expect pladB treatment to encourage their accumulation. Despite this, 10 of the 28 tRNA introns of length at least 30 had sufficient coverage for structure analysis. Each of these introns included at least one high confidence stem (Fig. S11), aligning with prior structure models for pre-tRNA^LEU^, pre-tRNA^PRO^, and pre-tRNA^ILE^ and again confirming structures identified through prior structural and functional experiments.^43,44^

### New structures found by probing *S. cerevisiae* pre-mRNA

Having established above that our DMS-guided structure predictions are coherent with previously identified intron structures, we next identified new intron structures and enriched structural features from our data. First, the *in vivo* probing supports the widespread formation of zipper stems, i.e., stems that reduce the distance between the 5’ splice site and branch point. Potential zipper stems have been noted in various introns,^17^ and a zipper stem in RPS17B has been shown to be essential for efficient splicing.^15,16^ To identify zipper stems across introns, we sought to precisely define the positional constraints on zipper stem formation, modeling intronic stems with various linker lengths in the context of the A-complex spliceosome using Rosetta^45^ (Fig. S12A-B, see Methods). Based on this modeling, we defined zipper stems as the longest stem with between 42 and 85 total nucleotides linking the stem, the 5’ splice site, and branch point sequences. We find that high confidence zipper stems (including at least 6 base-pairs with at least 70% helix confidence estimate) are predicted for 42 of 161 introns with sufficient read coverage (Fig. 3A).

**Figure 3:**
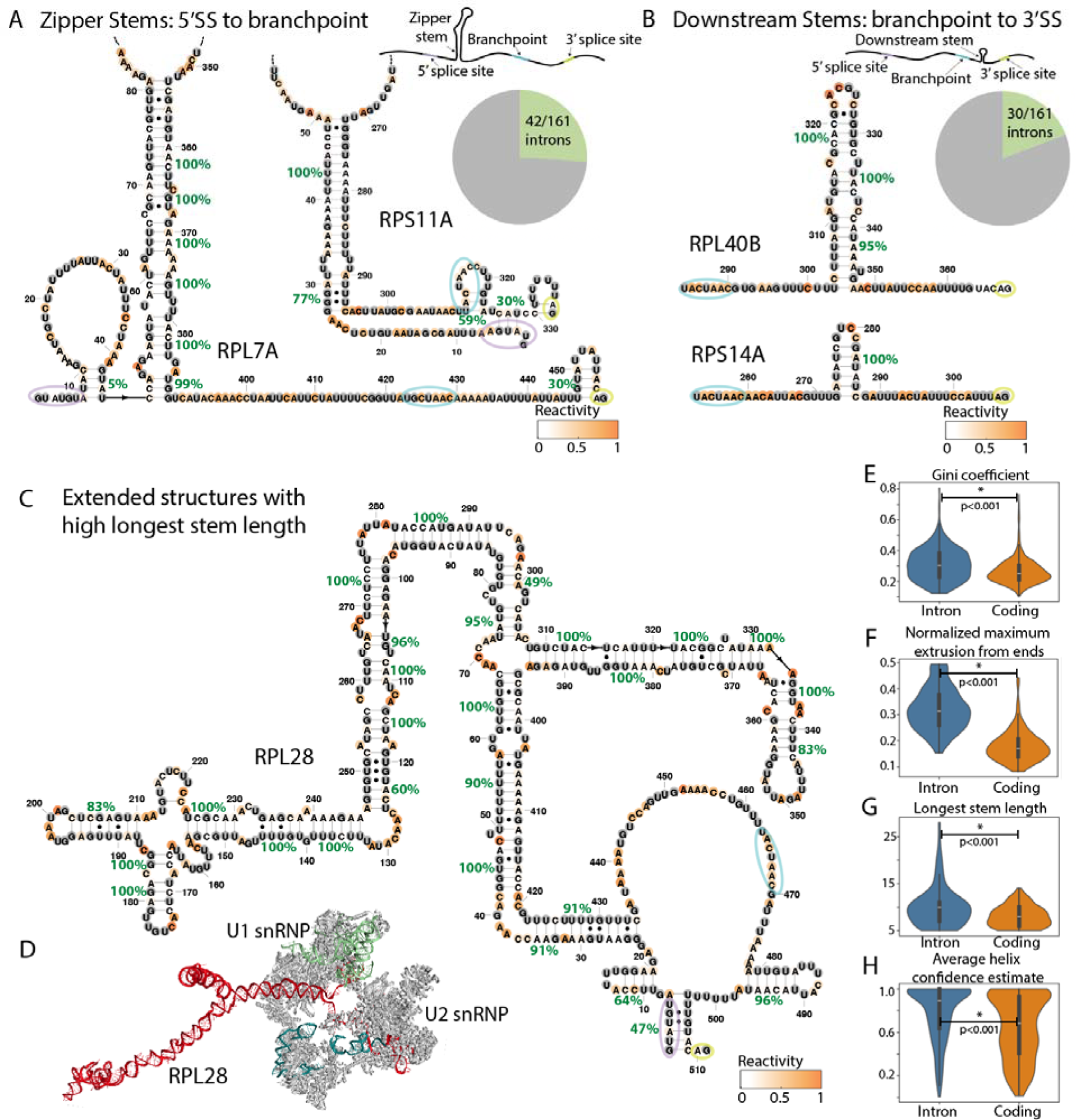
Structural insights from DMS probing of *S. cerevisiae* introns. A) Reactivity support for zipper stems in RPL7A and RPS11A. B) Reactivity support for stems connecting the branch point and 3’ splice site in RPL40B and RPS14A. C) Secondary structure for RPL28 predicted by RNAstructure guided with DMS reactivity. 5’ splice site, branch point, and 3’ splice site sequences are circled in purple, blue, and yellow respectively. D) Top-scoring 3D model for the RPL28 intron in the context of the A complex spliceosome (PDB ID: 6G90)^99^, modeled using the secondary structure derived from DMS-MaPseq. E)-H) Comparisons of secondary structure features between introns and coding regions were computed using Wilcoxon ranked sum tests to compare classes.

We next noted the presence of stems between the branch point and 3’ splice site across introns, which we term “downstream stems”. These stems are proposed to decrease the effective distance between the branch point and 3’ splice site to facilitate splicing.^19^ We find that downstream stems (at least 6 base-pairs in the stem with at least 70% helix confidence estimate) are present in 30 of 161 introns (Fig. 3B). As for zipper stems (Fig. 3A), the reactivity profiles for downstream stems (e.g. in RPL40B, RPS14A, and RPS23B) show higher reactivity in loop or junction residues than in base-paired positions, supporting their *in vivo* formation (Fig. 2B, Fig. 3B).

DMS reactivity for some introns suggest new elaborate extended secondary structures with long stems and multiway junctions. For instance, in the case of RPL28, in addition to a zipper stem connecting the 5’ splice site to the branch point and a downstream stem between the branch point and 3’ splice site, the pre-mRNA includes a three-way junction and long stems of over 10 base-pairs through the length of the intron (Fig. 3C). We speculate that these secondary structures allow internal stems to extend beyond the spliceosome, potentially poised to interact with other factors. In fact, when building a 3D model of the RPL28 intron in the context of the A-complex spliceosome, we observe that introns’ stems can extrude from the core spliceosome, potentially enabling binding partners to avoid steric clashes with the splicing machinery (Fig. 3D). We note that the 3D modeling approach here makes simplifying assumptions and samples only a few of many possible conformations (see Methods). However, sampled conformations suggest that intron structures can extend beyond the length of the spliceosome even for shorter introns like RPL36B (238 nucleotides), facilitated by stems through the length of the intron (Fig. S12C).

We next analyzed the placement of high confidence stems in introns relative to the positions of canonical and cryptic splice sites. To evaluate structures surrounding canonical splice sites, we first identified sequence intervals in pre-mRNA surrounding the 5’ splice site, branch point, and 3’ splice site where pre-mRNA structures would be expected to clash with the spliceosome (see Methods). We noted that these intervals were significantly depleted of high confidence stems when generating structures for introns with surrounding pre-mRNA sequence context (Fig. S13A). However, protection of these intervals by high confidence stems was not apparently correlated with fraction of spliced constructs, suggesting a more dominant role for other factors (Fig. S13B). We next identified cryptic splice sites across introns, expecting that these sites may be occluded by structure. Indeed, in multiple introns, downstream stems between the branch point and 3’ splice site occluded cryptic 3’ splice sites (Fig. S13C), aligning with prior work demonstrating that a RPS23B downstream stem blocks a cryptic 3’ splice site (Fig. 2B).^19^ Across introns, we noted that cryptic 3’ splice sites were enriched for high confidence stems (Fig. S13D), suggesting a potential role for these stems in enforcing splicing fidelity by repressing the use of incorrect 3’ splice sites.

### Introns are more structured than coding regions and unspliced decoys

Our DMS-guided structure predictions reveal that intron secondary structures have distinct properties compared to those of mRNA coding regions. As found in mammalian compartment-specific structure probing,^46^ intron regions have higher Gini coefficients, a measure quantifying the extent to which reactivity values diverge from an even distribution (Fig. 3E). Therefore, introns include more non-random secondary structure elements compared to coding regions. Additionally, intron structures predicted using DMS data extend further from their sequence endpoints than coding regions, which we measure as the normalized maximum extrusion from ends (MEE) (Fig. 3F, see Methods). Introns additionally include longer stems with helix confidence estimates of at least 90%, with some high confidence stems extended to over 20 base-pairs (Fig. 3G). Finally, intron stems with at least 6 base-pairs have higher helix confidence estimates on average, suggesting that reactivity values better support intron secondary structures (Fig. 3H). These conclusions were robust to extending analysis to introns with higher between-replicate reactivity correlation or when analyzing data from DMS-MaPseq replicates separately (Fig. S14).

The secondary structure patterns enriched in introns may distinguish spliced introns from other transcribed sequences that do not splice efficiently. To explore this possibility, we assembled a set of unspliced decoy introns in *S. cerevisiae* that included consensus 5’ splice site, branch point and 3’ splice site sequences in positions that matched canonical introns’ length distributions (Fig. S15A). DMS-guided structure predictions for authentic intron sequences enriched for more stable (lower ΔG) zipper stems, more stable downstream stems, longer stems, higher maximum extrusion from ends, and higher secondary structure distances from the 5’ splice site to the branch point (Fig. S15B). In contrast to spliced introns, the DMS-guided secondary structures of unspliced decoys were not enriched for these features (Fig. S15B). We conclude that compared to coding regions and unspliced decoys, introns in *S. cerevisiae* adopt extended secondary structures with long stems supported by DMS data.

### Comparing *in vivo* intron RNA folding with *in vitro* folding

To determine whether intron structures observed *in vivo* can form intrinsically in RNA folded *in vitro* even when potential protein binding partners are missing, we probed isolated introns transcribed *in vitro* with DMS. For this, we used mutate-and-map readout through next-generation sequencing (M2-seq^47^), which can identify base-pairing residues in addition to providing average per-residue accessibility data. We chose five introns from DMS-MaPseq that included zipper stems and had lengths short enough to map completely with *in vitro* M2-seq. In the case of the introns in QCR9 and RPL36B, Z-score plots from *in vitro* M2-seq included off-diagonal signals that are indicative of the presence of stems, and high base-pairing probabilities support the formation of these stems as these introns’ primary *in vitro* structure (Fig. 4A-B and Fig. S16A-B, see Supplemental Text for details). Secondary structures predicted from M2-seq agree with the stems observed *in vivo*, with most high confidence stems shared between these structures (Fig. 4C-D, Fig. S16C-D). For the introns in RPS11A, RPL37A, and RPS7B, though M2-seq Z-scores did not include visually apparent off-diagonal signals, helix confidence estimates computed using M2-seq data showed support for the stems observed *in vivo* (Fig. S17). We additionally assayed intron structures *in vitro* by refolding RNA extracted from yeast and probing accessible residues with DMS (see Methods). However, since only 3 introns reached between-replicate reactivity correlation of r > 0.75 from this *in vitro* probing experiment (Fig. S18), we focused our analyses on cases studied with *in vitro* M2-seq. Our *in vitro* M2-seq results suggest that intron sequences can form structures found *in vivo*, even when outside the context of the nucleus and without protein binding partners.

**Figure 4.**
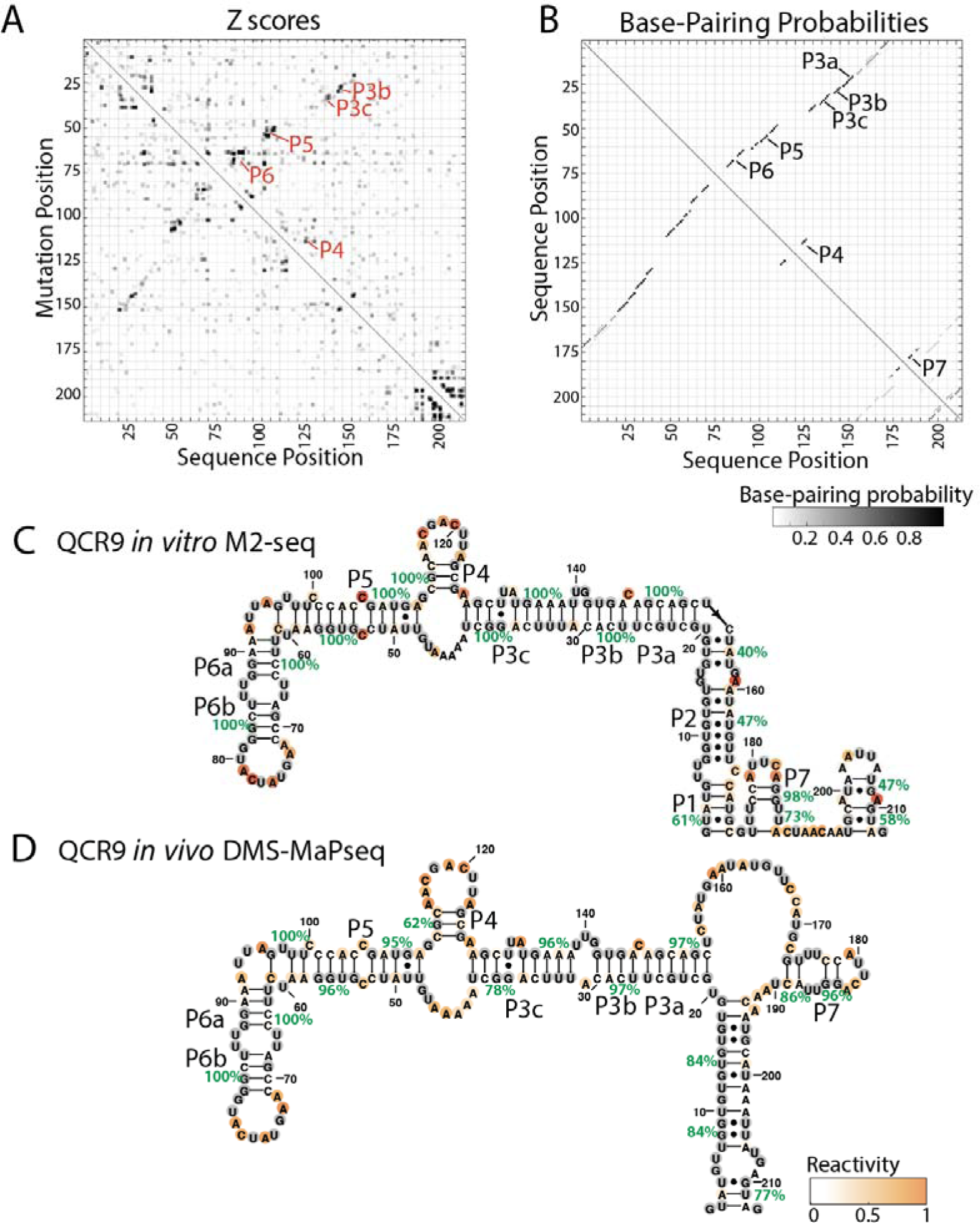
Comparing *in vivo* and *in vitro* folding of intron RNA structures. A) *In vitro* M2-seq Z scores for the intron in QCR9, with peaks representing helices annotated in red. B) *In vitro* chemical reactivity base-pairing probabilities for QCR9 using 1D and 2D chemical reactivity from M2-seq, with peaks representing helices annotated in black. Secondary structure predictions guided by 1D and 2D DMS probing data for the intron in QCR9 C) from *in vitro* M2 seq, and D) from *in vivo* DMS-MaPseq.

### Structural landscape for *S. cerevisiae* introns

To understand the distribution of structural patterns in introns across the yeast transcriptome, we clustered introns based on their structural features. For each intron with sufficient DMS-MaPseq coverage, we assembled a full set of features using probing data: zipper stem and downstream stem free energy (see Methods), the maximum extrusion from ends, the length of the longest stem, the average helix confidence estimate from bootstrapping for stems in the intron, the maximum Gini coefficient window, and the accessibility (average DMS reactivity) across the 5’ splice site, branch point, and 3’ splice site. In Table S1, we tabulate these secondary structure features, DMS-guided secondary structures, and lists of nucleotides in high confidence stems across all *S. cerevisiae* introns.

The clustered heatmap (Fig. 5) depicts the global distribution of secondary structure features across all introns with sufficient sequencing coverage. The first class of introns includes 12 that have both a zipper stem and downstream stem, with high average helix confidence estimates (Class 1, blue in Fig. 5). Class 2 includes 30 introns with zipper stems, and within this class, introns with more stable zipper stems clustered together (orange Fig. 5). The next class includes 18 introns with downstream stems; this class included lower longest stem lengths compared to introns with zipper stems (Class 3, yellow in Fig. 5). In *S. cerevisiae*, intron lengths are bimodal, with a set of shorter introns (shorter than 200 nucleotides, with most around 80 nucleotides) and a set of longer introns (with most between 400 and 500 nucleotides).^48^ Our structure probing data had sufficient coverage for 45 short introns and 116 long introns. Of the long introns, a sizable fraction include either a zipper stem or downstream stem in Class 1-3, with 51.7% of long introns including at least one of these motifs. Class 4 includes structured short introns with at least one high confidence stem (green in Fig. 5). With only 5 introns in this class, a minority of short introns show signal for high-confidence structure. The fifth class includes 13 structured long introns that do not meet the criteria for zipper stems or downstream stems, and yet include structures with high helix confidence, long stems, and windows with high Gini coefficients (Class 5, pink in Fig. 5). In contrast, the next class includes long introns which in some cases do not include high Gini coefficients or do not include long stems, indicating that introns in this class tend to be less structured than Class 5 (Class 6, brown in Fig. 5). Class 7 includes most short introns, which tend to be depleted of stem structures, without zipper stems, downstream stems, high helix confidence estimates, or high Gini coefficient windows (purple in Fig. 5). Finally, Class 8 includes a small set of long introns depleted of structure, including only short stems with low helix confidence (red in Fig. 5).

**Figure 5:**
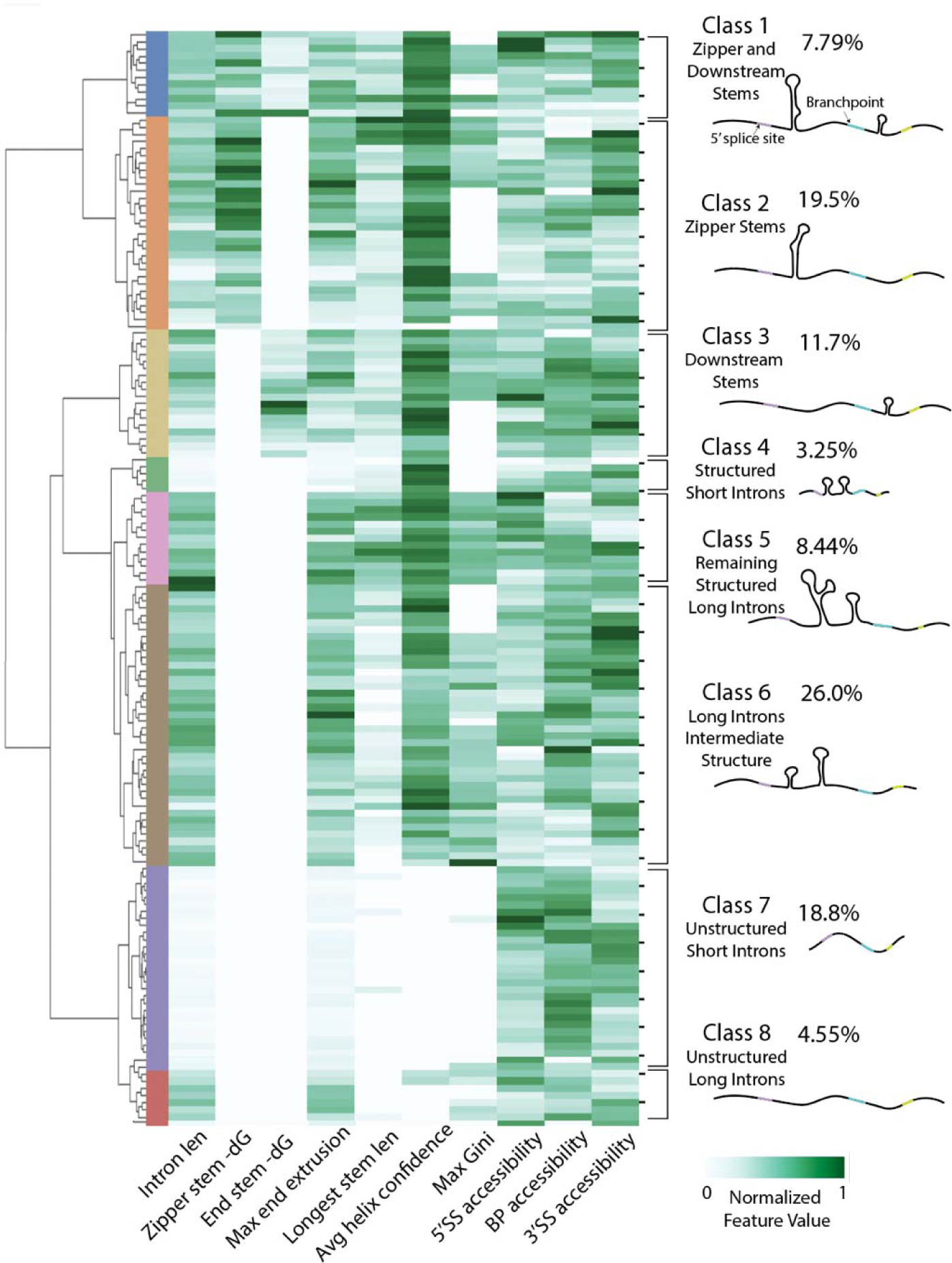
Structural landscape for *S. cerevisiae* introns. Heatmap and dendrogram summarizing intron structural classes, with hierarchical clustering based on secondary structure features. In addition to the features displayed on the heatmap, flags indicating whether zipper stems and downstream stems were present were included as features for hierarchical clustering.

### A high-throughput assay to test the function of intron stems

To assess the influence of intron structures on gene expression, we developed a high-throughput assay for evaluating variants of RNA structure (VARS-seq). VARS-seq includes a variant design pipeline for interrogating structures of interest, a functional assay with a sequencing readout, and analysis methods for measuring effects. Here we use VARS-seq to measure spliced and unspliced mRNA levels for intron structure variants, anticipating that these structures may influence splicing or pre-mRNA decay rates. Structures that slow pre-mRNA decay or splicing could lead to an accumulation of unspliced mRNA, whereas structures that increase splicing rates could in turn increase spliced mRNA levels (Fig. 6A). We chose 7 structured introns to assay with VARS-seq, including introns in Class 1 and Class 2 with zipper stems (RPL7A, QCR9, RPL36B, and RPL28), covarying base-pairs (RPS9A in Class 5, RPS9B in Class 5, and RPL7A in Class 2), and long stems that distinguished pre-mRNA from coding RNA.

**Figure 6:**
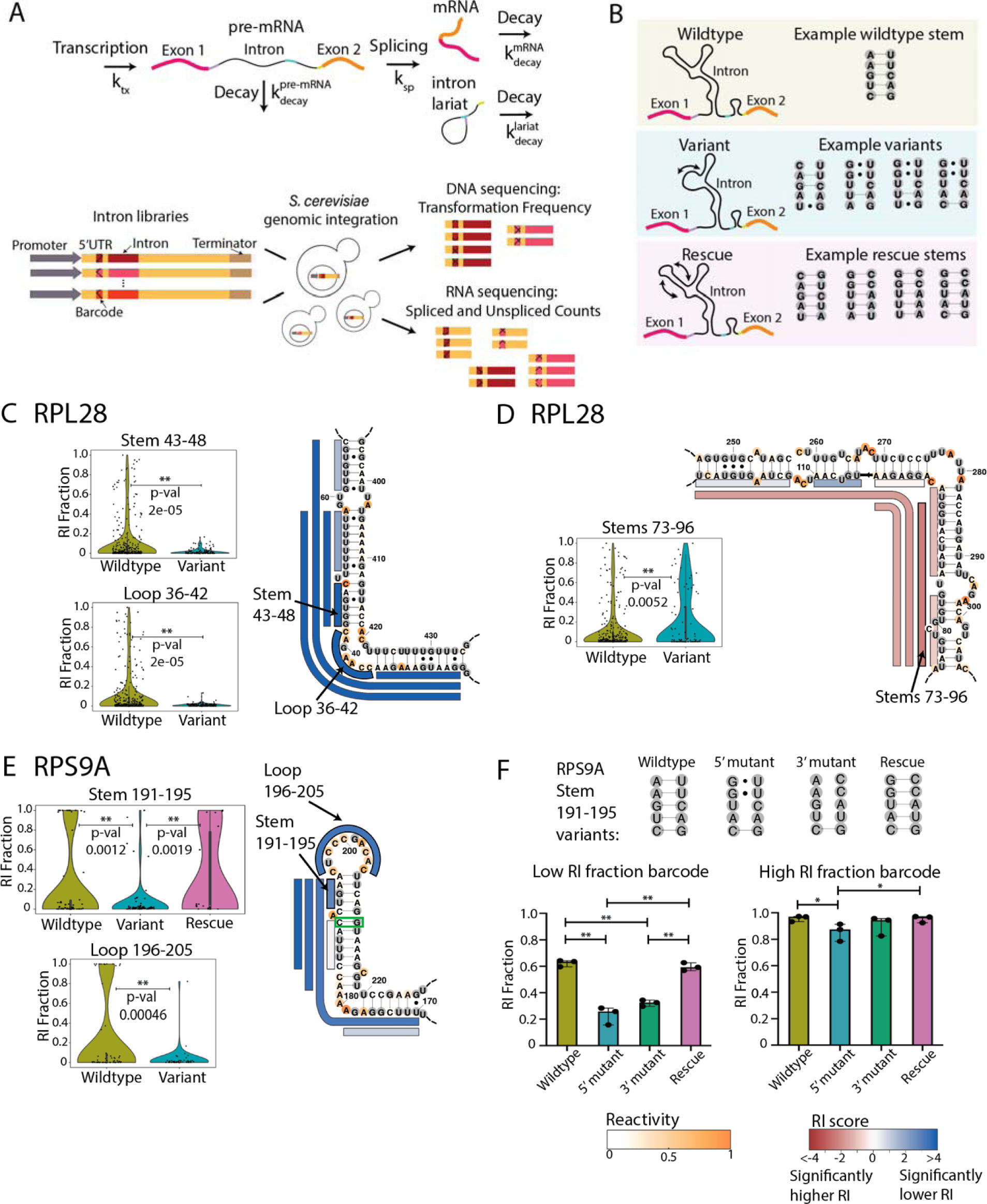
High-throughput structure-function assay for evaluating intron stem variants. A) Schematic for an example library design for one intron stem, with variants disrupting the stem and rescue sequences restoring base-pairing. B) Experiment overview for structure-function experiment. C)-E) The effects of structure variants on retained intron fraction for two regions of the RPL28 intron in C)-D) and for stems in RPS9A in E). For a given stem set or loop, violin plots depict data for the wildtype sequence and all variant sequences, with black points for each unique barcode. Data for rescue sequences are shown when included in the intron library. p-values are indicated for comparisons between wildtype and variant sequence sets, and between variant and rescue sequence sets. p-values are computed using two-sided permutation tests for the difference in mean statistic. Secondary structures are colored by reactivity data. Bars alongside the secondary structure indicate stem and loop disruption sets, with each bar representing a set of variant sequences mutating nucleotides across the full extent of the bar. These bars are colored by the retained intron (RI) score for the corresponding stem or loop disruption set. The RI score is computed as the negative log(p-value) when comparing RI values between wildtype and variant sequences, and the sign is used to indicate the effect direction, with positive values (shown as blue) for lower variant RI compared to wildtype, and negative values (shown as red) for higher variant RI compared to wildtype. The green box in E) indicates a significantly covarying residue. F) RI fractions as measured by RT-qPCR for individual strains containing a set of wildtype, variant, and rescue sequences for RPS9A stem 191-195 (top), with data shown for strains constructed with two different barcode sequences with three biological replicates (bottom). p-values are computed with 2-way ANOVA tests with multiple comparisons.

For each intron, we designed variants that systematically mutated secondary structure elements and rescued them with compensatory mutations where possible (Table S3). More specifically, for each target set of stems of interest, we chose variants that were computationally predicted to disrupt base-pairing within the stem set while maintaining base-pairing elsewhere in the structure (Fig. 6B, Fig. S19A-B). Where technically feasible within the length limits of library construction (see Methods), we also included rescue sequences that were computationally predicted to restore the native secondary structure, with compensatory mutations for each primary mutation (Fig. 6B, Fig. S19A-B). We tested 4-8 distinct intron variants for each target set of stems and loops, with around 200 variant sequences for each intron (Table S3). We integrated these intron variant libraries into the yeast genome^49^ in their native gene context (Fig. 6C, Fig. S19C), installing unique random barcodes upstream of intron variants to help match spliced RNAs to their original pre-mRNA variant (Fig. S19C-D, Fig. S20). With multiple unique barcodes and variant sequences assigned to each set of stems, we were able to observe subtle effects on gene expression due to changes in intron structure (see Supplemental Text for details). Using our RNA sequencing data, we computed two key readouts for each barcode-variant combination: the retained intron (RI) fraction (fraction of total RNA that is unspliced) and the normalized mRNA level (spliced mRNA levels normalized by representation in genomic DNA libraries from the same cell populations).

Across the 7 tested introns, we detected statistically significant effects on intron retention (RI) and mRNA levels for many of the tested sets of stems (52 of 136) and loops (4 in 15) (Fig. S21). Some stem disruptions significantly decreased RI levels (Fig. 6C), while others significantly increased them (Fig. 6D). For many introns, distinct variant sequences designed to disrupt overlapping sets of stems produced similar effects in our assay, providing a useful cross-check for our approach while also suggesting larger functional domains (Fig. S21A-D). For instance, variants disrupting all four sets of stems that included nucleotides 45-48 in the RPL28 intron significantly decreased RI (Fig. 6C, Fig. S21A). Mutations in junction nucleotides 36-42 produced a similar effect (Fig. 6C), indicating that nucleotides 36-48 in the wildtype RPL28 intron help to reduce splicing efficiency or slow pre-mRNA decay. Similarly, all mutations to nucleotides 73-96 of the RPL28 intron show increased RI (Fig. 6D), suggesting that the wildtype nucleotides promote splicing or pre-mRNA decay. Unexpectedly, zipper stem variants in the RPL28 intron (Fig. 6C) lowered RI, suggesting that not all zipper stems promote splicing by co-localizing splice sites.^16^ Combined with the secondary structure predictions from DMS-MaPseq, these results demonstrate that intronic mutations can influence splicing and gene expression, even when structurally distant from splice sites (Fig. 6D, Fig. S21).

In cases where we were able to generate compensatory mutations, stem disruption and rescue variants pinpoint functional intronic structural elements. For instance, in the case of a set of stems in RPL36B, sequence variants in this region reduced normalized mRNA levels and rescue variants restored higher mRNA levels (Fig. S22A), implicating the RNA structure rather than its primary sequence as the functional element. In the case of RPS9A, we identified a functional stem where disruptions reduced RI and stem rescue restored RI (Stem 191-195, Fig. 6E). An analogous stem-loop is present in the RPS9B intron with similar effects on RI levels (Stems 165-183, Fig. S21B). These stem-loops in RPS9A and RPS9B likely influence gene expression by inhibiting splicing or pre-mRNA decay, aligning with the prediction for covariation in these stems (Fig. 2H-I) and corroborating a prior study^3^ on the RPS9A intron.

To validate findings from VARS-seq, we constructed strains containing individual variant and rescue sequences for stems in RPL36B and in RPS9A, and we measured RI and spliced mRNA levels with RT-qPCR. In the case of RPS9A, RT-qPCR data from individually constructed variant and rescue sequences recapitulated structure effects from VARS-seq along with the effects of barcode sequences (Fig. 6E-F). Indeed, when assessing a barcode that yielded low wildtype RI (left, Fig 6F), variant sequences lower RI and rescue sequences restore RI to wildtype levels with RT-qPCR. In contrast, variant and rescue sequences designed for stems in RPL36B (Fig. S22A) did not show significantly different normalized mRNA levels by RT-qPCR (Fig. S22B-C). In this case, it is possible that small effects found by aggregating data across variant and barcode sequences through VARS-seq could not be discerned when analyzing individual variant sequences due to increased noise in RT-qPCR. In general, when comparing between VARS-seq and RT-qPCR data, we noted higher absolute RI and normalized mRNA levels read out from RT-qPCR, perhaps due to differences in length biases of the two assays (Fig. 6E-F, Fig. S22). Due to these differences in dynamic range, effects for higher RI barcodes on RPS9A’s intron are not visible with RT-qPCR (right, Fig. 6F), and significant zipper stem effects in RPL36B from RT-qPCR were not visible in the detection range for VARS-seq (Fig. S22B-C). We therefore suggest using VARS-seq to generate hypotheses on relative rather than absolute splicing differences, and we suggest verifying individual examples of interest with independent assays. To enable further dissection of effects of intron stem variants, we summarize all variant and rescue comparisons from VARS-seq in heatmaps in Fig. S23 and we include per-barcode spliced and unspliced RNA counts in Table S4.

### Computational prediction of enriched structural patterns across the *Saccharomyces* genus

Our transcriptome-wide structure mapping provides evidence for widespread structure in *S. cerevisiae* introns, and with VARS-seq, we find that intron structures can impact gene expression regulation. To extend our observations to other species in the *Saccharomyces* genus, we turned to computational structure prediction. We first evaluated whether *de novo* secondary structure prediction could recapitulate structural features observed through DMS-guided structure analysis, comparing structural features between introns and length-matched controls (Fig. 7A, Fig. S24). For *de novo* structure prediction, as a first approach, we predicted minimum free energy structures,^33^ and as a second approach, we generated secondary structure ensembles through stochastic sampling of suboptimal structures,^50^ which has previously enabled the study of structural patterns in introns.^16,19^ With either of these approaches, structural features from *de novo* secondary structure prediction were largely consistent with patterns from DMS-guided structures for *S. cerevisiae* (Fig. 7B, Fig. S25, see Supplemental Text for details).

**Figure 7:**
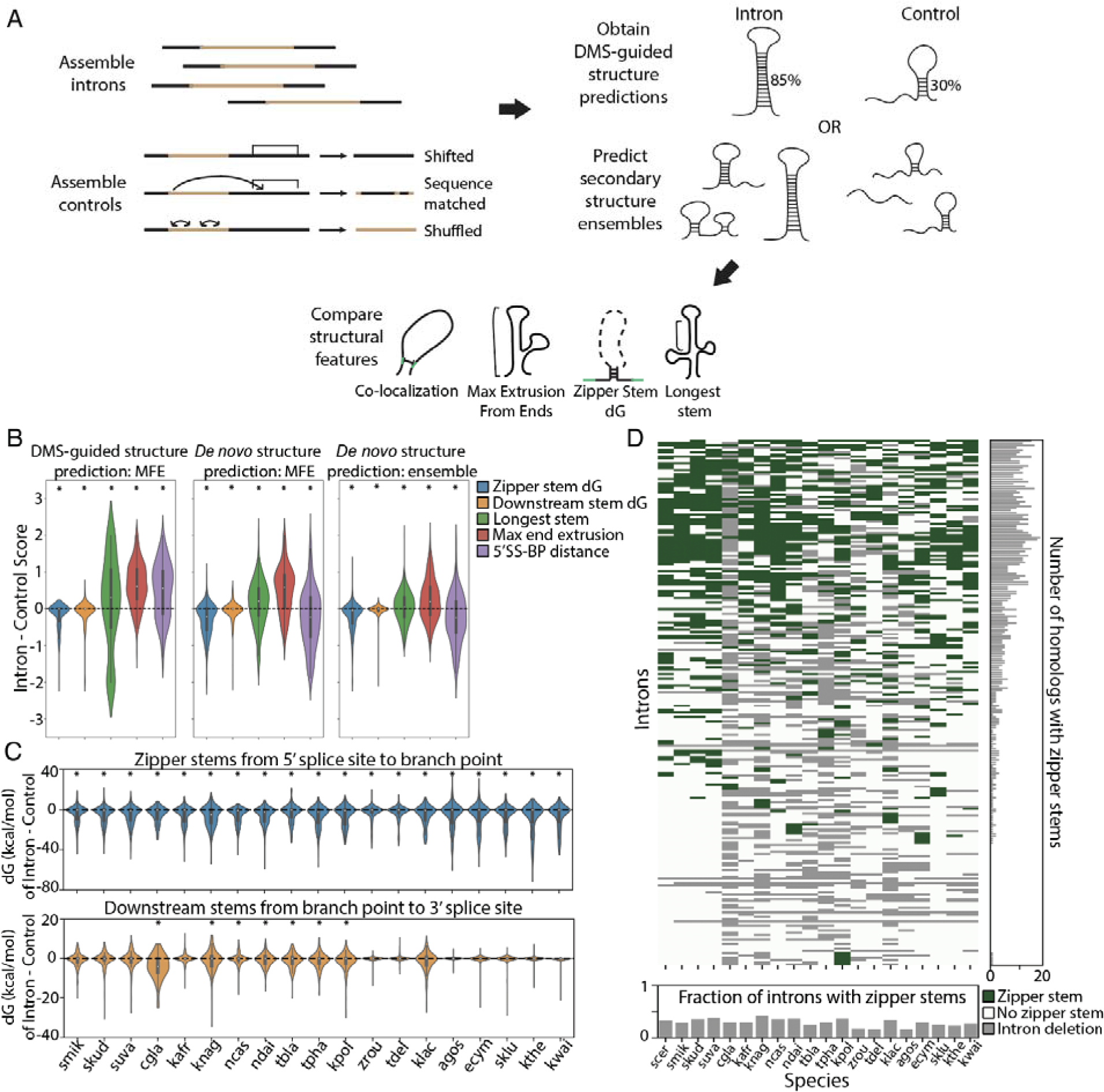
*De novo* secondary structure feature prediction for *S. cerevisiae* and the *Saccharomyces* genus. A) Workflow for comparisons between introns’ secondary structure ensembles and those of control sequences. B) Comparison of secondary structure feature enrichment between introns and control sequences for DMS-guided structure prediction (with folding engine RNAstructure, comparing introns to shifted genomic controls), *de novo* MFE structure prediction (with folding engine RNAstructure, comparing introns to shifted genomic controls), and *de novo* ensemble structure prediction (with folding engine Vienna 2.0, comparing introns to shuffled sequence controls). *p-value < 0.01 by Wilcoxon ranked-sum test. C) Difference in zipper stem (top) and downstream stem (bottom) dG between introns and shuffled sequence controls for introns in the *Saccharomyces* genus, using Vienna 2.0 ensemble predictions. *p-value < 0.01 by Wilcoxon ranked-sum test. D) Distribution of zipper stems across introns in the *Saccharomyces* genus. Heatmap green values indicate a predicted zipper stem, white indicates no predicted zipper stem, and gray values indicate deleted introns. Ohnologous introns are combined into a single row, and a zipper stem is annotated if present in either ohnolog. The species represented in this figure are abbreviated: *E. gossypii* (agos), *C. glabrata* (cgla), *E. cymbalariae* (ecym), *K. africana* (kafr), *K. lactis* (klac), *K. naganishii* (knag), *V. polyspora* (kpol), *L. thermotolerans* (kthe), *L. waltii* (kwal), *N. castellii* (ncas), *N. dairenensis* (ndai), *S. kudiavzevii* (skud), *S. mikatae* (smik), *S. uvarum* (suva), *T. blattae* (tbla), *T. delbrueckii* (tdel), *T. phaffii* (tpha), *Z. rouxii* (zrou).

We next made *de novo* structure predictions across species in the *Saccharomyces* genus, focusing on the 20 species with curated intron alignments from Hooks et al. (2014).^51^ Enrichment for numerous secondary structure patterns is conserved in introns across the *Saccharomyces* genus. In particular, when compared to shuffled sequence controls, introns across the genus include more stable zipper stems and downstream stems (Fig. 7C), along with higher maximum extrusion from ends and shorter distances between the 5’ splice site and branch point (Fig. S26A). Furthermore, many of these features remain enriched when comparing each species’ introns to phylogenetic controls, which include random sequences constructed to match the mutation and indel frequency between an intron and its homologous *S. cerevisiae* intron (Fig. S26B).

Secondary structure patterns are maintained across *Saccharomyces* species despite extreme sequence-level divergence between introns, suggesting that these structures have conserved functions. Complete intron deletions between these species are common, with many introns having orthologs in only a subset of the *Saccharomyces* species (Fig. S26C), and most intron regions have low sequence conservation between species, with most intron sequences 50-60% conserved between these species (Fig. S26D). Some key functional intervals in introns are more conserved across the *Saccharomyces* species, with higher sequence conservation (72.3% to 83.6%) across the seven snoRNAs that are included in these intron sequence alignments. In contrast, zipper stem regions in *S. cerevisiae* introns diverge significantly between *Saccharomyces* species, with most zipper stems only 20-30% conserved in primary sequence. Strikingly, although most species in the genus include zipper stems in 20-40% of their introns, this structural motif appears in disparate introns across species, with many zipper stems only present in a small number of orthologous introns (Fig. 7D). Therefore, in some introns, zipper stems may have evolved separately across species, pointing to a potential functional role for these stems. It is possible that functional secondary structures in these regions have been challenging to find by covariation due to high intron sequence divergence in closely related fungal genomes.

## Discussion

RNA structures play critical roles in regulating a wide array of nuclear processes including transcription, RNA modification and editing, and splicing. Intronic RNA structures are poised to participate in these processes, for instance by altering splicing kinetics, by changing RNA decay rates, or by interacting with other nuclear factors and performing orthogonal regulatory functions. Here, we sought to understand the structural landscape of introns in *S. cerevisiae*, a model system for eukaryotic splicing. First, we evaluated the presence of structural patterns through transcriptome-wide DMS probing while enriching for unspliced RNA, revealing extended secondary structures in introns that distinguished these regions from coding mRNA. These structural data allow for clustering introns into eight classes, with most falling into classes that contain introns with zipper stems or downstream stems (present in 51.7% of the long introns), long introns with intermediate structure, and unstructured short introns. In Fig. S27 and Fig. S28, we display secondary structures and reactivity profiles for all introns from DMS-MaPseq, grouped into these classes. With a high-throughput structure-function assay, VARS-seq, we evaluated the roles for these structures through deep mutagenesis of 7 introns, finding structural elements that influence spliced and unspliced mRNA levels. Finally, through computational structure prediction, we identified signals for structure in introns across the *Saccharomyces* genus.

Structure probing experiments have enabled the measurement of RNA structure transcriptome-wide, but these experiments often lack sufficient coverage to provide information for low abundance transcripts including many unspliced pre-mRNA molecules. Specific low abundance introns can be probed by target-specific enrichment^27,52^ or nuclear RNA enrichment,^46,53^ but here we enhance detection of pre-mRNA sequences generally by using global splicing inhibition. We expect that most pre-mRNA in pladB-treated cells remain unspliced rather than reaching later stages of splicing, as accumulating pre-mRNA are expected to outnumber spliceosome components.^54^ It is possible that splicing inhibition could alter the structures of pre-mRNA compared to untreated cells, as accumulating pre-mRNA may be unbound or interacting with the nuclear pore complex,^55^ nuclear exosome,^56^ or nonsense mediated decay machinery.^57^ Nevertheless, for the cases with sufficient sequencing coverage from experiments with untreated cells, we found similar high confidence stems identified with and without splicing inhibition. In future work, it would be interesting to directly observe long-range RNA base-pairing and higher-order RNA structures in introns with approaches like PARIS^58^ and KARR-seq^59^ in pladB-treated cells.

With DMS data for introns transcriptome-wide, we discerned structural patterns that distinguish introns from coding regions. For instance, with higher Gini coefficients and longer stems, stable structures were enriched in introns compared to coding regions. While it is possible that active translation may unwind structures in coding regions, our computational secondary structure predictions suggest that intron sequences have the capacity to form more stable structures regardless of their translation status. It is for instance possible that introns’ intramolecular structures are necessary to avoid spurious interactions between intron RNA and other nucleic acids in the crowded nuclear environment, preventing gene expression dysregulation. Additionally, while there is evidence across species for depletion of double stranded RNA in coding RNA to avoid activating cytosolic antiviral cellular responses,^60^ this selection would not be present for introns in the nucleus. Long stems may also be depleted from coding mRNA because cytosolic mRNAs with stable stems may accumulate stalled ribosomes, becoming subjected to decay pathways like no-go decay.^31,61^ Finally, without the evolutionary constraints of adhering to a viable coding sequence, introns may be free to form extended structures that can play roles in regulating a host of nuclear processes.

With structural motifs enriched in *S. cerevisiae* introns compared to coding regions and random control sequences, we hypothesized that these introns could harbor functional structures. While scans for covariation have provided evidence for functional structures in a handful of *S. cerevisiae* introns (RPS9A,^7^ RPS9B, RPS13,^23^ RPL7B,^23^ and RPL7A), this analysis may miss functional structures due to the limited power of existing intron sequence alignments,^62^ with high sequence variation and large-scale deletions often present between aligned intron sequences. Indeed, tools like R-scape can be limited by the quality of sequence alignments, the conservation of positions in alignments, and the number of sequences in alignments.^42,62^ By assaying intron variants and rescue sequences in high-throughput with VARS-seq, we identified additional functional structures in introns. While prior intron variant libraries have been designed to assess the effects of sequence motifs on splicing patterns,^63,64^ libraries assessing the role for structured elements in introns have been limited. For interrogating the role of structure, it was critical to include both variant and rescue sequences where possible and to test a library of multiple sequences and barcodes rather than single sequence variants due to variation between constructs. With VARS-seq, we found that some introns included domains that could influence gene expression despite being distal from splice sites in the intron’s secondary structure. These examples challenge the notion that introns are largely nonfunctional junk DNA sequences, adding to the known complexity of potential functional roles for introns.^9,10^ We noted that absolute effect sizes obtained from VARS-seq are likely impacted by length biases from the assay, and we encourage evaluating individual strains with orthogonal assays to validate effects for specific stems of interest, which we carried out for stem sets in RPS9A and RPL36B.

Together, our structural, computational, and functional experiments point to structural patterns across introns that impact gene expression. In one common pattern, we found an enrichment for structures around cryptic splice sites and a depletion of structure around canonical splice sites, potentially indicating a role for structures in encouraging use of the canonical sites. Another pattern that emerged was the formation of zipper stems co-localizing the 5’ splice site and the branch point. Introns included highly stable zipper stem structures *in vivo*, and these structures were additionally supported by multi-dimensional chemical mapping (M2-seq) *in vitro*. These zipper stems could enable efficient splicing by reducing the physical distance between the 5’ splice site and branch point^65^ or by enhancing specific interaction with the spliceosome, with intron helical density seen interacting directly with the E and pre-A spliceosome complexes.^25,36^

To interrogate the mechanisms by which these and other structured elements regulate gene expression, we will need customized experiments for individual introns. Structures located close to splice sites could have functional roles in recruiting the spliceosome, while structures extruded away from splice sites could play a role in recruiting other nuclear factors. Intron secondary structures can regulate splicing by sequestering alternative splice sites,^66,67^ occluding exonic splicing enhancers,^68^ co-localizing selected splice sites,^69^ facilitating co-transcriptional splicing,^70^ or mediating protein interactions that influence splicing patterns.^71^ Additionally, these intronic structures could have regulatory functions in pathways orthogonal to splicing, much like the snoRNAs encoded in *S. cerevisiae* introns.^41^ In fact, RNA secondary structures in introns associate with RNA-binding proteins involved in varied processes including transcription, tRNA and rRNA processing, ribosome biogenesis and assembly, and metabolic processes.^72,73^ Furthermore, secondary structures in *S. cerevisiae* introns are poised to regulate gene expression in auto-regulatory circuits as seen previously in the cases of RPS14B^21^ or RPS9A^7^, with structural elements within a gene’s introns binding its protein products and thereby downregulating subsequent splicing and gene expression. Intron structures may additionally influence numerous pre-mRNA decay pathways, including nuclear retention followed by decay by the nuclear exosome, nonsense-mediated decay (NMD), and NMD-independent decay by cytoplasmic exonucleases.^31,32,55,56,74^ Finally, structures can play a functional role in regulating adaptation to starvation by influencing the accumulation of linearized introns under nutrient depletion,^5,6^ and it will be interesting to probe structures in saturated-growth conditions or other stress conditions.

Our work identifies a set of intron structures with properties distinct from those in coding regions, support for *in vivo* formation from DMS-MaPseq, signals in other related yeast species, and in some cases, covarying residues. Furthermore, we identify structured intervals of introns that modulate gene expression, impacting levels of retained introns in cells. These functional experiments provide candidates for further mechanistic characterization and provide a glimpse into the broad regulatory potential for intron sequences beyond splice sites. The widespread presence of structured elements in *S. cerevisiae* introns raises the possibility that similar motifs and stable secondary structures play a role in introns in higher-order eukaryotes, perhaps forming regulatory elements in human pre-mRNA.

## Online Methods

### Strains, media, and growth conditions for DMS probing

The strain OHY001 was constructed from the background strain JRY8012,^75^ which includes three deletions of ABC transporter genes (*prd5::kan^r^, snq2::kan^r^, yor1::kan^r^*) to reduce drug efflux. OHY001 was generated via CRISPR editing of JRY8012 to mutate portions of HSH155 HEAT repeat domains 15-16 to match the sequence found in human SF3B1 (see sequence in Table S5).

Strains were grown at 30 °C on YPD plates and in YPD liquid medium. Single colonies of OHY001 were used to inoculate overnight cultures, diluted to OD600 0.1, and grown to OD600 0.5-0.6. Biological replicates were obtained from distinct single colonies.

### Splicing inhibition and DMS treatment

We carried out splicing inhibition and DMS treatment for two biological replicates, splicing inhibition only for a no-modification control, and DMS treatment only for the control without splicing inhibition. We grew 15 mL of culture to OD600 ∼0.5 for replicate 1, 60 mL for replicate 2, 15 mL for the no-modification control, 20 mL for the condition without pladB treatment, 20 mL for *in vitro* replicate 1, and 20 mL for *in vitro* replicate 2. We treated cultures with 5 μM Pladeinolide B (pladB, Cayman Chemicals) using 50 μL of 1 mM pladB for every 10 mL of culture, and we incubated cultures at 30 C for 1 hour with shaking. The condition without splicing inhibition was treated with an equal volume of DMSO.

For *in vivo* DMS modification, we treated cultures with 3% DMS or an equivalent volume of H_2_O for the no-modification control. Treated cells were incubated with occasional stirring in a water bath at 30 °C for 5 minutes, and the reaction was quenched by adding 20 mL stop solution (30% 2-mercaptoethanol, 50% isoamyl alcohol) for every 10 mL of culture. The culture was mixed multiple times by inversion and transferred quickly to ice. Cultures were spun down for 3 minutes at 1500g at 4 °C, and washed first with 5 mL wash solution (30% 2-mercaptoethanol) for every 10 mL of culture. A second wash was performed with 3 mL YPD per 10 mL culture. RNA was extracted using the YeaStar RNA Kit (Zymo Research), using 7.5 μL of Zymolase for every 2.5 mL starting cell culture and shortening the Zymolase incubation to 15 minutes at 30 °C to reduce RNA fragmentation.

For *in vitro* DMS modification, we followed a protocol similar to the one used in Rouskin, et al. (2014).^76^ We first obtained RNA from cultures after splicing inhibition, pelleting cells by spinning at 1500g for 3 minutes, resuspending in YPD, and extracting RNA with the YeaStar RNA Kit (Zymo Research) using 1 column for every 5 mL starting culture. We then re-folded RNA *in vitro*, first denaturing 200 μg of RNA at 95 °C for 2 minutes, cooling RNA on ice for 2 minutes, and then folding RNA at 30 °C for 30 minutes in 10 mM Tris HCl pH 8.0, 100 mM NaCl, and 6 mM MgCl_2_. We treated RNA with 3% DMS at 30 °C for 5 minutes, and we quenched with 25% 2-mercaptoethanol to stop the reaction. RNA was purified by ethanol precipitation and eluted in 20 μL RNAse-free H_2_O.

### RT-PCR for verifying splicing inhibition

As initial verification of splicing inhibition by pladB, we used RT-PCR to compare unmodified RNA extracted after 1 hour of either 5 μM pladB or DMSO treatment. We first Dnase treated the +pladB and -pladB samples by mixing 1 μL of TURBO Dnase (Thermo Fisher), 20 ng of RNA, and Rnase-free H_2_O for a 20 μL reaction volume. We incubated the reactions at 37 °C for 1 hour, added 2 μL of 50 mM EDTA, incubated the reactions at 65 °C for 10 minutes, and placed them on ice. We then carried out reverse transcription with the iScript Reverse Transcription Supermix (Bio-Rad), using 10 μL of each RNA sample for a reaction that included the reverse transcriptase, and the remaining 10 μL for a control without this enzyme. 4 μL of the 5X RT Supermix was mixed with 10 μL of RNA sample and water to a reaction volume of 20 μL, and the reaction was incubated in a thermocycler for 5 minutes at 25 °C, 40 minutes at 46 °C, and 1 minute at 95 °C. The samples were purified with Oligo Clean and Concentrator columns (Zymo Research) and eluted in 15 μL. PCR was then performed using 2 μL of cDNA template, 25 μL of the 2X NEBNext Ultra II Q5 Master Mix (NEB), 18 μL of H_2_O, and 2.5 μL of 10 μM primers (RR063 and RR064 for RPL36B, and RR067 and RR070 for MATa1; Table S5). The reactions were denatured 98 °C for 30 seconds, followed by 30 cycles of 98 °C for 10 seconds, 56 °C for 30 seconds, and 72 °C for 30 seconds, and ending with a final extension period at 72 °C for 5 minutes. PCR products were visualized on a 2% agarose gel with ethidium bromide.

### DMS-MaPseq sequencing library preparation

For preparation of DMS-treated RNA for sequencing, we first depleted the extracted RNA of rRNA using Rnase H to deplete rRNA with complementary oligos. We concentrated extracted RNA for each condition using RNA Clean and Concentrator-5 columns (Zymo Research), yielding 47.4 μg RNA for *in vivo* replicate 1, 200 μg RNA for *in vivo* replicate 2, 47.2 μg RNA for the no-modification condition, 53.2 μg RNA for the condition without splicing inhibition, 58.0 μg RNA for *in vitro* replicate 1, and 46.1 μg RNA for *in vitro* replicate 2. We pooled 108 50-mer oligos (RR-rRNAdep-1-108; Table S5) that tiled the 5S, 5.8S, 18S, and 25S rRNA in *S. cerevisiae*, and we concentrated these oligos with Oligo Clean and Concentrator column. Up to 40 μg of total RNA was included in each rRNA depletion reaction, with multiple reactions as needed for each sample. First, a 15 μL annealing reaction was prepared with total RNA, an equal mass of rRNA depletion oligos, and 3 μL of 5X hybridization buffer (500 mM Tris-HCl pH 7.5 and 1 M NaCl). For the annealing reaction, the reaction mix was heated to 95 °C for 2 minutes and the temperature was ramped down to 45 °C by 0.1 °C/sec with a thermocycler. 7.5 μL of Hybridase thermostable Rnase H (Lucigen) with 2.5 μL of 10X digestion buffer (500 mM Tris-HCl pH 7.5, 1M NaCl, 100 mM MgCl_2_) was preheated to 45 °C. We combed the annealing mix and the Rnase H mix and incubated at 45 °C for 30 minutes. For each reaction, we then used an RNA Clean and Concentrator-5 column with the size-selection protocol to exclude RNA below a size cutoff of 200 nucleotides, cleaning up the shorter oligos used for rRNA depletion and removing small RNA from the total RNA sample. We then Dnase treated each reaction by combining each sample with 10 μL of TURBO Dnase in a 167 μL reaction. We purified the reaction using an RNA Clean and Concentrator-5 column, eluting in 9 μL of Rnase-free H_2_O. After rRNA depletion and size-exclusion of small RNA, we obtained 1.76 μg RNA for *in vivo* replicate 1, 4.58 μg RNA for *in vivo* replicate 2, 1.64 μg RNA for the no-modification control, 1.57 μg RNA for the condition without splicing inhibition, 2.74 μg RNA for *in vitro* replicate 1, and 1.81 μg RNA for *in vitro* replicate 2. We split the RNA from replicate 2 into 4 samples with 1.15 μg RNA each to scale up the enzyme amounts used for the following library preparation steps.

We fragmented each reaction by combining 1 μL of 10X RNA Fragmentation Reagent (Ambion) with 9 μL of RNA sample, incubating for 8 minutes at 70 °C. We added 1 μL of the Stop Solution (Ambion), mixed thoroughly, and placed immediately on ice. Each reaction was cleaned with an RNA Clean and Concentrator-5 column, eluting in 11 μL of Rnase-free H_2_O. We removed the 3’ phosphate groups left by fragmentation by adding 1.5 μL rSAP (NEB), 1.5 μL 10X Cutsmart buffer (NEB), and 1 μL SUPERase inhibitor (Thermo Fisher), and incubating the 15 μL reaction at 37 °C for 1 hour. The reaction was stopped at 65 °C for 4 minutes.

We next ligated a universal cloning linker to the RNA to serve as a handle for reverse transcription. To prepare linker for this reaction, we first phosphorylated 1 nmol of the DNA universal cloning linker with a 3’ amino blocking group (oligo RR118; Table S5) by adding 3 μL of T4 PNK (NEB), 15 μL of 10X T4 PNK buffer (NEB), and 15 μL of 10 mM ATP in a 150 μL reaction; incubating at 37 °C for 30 minutes; and inactivating at 65 °C for 20 minutes. We then purified the reaction with an Oligo Clean and Concentrator column. Next, we adenylated the linker by adding 20 μL of Mth RNA Ligase (NEB), 20 μL of 5’ DNA adenylation reaction buffer (NEB), and 20 μL of 1 mM ATP to the sample in a 200 μL reaction. We incubated the reaction at 65 °C for 1 hour and inactivated the reaction at 85 °C for 4 minutes. We then purified the reaction with an Oligo Clean and Concentrator column and measured the final adenylated linker concentration. We added adenylated linker in 2-fold molar excess to each RNA sample, obtaining the molarity of the RNA samples by estimating the library to contain RNA of average length 150. 3 μL of T4 RNA ligase 2 truncated KQ (NEB), 3.5 μL 10X NEBuffer 2 (NEB), 1.75 μL 100 mM DTT, and 10 μL 50% PEG-8000 were added to adenylated linker and RNA sample to make a 35 μL reaction. The reaction was incubated at 25 °C for 2 hours in a thermocycler.

RNA ligated to the DNA linker was purified using an RNA Clean and Concentrator-5 column, eluting in 15 μL of Rnase-free H_2_O. The excess DNA linker was then degraded by adding 1 μL of 5’ Deadenylase (NEB), 1 μL of RecJf (NEB), 1 μL of SUPERase inhibitor, and 2 μL of NEBuffer 2 in a 20 μL reaction. The mixture was incubated at 30 °C for 1 hour and purified with an RNA Clean and Concentrator-5 column.

We then proceeded to reverse transcription (RT) with mutational readthrough and denaturing PAGE gel size selection of the resulting cDNA library. For the RT primer, we used oligo RR114 (Table S5) which included a sequence complementary to the universal cloning linker, a 5’ phosphate modification that would allow for circularization after RT, a 10-nucleotide randomized UMI sequence, and sequences complementary to Illumina sequencing primers that are separated by a spacer to allow for PCR amplification of the final library. We added 2 μL of 5X TGIRT buffer (250 mM Tris-HCl pH 8.0, 375 mM KCl, 15 mM MgCl_2_), 0.5 μL of 2 μM RT primer, and 6 μL of the RNA sample. The reaction was denatured at 80 °C for 2 minutes and left at room temperature for 5 minutes. 0.5 μL TGIRT enzyme (InGex), 0.5 μL SUPERase inhibitor, 0.5 μL 100 mM DTT, and 1 μL 10 mM dNTPs were added to the reaction. The reaction was incubated at 57 °C for 1.5 hours in a thermocycler for reverse transcription. RNA was then degraded by adding 5 μL of 0.4 M NaOH for 3 minutes at 90 °C, and the reactions were neutralized by adding 5 μL of an acid quench mix (from a stock solution of 2 mL of 5 M NaCl, 2 mL of 2 M HCl, and 3 mL of 3 M NaOAc). The reaction was purified with an Oligo Clean and Concentrator column, eluting in 7.5 μL of Rnase-free H_2_O. cDNA was then purified with a denaturing PAGE (dPAGE) gel to remove excess RT primer. dPAGE gels were cast with 10% 29:1 bis-acrylamide, 7 M urea, and 1X TBE using 150 μL of 10% APS and 15 μL of TEMED. After pre-running the gel at 17 W for 1 hour, we loaded the gel with samples that had been denatured at 90 °C for 3 minutes in 50% formamide and a 1X TBE loading buffer with xylene cyanol and bromophenol blue markers. Gels were run for 40 minutes at 17W and stained for 20 minutes with 5 μL of 10,000X Sybr Gold (Invitrogen) in 50 mL of 1X TBE. We cut cDNA in the 200 to 400 nucleotide size range and eluted cDNA using the ZR small-RNA PAGE Recovery Kit (Zymo Research), following the manufacturer’s protocol except for the changes noted as follows. We centrifuged the macerated gel slice for 15 minutes at max speed for the first column step, centrifuged at 5000 g for 5 minutes for the second column, and eluted the final sample in 7.5 μL of Rnase-free H_2_O warmed to 65 °C.

The purified and size-selected cDNA was circularized and then amplified for sequencing. For circularization, 0.5 μL of CircLigase ssDNA Ligase (Lucigen), 2 μL of 10X CircLigase buffer (Lucigen), 1 μL of 1 mM ATP, and 1 μL of 50 mM MnCl_2_ were added to the purified cDNA, and the reaction volume was adjusted to 20 μL. The reaction was left to incubate overnight at 60 °C in a thermocycler with a heated lid, and then the reaction was stopped at 80 °C for 10 minutes. The circularized cDNA was purified with an Oligo Clean and Concentrator column and eluted in 7.5 μL of Rnase-free H_2_O. Samples from replicate 2 that were processed in parallel from RNA fragmentation to circularization were combined in this column purification step. Residual RT primer was removed by degrading linear DNA. First, we added 1 μL of RecJf and 1 μL of NEBuffer2 to the sample and adjusted with H_2_O to make a 20 μL reaction, incubating at 37 °C for 10 minutes. We then added 2 μL each of ExoCIP A and ExoCIP B (NEB) and incubated for another 10 minutes at 37 °C, followed by 1 minute of heat inactivation at 80 °C. The sample was purified with an Oligo Clean and Concentrator column, with elution volume adjusted to 10 μL.

We carried out 9 cycles of indexing PCR to add i5 and i7 index sequences to the sequencing library, making a 50 μL reaction with 25 μL of the 2X NEBNext Ultra II Q5 Master Mix (NEB), 10 μL of cDNA, and 2.5 μL of 10 μM primers including different index sequences for each sample (i5_4 and i7_4 for replicate 1, i5_3 and i7_3 for replicate 2, i5_5 and i7_5 for the no-modification control, and i5_2 and i7_2 for the -pladB control; Table S5). The reactions were denatured at 98 °C for 30 seconds, followed by 9 cycles of denaturing at 98 °C for 10 seconds, annealing at 70 °C for 30 seconds, and extending 72 °C for 30 seconds, and ending with a final extension period at 72 °C for 5 minutes. The reaction mix was purified with the DNA Clean and Concentrator-5 kit (Zymo Research), with elution in 10 μL of H_2_O. To determine the number of amplification cycles required for obtaining sufficient library concentration for sequencing, we quantified the library size through qPCR using the P5 and P7 primer sequences (Table S5), using the iTaq Universal SYBR Green Supermix (Bio-Rad). Based on this qPCR quantification, we carried out 2 more cycles of PCR for *in vivo* replicates 1 and 2 and the no-modification control, 5 more cycles for the -pladB control, 9 more cycles for *in vitro* replicate 1, and 3 more cycles for *in vitro* replicate 2. For this final PCR, we again used 25 μL of the Q5 Master Mix and 2.5 μL of 10 μM primers (P5 and P7 primers), using the same cycling parameters as the first indexing PCR. We purified the final library with two rounds of bead purification with size selection to remove remaining excess RT primer. We used RNACleanXP beads (Beckman Coulter), mixing 42.5 μL beads with the 50 μL PCR reaction (0.85 reaction volume ratio for size selection), shaking for 10 minutes and then separating beads from supernatant using a magnetic post for 10 minutes. We then washed the beads twice with 200 μL 70% EtOH, dried the beads, and eluted in H_2_O after shaking the beads with H_2_O for 10 minutes. For the first round of bead-based size selection we eluted in 50 μL of H_2_O, and for the second round, we eluted in 20 μL of H_2_O. The sequencing libraries were quantified with a Qubit high sensitivity dsDNA Assay Kit (Invitrogen) and Bioanalyzer HS DNA assay, and sequenced with a 10% PhiX library spike-in. The dsDNA libraries for *in vivo* replicate 1, the no-modification control, and the -pladB control were sequenced across three Illumina HiSeq lanes with paired-end reads of length 150. The dsDNA library for *in vivo* replicate 2 and the *in vitro* replicates were sequenced with NovaSeq S4 partial lanes, also with paired-end reads of length 150.

### DMS-MaPseq sequencing data analysis

We obtained 557 million reads for *in vivo* replicate 1, 1.30 billion reads for *in vivo* replicate 2, 375 million reads for the no-modification control, 169 million reads for the control without splicing inhibition, 335 million reads for *in vitro* replicate 1, and 174 million reads for *in vitro* replicate 2. We used UMI-tools^77^ to extract UMI tags from reads by matching to the expected pattern from our RT primers, and we used cutadapt to trim low quality reads (Q-score cutoff 20) and remove adapter sequences including indexing primer sequences and the cloning linker. We then aligned sequencing reads to sequence sets of interest, including rRNA sequences, introns, complete pre-mRNA ORFs for genes containing introns, coding mRNA sequences for genes containing introns, decoy intron sequences (see below), and sequences for structured controls (the ASH1 and HAC1 mRNA sequences). *S. cerevisiae* intron annotations including genomic coordinates and branch point positions were obtained from Talkish, et al. 2019.^78^ Coding ORF annotations were obtained from the Saccharomyces Genome Database for the S288C reference genome, and coding sequences corresponding to introns were identified for all cases except the two introns in snoRNAs (SNR17A and SNR17B). Paired-end alignment was performed with Bowtie2^79^ using the alignment parameters used in ShapeMapper 2^80^: --local –sensitive-local –maxins=800 –ignore-quals –no-unal –mp 3,1 –rdg 5,1 –rfg 5,1 – dpad 30. To obtain mutational frequencies and normalized reactivities for both replicates combined, we additionally used alignment files merged with samtools^81^. Alignments were sorted and indexed using samtools. Reads with matching mapped positions and UMI tags were then deduplicated using the UMI-tools^77^ dedup function with default parameters for paired-end reads.

Mutational frequencies, coverage values, and normalized reactivities were obtained by processing alignment files using RNAframework^82^ executables. For each library and reference sequence set, we first ran rf-count with the flag -m to compute mutation counts, using all other default parameters. We obtained coverage statistics for each sequence with rf-rctools stats, and we obtained per-position mutation counts and coverage statistics with rf-rctools view. We obtained reactivity values with rf-norm, using Rouskin scoring with 90% Winsorizing for normalization, reactive bases A and C, and dynamic resizing of the normalization window to account for A/C frequencies (flags: -sm 2 -nm 2 -ow -rb AC -dw).

### DMS-MaPseq data quality assessment

For each construct, per-base coverage was computed as the number of paired-end reads mapped to the construct obtained from rf-rctools stats multiplied by the total read length and divided by the length of the construct. To compare the retained intron fraction across all introns before and after pladB treatment, these coverage values were obtained for all introns and all coding regions for genes containing introns in the -pladB and +pladB conditions. The retained intron fraction was the ratio of these values. We removed introns in GCR1 from further consideration as the gene includes multiple distal alternative 5’ splice sites.^83^ We combined data from two introns in SRC1 from nearby alternative 5’ splice sites differing by 4 nucleotides. We additionally consolidated data from two-intron genes with multiple annotated isoforms (in RPL7B, VMA9, DYN2, and SUS1) where annotated introns included both single individual introns and the longer intron representing the skipped isoform.^84^ We used annotations for the ribosomal protein-coding genes from Hooks, et al. 2014.^51^ We excluded eight snoRNA-containing introns from further analysis,^41^ as the majority of these constructs’ reactivities represented the excised snoRNA structure rather than the complete intron structure.

We next found the Pearson correlation between intron reactivities between replicate 1 and replicate 2 as it related to the average coverage between these replicates. A linear fit for log coverage versus replicate correlation for each intron indicated that the replicate correlation was best approximated as 0.2 log(coverage) – 1.03 for the DMS-MaPseq replicates. Based on this relationship, to reach 0.5 Pearson correlation between replicates, we used the coverage cutoff of 1971 averaged between replicate 1 and replicate 2, or 3942 when combining data from the two replicates.

Per-base mutational frequencies were computed starting with the per-position mutation counts and coverage values from rf-rctools view. For each nucleotide identity, we found the average mutational frequency for each intron by averaging mutation rates across all positions in the intron with this nucleotide identity. We then found the average mutation frequencies for each nucleotide across all introns meeting the coverage threshold of 3942.

To obtain the correlation between Zubradt, et al. (2017)^27^ rRNA mutational frequencies and our data, we obtained this paper’s DMS-MaPseq reads for *S. cerevisiae* with TGIRT reverse transcriptase (replicate 1 accession number SRX1959209, run number SRR3929621). We aligned these reads to the 18S and 25S yeast rRNA sequences with Tophat v2.1.0^85^ using alignment parameters stated in Zubradt, et al. (2017)^27^: -N 5 –read-gap-length 7 –read-edit-dist 7 –max-insertion-length 5 –max-deletion-length 5 -g 3. We obtained mutational frequencies with rf-count as described above. We compared mutational frequency values for all A/C positions where per-position coverage values met our coverage threshold, finding the Pearson correlation for these values between our data and those in Zubradt, et al. (2017)^27^.

We evaluated whether reactivity values could distinguish surface accessible, unpaired residues from base-paired residues across the 18S and 25S rRNA. To identify surface accessible residues in rRNA, we followed a similar protocol to Rouskin, et al. (2014)^76^. We used the high-resolution *S. cerevisiae* ribosome structure with PDB ID 4V88,^86^ finding solvent accessibility for rRNA residues in the complete context of the ribosome. We computed solvent-accessible surface area (SASA) values for the rRNAs’ N1 atoms on A residues and N3 atoms on C residues in PyMOL, approximating DMS as a sphere with solvent_radius 3 and with dot_solvent and dot_density parameters set to 1. We additionally used this ribosome structure with DSSR^87^ to find the secondary structure of the 18S and 25S rRNA sequences. We found the ROC curve for distinguishing unpaired and solvent-accessible rRNA residues from Watson-Crick base-paired positions, using a SASA cutoff of 2 to determine solvent accessibility.

We assessed whether reactivity profiles for known stems in the HAC1 and ASH1 mRNA sequences accurately reflected their secondary structures. Secondary structures for these stems were obtained from Zubradt, et al. (2017)^27^.

### DMS-guided structure prediction and validation

For introns meeting coverage cutoffs as discussed above, we performed secondary structure prediction by RNAstructure^33^ guided by DMS reactivity with 1000 bootstrapping iterations without pseudoknot predictions, using the package Biers^47^ with default parameters for RNAstructure. Minimum free energy structures and base-pair confidence matrices from bootstrapping were obtained for each construct. Structures were visualized with VARNA, using Biers to display DMS reactivity values and per-helix helix confidence scores from bootstrapping.

To assess DMS-guided structure prediction and the helix confidence scores, we performed structure prediction for a set of controls with known secondary structures. These included rRNAs, snRNAs, tRNAs, and mRNA segments, all of which were included in our transcriptome-wide DMS-MaPseq dataset at high coverage. The ground truth secondary structures for the 5S, 5.8S, and 18S rRNA was obtained from a high-resolution X-ray crystallography structure of the eukaryotic ribosome (PDB ID: 4V88)^86^, and DSSR^87^ was used to obtain the secondary structure from these 3D models. The ground truth U5 snRNA secondary structure was obtained from Nguyen, et. Al. (2016)^88^, and the U1 snRNA secondary structure was obtained from Li, et. Al. (2017)^89^. Four tRNA ground truth structures were obtained from Rfam-derived secondary structures.^90^ Finally, secondary structures for mRNA segments in HAC1, ASH1, RPS28B, and SFT2 were obtained from Zubradt, et al. (2017)^27^. We did not include RNAs with known pseudoknots in the control set (for instance, the Rnase P RNA^91^), as our structure prediction approach would not be able to recover pseudoknots. Additionally, we excluded control cases where the RNA was expected to take on multiple distinct structural conformations, such as the U2 snRNA.^92^ For DMS-guided structure prediction, 1000 bootstrapping iterations were performed for all cases except the 18S rRNA, for which we performed 100 bootstrapping iterations. Predicted stems were called using varying helix confidence estimate cutoffs. As a point of comparison, structure predictions were also made using ViennaRNA 2.0^50^ to generate predictions unguided by DMS data. All stems in control structures and predictions were evaluated for true positive, false positive, and false negative stem predictions. Positive stem predictions were defined as cases in which a stem in the predicted structure included at least five base-pairs and the helix confidence estimate threshold was met. False positive stem predictions were those where less than 50% of the predicted stem’s base-pairs were included in the ground truth structure. False negative stem predictions were stems of length at least five in the native structure that either had fewer than 50% of their base-pairs in the predicted structure, or did not meet the helix confidence estimate threshold in the predicted structure.

We additionally explored other methods for structure prediction using DMS data. First, using the Arnie package (https://github.com/DasLab/arnie), we ran ShapeKnots^93^ (which allows for pseudoknot prediction) on all 161 introns with sufficient coverage, guiding predictions with DMS data in 100 bootstrapping iterations with default parameters. As a control, we ran structure predictions with ShapeKnots for the Rnase P RNA with 100 bootstrapping iterations, comparing to the secondary structure obtained by using DSSR on the high-resolution cryo-EM structure of the yeast RNase P (PDB ID: 6AGB)^91^. We additionally made predictions using DREEM^94^ for the following eight intron regions that included predicted structures and exhibited high coverage from DMS-MaPseq, with intervals numbered starting at the beginning of each intron: RPL28 75-300, RPL7A 120-340 (first intron), ECM33 35-270, RPL26B 300-400, RPS13 140-212, RPL25 148-255, RPS9B 147-266, and RPL30 25-165. We ran DREEM using treating sequencing reads as single-stranded, and used default parameters otherwise.

### Two-dimensional chemical mapping (M2-seq)

We carried out two-dimensional chemical probing to assess the formation of base pairs in secondary structures predicted from DMS-MaPseq. DNA sequences for the introns in QCR9, RPL36B, RPS11A, RPL37A, and RPS7B were obtained as gene fragments from Twist Biosciences (Table S5). Each gene fragment included the full intron sequence, a T7 promoter sequence, reference hairpins for normalizing structure probing data, and sequences complementary to universal RT and PCR primers used for library preparation. To generate a pool of DNA variants through error-prone PCR, we first assembled a reaction mix with the following for each intron: 10 μL of 2 ng/μL template, 10 μL of 100 mM Tris pH 8.3, 2.5 μL of 2 M KCl, 3.5 μL of 200 mM MgCl_2_, 4 μL of 25 mM dTTP, 4 μL of 25 mM dCTP, 4 μL of 5 mM dATP, 4 μL of 5 mM dGTP, 2 μL of 25 mM MnCl_2_, 1 μL of Taq polymerase (Thermo Fisher), 2 μL of 100 mM primers (RR1 and RR107; Table S5), and 51 oμL of H_2_O. We ran the reaction on a thermocycler for an initial 98 C denaturation, and then 24 cycles of 94 °C denaturation for 30 seconds, 64 °C annealing for 60 seconds, and 72 °C elongation for 3 minutes, followed by a final elongation at 72 °C for 10 minutes. Construct sizes were verified on a 2% agarose gel, and samples were purified as described above using RNACleanXP beads (Beckman Coulter) with a 1.8 Ampure bead to sample volume ratio. We then proceeded with *in vitro* transcription of these RNA fragments by assembling the following reaction mix: 5 μL of 1X T7 RNA polymerase (NEB), 2 μL of 1 M DTT, 6 μL of 25 mM NTPs, 5 μL of 40% PEG-8000, 5 μL of T7 transcription buffer (NEB), template DNA, and Rnase-free H_2_O added to a final reaction volume of 50 μL. Reactions were incubated at 37 °C overnight. Samples were Dnase treated by adding 1 μL of TURBO Dnase (Thermo Fisher) to the reaction and incubating at 37 °C for 30 minutes. RNA was purified with RNACleanXP beads using a 70:30 ratio of beads to 40% PEG-8000 as the beads mixture, and using a 1.8 bead mixture to sample volume ratio.

We proceeded with structure probing, reverse transcription, and library preparation for these RNA pools. We prepared 3 μL of 12.5 pmol RNA for each intron for DMS treatment and a no-modification control. We denatured the RNA by unfolding at 95 °C for 2 minutes and left RNA on ice for 1 minute. RNA was folded by adding 5 μL of 5x folding buffer (1.5 M sodium cacodylate pH 7.0. 50 mM MgCl_2_) and 14.5 μL of Rnase-free H_2_O, and incubating for 30 minutes at 37 °C. We modified RNA by adding 2.5 μL of 15% DMS (DMS condition) or 100% EtOH (no-modification control), and modifying at 37 °C for 6 minutes. The reaction was quenched by adding 25 μL of 2-mercaptoethanol and purified by Ampure bead purification as described above, eluting in 7 μL of Rnase-free H_2_O. We reverse-transcribed the modified RNA with mutational read-through using TGIRT. We assembled a TGIRT reverse transcription master-mix by combining 2.4 μL 5X TGIRT buffer (see above), 1.2 μL of 10 mM dNTPs, 0.6 μL of 100 mM DTT, 0.5 μL TGIRT enzyme (InGex), and 1.82 μL of H_2_O. For reverse transcription, we used FAM-labeled primers that included a distinct index sequence for each construct and condition. We mixed 0.93 μL of RT primer (RTB primers in Table S5), 4.6 μL of RNA, and 6.52 μL of the reverse transcription master mix. We incubated the reaction mix at room temperature for 5 minutes and then at 57 °C for 3 hours. The reaction was stopped by adding 5 μL of 0.4 M NaOH and incubating at 90 °C for 3 minutes to degrade RNA, and the solution was neutralized by adding 2.2 μL of an acid quench mix (see above). cDNA was purified using RNACleanXP beads and eluted in 10 μL H_2_O, using the protocol for RNA bead purification as described above. The cDNA library was then amplified, adding Illumina adapters for sequencing. We mixed 5 μL of 5X Phusion HF buffer (NEB), 0.5 μL of 10 mM dNTPs, 0.5 μL of Phusion high-fidelity DNA polymerase (NEB), 2.5 μL of cDNA, and 1 μL of 100 μM primers (MaP forward and reverse primers; Table S5) in a reaction volume of 25 μL. The PCR reaction used the following thermocycler conditions: denaturation at 98 °C for 30 seconds, 20 cycles of 98 °C for 10 seconds, 65 °C for 30 seconds, and 72 °C for 30 seconds, and a final elongation at 72 °C for 10 minutes. Amplicon sizes were verified on a 2% agarose gel with ethidium bromide staining. We collected library pools for the sequencing run, combining 0.57 fmol for each DMS treatment case and 0.18 fmol for each no-modification control, and adding a 25% PhiX spike-in. The resulting libraries were sequenced in two partial sequencing runs using Illumina MiSeq v3 600-cycle reagent kits, providing paired-end reads of length 300.

M2-seq sequencing reads were processed using the M2seq package.^47^ First, barcodes for the different introns and conditions were demultiplexed. We obtained at least 500,000 reads for each intron’s DMS treatment condition. The ShapeMapper 1.2^80^ software was used to align reads to each reference sequence using Bowtie2 and compute mutation rates. The output from ShapeMapper was processed by the simple_to_rdat.py script to obtain 2D reactivity values and raw counts in an RDAT file. These RDAT files were obtained for both DMS and no-modification conditions, and processed with the Biers package^47^ to generate Z-score plots, base-pairing probability matrices from bootstrapping, and predicted secondary structures. These secondary structure predictions were made using RNAstructure’s Fold executable^33^ with 500 bootstrapping iterations, guided predictions with both 1D and 2D DMS reactivity data. Structures were visualized using VARNA, using Biers to display reactivity profiles and helix confidence estimates from bootstrapping for all stems.

### Assessing proposed functional structures with DMS data

Using our DMS data, we assessed proposed structures from prior functional experiments and analysis of sequence alignments. For each evaluated structure, we displayed the proposed structure with reactivity values and helix confidence estimates using Biers.^95^ Structure from Hooks, et al. 2016^37^ were obtained by request. For structures with covarying residues, DMS-guided structure predictions were displayed when they agreed with the proposed covarying base-pairs (RPS9A, RPS9B, and RPL7A), and structures from CaCoFold^96^ were displayed otherwise (RPS13).

To expand the set of covarying residues identified across introns, we used the protocol and software developed by Gao, et al. 2020^23^ with thresholds as follows. We obtained all complete and chromosome-level assemblies of genomes in the Ascomycota phylum from NCBI GenBank^97^ (accession date: August 17, 2021), yielding 107 genome sequences. For each *S. cerevisiae* intron, we generated multiple-sequence alignments using the scripts from Gao, et al. 2020^23^ in flanked mode with a noncoding threshold of 50. We then used R-scape^42^ to predict covarying residues with a 0.05 E-value threshold, and we generated structures with CaCoFold.^96^ We identified all introns for which at least 1 significantly covarying pair was identified between two non-consecutive residues in a stem that contained at least 3 base-pairs in the CaCoFold structure. We evaluated the presence of these covarying residues in minimum free-energy secondary structures predicted from ViennaRNA 2.0^50^. The set of snoRNA-containing introns was obtained from the *S. cerevisiae* snoRNA database^41^, and snoRNA gene coordinates were obtained from the Saccharomyces Genome Database.^98^

### Zipper stem identification and stability calculation

We identified the positional constraints for forming a stem between the 5’ splice site and branch point sequence (termed “zipper stems”) by modeling 3D structures for introns in the context of the spliceosome using the Rosetta protocol FARFAR2.^45^ As test constructs, we used variants of the intron in RPS17B since a zipper stem in this sequence has been characterized.^16^ To identify the minimum linker length for which formation of a zipper stem is sterically compatible with an intron binding to the spliceosome, we built models as described in the following paragraph for variants of the RPS17B intron with a range of linker lengths between the 5’ splice site, zipper stem, and branch point. We based models on the cryo-EM structure of the *S. cerevisiae* A-complex spliceosome (PDB ID: 6G90).^99^ The tested variants included 36, 40, 45, 50, and 56 total nucleotides of linker sequence between the 5’ splice site, zipper stem strands, and the branch point sequence. All models were built using the Rosetta software (release v3.10) using the general approach for RNA homology modeling with Rosetta described previously.^100^

For each modeled RPS17B intron variant, we used Rosetta’s rna_thread application to replace the nucleotides proximal to the 5’ splice site and branch point that were resolved in the A-complex spliceosome structure with the sequence from RPS17B. We docked the RPS17B zipper stem into the binding site for an intron stem in the U1 snRNP, as determined from the E state spliceosome structure.^25^ More specifically, we aligned the U1 snRNP from the E-complex spliceosome structure including this zipper stem (PDB ID: 6N7R^25^) onto the U1 snRNP from the A-complex in PyMOL; we then saved the coordinates of the zipper stem relative to the A-complex as a rigid body. For each variant, 1250 structures were sampled using Rosetta’s rna_denovo application. The A-complex spliceosome structure, docked zipper stem, and rethreaded RPS17B intron residues were treated as rigid bodies through these modeling runs, and remaining linker residues and other positions in RPS17B were sampled freely. To accelerate structure sampling in the large context of the spliceosome, we generated models using a simplified score function that only included three terms: rna_vdw at weight 10.0 (the RNA van der Waals score term), rnp_vdw at weight 50.0 (the RNA-protein van der Waals score term), and linear_chainbreak at weight 10.0 (a score term penalizing chain breaks). Out of 1250 sampled structures, we chose the top 10 structures based on Rosetta score using these three terms. We collected the linear chain break score for each sampled structure, a score term which penalizes breaks in covalently attached neighboring residues that can occur when linker lengths are too short to stretch between docked nucleotides.

We note that the 3D modeling approaches used here make numerous approximations that may lead to model inaccuracies. First, we include components like the zipper stem docked to the A-complex spliceosome as a rigid body, simplifying a challenging modeling problem with many protein and RNA residues. Treating these components as rigid bodies precludes capturing local dynamics in these components that occurs in the context of full-length pre-mRNA. Second, we use a simplified score function with only van der Waals and chain break penalties, and we do not model explicit waters and ions surrounding our system. Without additional score terms (e.g. score terms that capture electrostatics), this simplified setting will not allow for modeling specific interactions between positions with accuracy. However, these simplifications enable fast scoring for a large system while still penalizing steric clashes between components and chain breaks within single strands. Finally, since these intron structures highly flexible, we expect that this system is best modeled with an ensemble of conformations; however, we only analyze a small set of structures to obtain statistics. We would not expect to be able to validate these flexible structures with experimentally determined 3D structures. In this setting, we emphasize that these structural models are used to obtain statistics on pre-mRNA length constraints rather than to analyze detailed structural features and interactions. Therefore, despite modeling limitations, this structural modeling can provide guidelines for zipper stem formation.

Based on the linear chain break scores for the tested series of linker lengths, to designate an intron stem as a zipper stem we required at least 42 total nucleotides between the 5’ splice site and the zipper stem’s 5’ end, and between the zipper stem’s 3’ end and the branch point. Additionally, we required that these stems begin at least 10 nucleotides from the 5’ splice site and end at least 20 nucleotides from the branch point, as positions more proximal to the splice sites were resolved in the A-complex spliceosome structure and would not be able to form base-pairs. Finally, we required that zipper stems are separated from the 5’ splice site and branch point at most 85 total linker nucleotides, so that they serve to co-localize the 5’ splice site and branch point. For a given secondary structure, we first identified the longest zipper stem in these windows allowing for at most 5 bulge nucleotides, and we computed the stability of this stem by finding the dG of folding in RNAcofold^101^ as implemented in Vienna 2.0^50^. In cases where no stem was present that satisfied these positional constraints in the secondary structure, we gave the intron a best zipper stem dG of 0 kcal/mol. Vienna 2.0 carries out free-energy calculations using a nearest neighbor energetic model, which computes the total free energy of stem formation as the sum of free energy contributions for each pair of stacked base-pairs (including enthalpic and entropic contributions). As a caveat, these calculations will not capture the energetics of non-local interactions that might be modeled by sampling 3D structures. In this case, free energies are computed at the default temperature (37C) and salt concentration (1.021 M); changing these parameters would alter computed free energies.

### Analyzing intron secondary structures surrounding canonical splice sites

We first designated intervals surrounding the 5’ splice site, branch point, and 3’ splice site where pre-mRNA secondary structures would be expected to clash with the spliceosome by analyzing spliceosome structures through the stages of splicing. We identified the number of nucleotides surrounding these splice sites that threaded through the spliceosome in structures of the E, A, pre-B, B, Bact, C, and C* complexes.^25,99,102–106^ We noted that the following nucleotides were built as single stranded nucleotides threading through the spliceosome in at least one of these spliceosome structures: 16 nucleotides upstream and 10 nucleotides downstream of the 5’ splice site; 33 nucleotides upstream and 21 nucleotides downstream of the branch point; and 8 nucleotides upstream and 20 nucleotides downstream of the 3’ splice site. We expected that these sequence intervals would be depleted of structure. For all 161 introns passing coverage thresholds, we generated DMS-guided secondary structure predictions for pre-mRNA intervals including the intron along with 50 nucleotides upstream and downstream of the intron for surrounding sequence context. We analyzed the proportion of nucleotides occluded by high confidence stems in these intervals. We additionally assessed the relationship between nucleotide protection in these intervals and retained intron fractions.

### Analyzing intron secondary structures surrounding cryptic splice sites

We identified cryptic splice sites by searching for alternate 5’ splice site, branch point, and 3’ splice site consensus sequences in relevant intron regions. More specifically, we searched introns for additional instances of any 5’ splice site, branch point, and 3’ splice site sequences that appear in either standard introns or previously annotated proto introns.^78^ We noted that at least 42 nucleotides were required between the 5’ splice site and branch point sequence for pre-mRNA to be compatible with spliceosome structures (see above). Thus, we located cryptic 5’ splice site sequences by searching for the sequences matching the 5’ splice site consensus that were located between the canonical 5’ splice site and 42 nucleotides upstream of the canonical branch point. Similarly, we located cryptic branch point sequences by identifying sequences matching the branchpoint consensus between 42 nucleotides upstream of the 5’ splice site and the canonical 3’ splice site. Finally, we identified cryptic 3’ splice sites that were at least 10 nucleotides downstream of the branch point sequence and upstream of the canonical 3’ splice site sequence. For all 161 introns passing coverage thresholds, we generated DMS-guided secondary structure predictions for pre-mRNA intervals including the intron and 50 nucleotides of surrounding context. We evaluated the protection of these cryptic sites by high confidence stems, comparing to the background protection of nucleotides in each region. Cryptic 3’ splice sites showed enriched protection by secondary structure relative to background nucleotides in these regions.

We additionally tested whether cryptic 3’ splice sites used only upon Prp18p inactivation (GAG, UG, CG, and GG splice sites) were protected by secondary structure,^107^ also finding a significant enrichment of secondary structure at these secondary cryptic 3’ splice sites relative to surrounding background (126/585=21.5% protection at these sites relative to 821/5926=13.8% protection at background sites; p-val < 0.001). However, cryptic 3’ splice sites matching the standard consensus sequence (UAG, CAG, or AAG) were significantly more protected (19/61=31.1% protection, Fig. S13D).

### DMS reactivity and structure analysis for introns

We made reactivity-guided secondary structure predictions to identify structural features of *S. cerevisiae* introns, making predictions for all introns and coding regions meeting the coverage threshold. Secondary structure predictions were made separately for introns and coding regions using *in vivo* DMS-MaPseq data and *in vitro* refolded DMS probing data. We restricted our coding region structure predictions to genes containing introns. We counted the number of introns that included zipper stems and stems between the branch point and 3’ splice site, using the following criteria. Our zipper stem definition was based on modeling of zipper stems in the context of the spliceosome as described above. We additionally identified “downstream stems” between the branch point and 3’ splice site with length at least 6 base-pairs and bootstrapping support at least 70%.

To calculate secondary structure properties for each intron and coding region with sufficient coverage, we represented the secondary structure for each sequence as a fine-grained and coarse-grained graph. Fine-grained graphs included a node for every base-pair and single-stranded position, with edges between consecutive positions or base-pairs. Coarse-grained graphs included a single node for each stem, junction, loop, single-stranded 5’ end, and single-stranded 3’ end. Fine-grained graphs were used to compute the maximum extrusion from ends metric, using the arnie package^108^ to find the length of the shortest path from each position in the sequence to either end of the sequence. The maximum value for this shortest path across all nodes in the network yielded the maximum extrusion from ends for the sequence, corresponding to the secondary structure distance between the sequence ends and the position furthest from these ends. The maximum extrusion from ends values were normalized by the length of each construct and compared between intron and coding regions with the Mann-Whitney U rank test. Coarse-grained graphs were used to identify the longest stem with at most 10 total loop nucleotides and at least 90% bootstrapping probability. Additionally, coarse-grained graphs were used to compute average helix confidence estimates for all stems of length at least 6 in introns and coding regions meeting coverage thresholds. The average helix confidence estimates and longest stems were compared between intron and coding sequences with nonparametric Mann-Whitney U rank tests.

Gini coefficients for comparing intron and coding reactivity values were calculated based on mutational frequency values for intron and coding regions. We obtained mutational frequencies across intron and coding sequences in all windows of 20 nucleotides, scanning in intervals of 10 nucleotides. For *in vivo* samples, we required that every position in each included window had per-position coverage at least 6456 as computed from rf-rctools view, corresponding to a 0.6 Pearson correlation between replicates for normalized reactivities. To compare *in vivo* to *in vitro* reactivity windows, we required that all positions in each window had at least 3160 coverage (the coverage cutoff used for the *in vitro* reactivity profiles). Gini coefficients across all passing intron and coding regions were compared with a nonparametric Mann-Whitney U rank test.

### Modeling of intron stems in spliceosome structure

We modeled the RPL28 and RPL36B introns in the context of the A-complex spliceosome (PDB ID: 6G90)^99^, using a protocol similar to the RPS17B zipper stem modeling as described above. Intron nucleotides resolved in 6G90 were replaced by the corresponding sequences from the intron in RPL28 and in RPL36B with the rna_thread application as discussed in the previous section.^109^ The secondary structure used for modeling was obtained from M2-seq guided RNAstructure prediction, and helices were supplied as rigid bodies for modeling. 4564 models were sampled using the Rosetta application rna_denovo.^45^ To accelerate structure sampling, we ran modeling with only rna_vdw, rnp_vdw, and linear_chainbreak score terms as described previously. The top-scoring models were visualized in PyMOL using the RiboVis package (https://ribokit.gihub.io/RiboVis/). This 3D modeling involved making numerous simplifying approximations, as discussed in the earlier section on the RPS17B stem modeling. However, because these models are used only to observe the size of intron structures relative to the spliceosome rather than make inferences on detailed interactions between components, this simplified spliceosome modeling effort can provide insights.

### Comparison of introns with decoys

To determine whether secondary structure features distinguish authentic introns from unspliced genomic sequences that match splicing motifs, we assembled a set of decoy introns from the genome. To find these decoy introns, we first computed the position weight matrix (PWM) for the 5’ splice site (6 nucleotides with consensus GUAUGU), branch point (8 nucleotides with consensus UACUAACN), and 3’ splice site (3 nucleotides with consensus YAG) for all canonical annotated introns in *S. cerevisae*, finding the log of the fraction of sequences that included each of the 4 nucleotides for every position in the sequence motif. We then obtained three length distributions for annotated *S. cerevisiae* introns: the complete intron length, the distance between the 5’ splice site and branch point, and the distance between the branch point and 3’ splice site. Gene annotations were obtained from the sacCer3 UCSC genome assembly. To find candidate decoy intron windows, we scanned across all genes for intervals containing 5’ splice site, branch point, and 3’ splice site sequences matching the PWMs and three length distributions for introns, using PWM score and length cutoffs that captured at least 95% of canonical introns. We filtered these sequences to exclude annotated introns. The resulting sequences represented “decoy introns”, transcribed gene intervals that matched splice site motifs and intron length distributions and yet were not spliced. For each authentic intron and decoy sequence, we assembled length-matched control sequences that were shifted in the genome by 500 nucleotides outside the intron sequence.

DMS-guided structure predictions were made for each intron, decoy, and matched control sequence. Sequences were only considered if both the construct and its shifted genomic control had sufficient coverage from DMS-MaPseq for structure prediction. For each secondary structure, we computed zipper stem dG values, downstream stem dG values, maximum extrusion from ends, and longest stem lengths as described above. Additionally, we computed the distance between the 5’ splice site and branch point in the intron’s secondary structure by computing the length of the shortest path in secondary structure graph between the 5’ splice site and branch point. We then determined whether authentic introns and decoy intron sequences enriched for these structural features more than shifted genomic controls. We compared the resulting values between these sets using the non-parametric Wilcoxon ranked sum test. We displayed the difference between intron and control feature values in violin plots after normalizing each metric between 0 and 1 and computing the feature value percent change.

### Visualizing the structure landscape for *S. cerevisiae* introns

We assembled secondary structure features for all introns passing DMS-MaPseq coverage thresholds and used these features to classify *S. cerevisiae* introns. For each intron, we included the following metrics which were calculated as described previously: the total sequence length, the presence of a zipper stem (binary 0 or 1), zipper stem free energy, the presence of a downstream stem (binary 0 or 1) between the branch point and 3’ splice site, downstream stem free energy, maximum extrusion from ends, the longest stem length, the average stem helix confidence estimate, the maximum Gini coefficient window, and average DMS accessibility for the 5’ splice site, branch point, and 3’ splice site. Hierarchical clustering was performed using scipy’s hierarchy module^110^ with the Ward linkage criteria and optimal leaf ordering, generating 9 clusters, and seaborn’s clustermap was used to generate a dendrogram and heatmap visualization. To create more interpretable classes, starting from the hierarchical clustering classes, we combined two clusters to generate Class 2 (zipper stems), reordered dendrogram classes to place Class 3 (downstream stems) after Class 2, and moved two short introns from Class 8 (unstructured long introns) to Class 7 (unstructured short introns). Additionally, tSNE classification was performed with all structural features using sklearn, with 2 embedded space dimensions, a perplexity of 40, and 300 optimization iterations.

### Designing intron structure variants for structure-function assay

We chose 7 structured introns to mutate systematically in our variant library by identifying secondary structures with zipper stems, covarying residues, and long stems. Intron library sequences were constrained to 300 total nucleotides including fixed primer binding sites and extensions into the 5’UTR for inserting barcodes, leaving around 200-250 nucleotides from the 5’ splice site in introns as the variable region for each construct. For each intron, we designated stem and loop sets to mutate within these length constraints, with each stem set containing one or more continuous stems in the secondary structure, and each loop set containing a single junction or loop in the secondary structure.

For each stem set and loop set, we generated candidate variants and rescue sequences where available, and we scored these candidates to find a final set of designs based on desired secondary structure properties. For each stem set, to generate candidate library sequences, we generated 10000 randomized nucleotide sequences and 10000 shuffled sequences mutating the 5’ strand of the stems. Additionally, if the 3’ end of the stem set fell within the length constraints of the library, we generated rescue sequences for each of these mutation candidates, installing compensatory mutations in the 3’ strand. For each loop set, we generated 20000 random variants as discussed above for all nucleotides in the loop accessible within the library. For each variant and rescue sequence, we computed the minimum free energy structure with Contrafold^111^ after placing the intron variable region in the full intron context.

We then assessed whether the variant disrupted any targeted stem structures while leaving the remainder of the secondary structure unchanged. More specifically, for each variant, we accumulated any junction or stem residues in the stem or loop set, and we computed two penalty scores: a variable region penalty and a constant region penalty. The variable region penalty was the sum of the paired residues in the variable region along with the number of base-pairs present in this region in the variant. The constant region penalty focused on nucleotides outside the variable region, computing the number of base-pairs in this region of the native secondary structure disrupted by the variant, and adding half of the number of base-pairs gained in the variant. If the variant also had a rescue sequence available, we computed the rescue penalty as the number of base-pairs lost or gained between the native secondary structure and the rescue sequence. The final total score for the variant sequence was the sum of the variable region penalty, constant region penalty, and the rescue penalty. We ranked variants by this total score and chose the top 4-8 unique variants per region. We visually checked the base-pair probability matrices for wildtype, variant, and rescue sequences to ensure disruption and rescue of stems in the desired sequence interval. Additionally, we ensured that variants did not introduce BglII and XcmI cut sites, as these restriction enzymes would be used subsequently for library integration. Our variants did not mutate the 5’ splice site, branch point, or 3’ splice site sequences. For each variant set, we additionally included 4-8 copies of the wildtype sequence to ensure sufficient coverage of the native intron in our experiment. The final chosen variant sequences along with the variable region penalty, constant region penalty, and total penalty scores are in Table S3.

### Designing gene context for intron variants

Introns were cloned into their native gene context to better mimic the effects of structures in native *S. cerevisisae* pre-mRNA on gene expression. For each construct, we included the promoter TDH3, the 5’ UTR beginning at the transcription start site, the full gene including the intron sequence, and the ADH1 terminator (Fig. S19C). The TDH3 promoter was chosen for yielding high expression levels,^49^ and the ADH1 terminator was chosen for yielding high mRNA half life.^112^ For each gene, we used the YeasTSS database to find the most prominent transcription start site upstream of the gene, using this position as the start of the 5’UTR in our constructs.^113^ When cloning intron libraries into this gene context, we included a short 8-nucleotide fixed sequence in the 5’UTR to distinguish the inserted gene from the endogenous gene copy (AGCGGACG for QCR9, RPL28, RPS9A and RPL36B; AGAAGACG for RPS14B; AGAAGAGC for RPL7A; and AACTGCCC for RPS9B). Additionally, we included a random 12-nucleotide (12N) barcode ahead of the intron libraries in the 5’UTR that we aimed to use to identify pre-mRNA variants even after introns are spliced out. We ran simulations by randomly sampling 12N barcodes to check that for our expected number of E. coli and yeast library clones (10,000-50,000), we would not have substantial overlap between barcodes for separate clones, with most barcodes expected to have at least edit distance two from all other barcodes (Fig. S19D). We therefore expected that these random barcodes would allow us to unambiguously assign spliced reads to intron variants.

### Growth media for structure variant library assay

We used the following growth media and plates for our structure-function assay:

- YPD media: YPD powder (Difco; Yeast extract 10g/L, Peptone 20g/L, Dextrose 20g/L)
- YPD plates: Difco YPD powder 50g/L, Difco agar powder 20g/L
- SD-LEU plates: Premix from Takara Bio
- SD-HIS plates: Premix from Takara Bio

### Constructing yeast background strain for structure variant library integration

We aimed to integrate our variant library into the yeast genome using homologous combination and CRISPR/Cas9-induced double stranded breaks to increase efficiency. We chose the ARS416 locus as our integration site, as it was previously identified as a highly efficient integration site when using Cas9-facilitated genomic integration.^114^ To ensure integration of our library in the expected locus, we first constructed the background strain PLH001 that inserted a partial LEU2 selection marker at this locus (starting from the second codon of the coding sequence). We then designed our genomic inserts for our structure variant library to include the LEU2 promoter and start codon, such that genomic integration of the library at the correct locus was required to complete the LEU2 selection marker and allow for selection with SD-LEU.

The strain PLH001 was constructed from background strain BY4741 (Mata, his3Δ1, leu2Δ0, met15Δ0, ura3Δ0) to serve as a landing pad for genomic integration of the structure variant library. A partial LEU2 selection marker and a complete HIS3 selection marker were integrated into the ARS416 gene locus immediately downstream of the previously tested CRISPR/Cas9 cut site.^114^ To clone the construct for yeast genomic integration, LEU2 and HIS3 were obtained through PCR from the pHLUM plasmid,^115^ adding homology arms for genomic integration into ARS416. These constructs were cloned into the pUC19 backbone^116^ using NEBuilder Hifi DNA Assembly (NEB) (Fig. S19C, insert sequence RRLH in Table S5). We obtained a linear insert for transformation with PCR and integrated into the ARS416 locus with heat shock transformation of BY4741 using SD-HIS selection.^117^

### Constructing plasmid backbones for structure variant library

We first cloned a plasmid backbone for each intron variant containing the intron’s gene context (pRR1-pRR7), and we then cloned in our intron libraries into this background with a high efficiency transformation protocol. For each intron, the background gene consisted of the TDH3 promoter, the 5’UTR, the gene and intron excluding the variant library, the ADH1 terminator, and homology arms for insertion into the ARS416 locus (homologous to ARS416 on the 5’ end and part of the inserted LEU2 selection marker on the 3’ end) (Fig. S19C). All PCR reactions in the following section followed the following protocol unless otherwise specified. We used 25 μL of the 2X NEBNext Ultra II Q5 Master Mix (NEB) with 2.5 μL of 10 μM primers using the following thermocycler conditions: denaturation at 98 °C for 30 seconds; 30 cycles of 98 °C for 10 seconds, 30 seconds at the primers’ annealing temperature, and 72 °C for 30 seconds; and a final elongation at 72 °C for 10 minutes.

We used the following gene fragments from IDT to build gene background constructs: 1. Gene blocks with the homology arm and TDH3 promoter; 2. A gene block with the ADH1 terminator, LEU2 promoter, LEU2 start codon, and LEU2 homology arm; and 3. Gene blocks with the target gene and constant regions in the intron (gblockRR4-12; Table S5). Gene blocks were amplified with PCR using primers RR222-RR233, adding homology sequences used for construct assembly (Table S5). pUC19 was linearized with PCR and treated with DpnI (NEB) to remove intact plasmid template. Gene blocks and the background plasmid were cloned using a 4-part NEBuilder Hifi DNA Assembly (NEB) reaction.

### Cloning structure variant library with high efficiency transformation

Intron library sequences for the 7 intron sub-libraries were obtained as a single oPool from IDT, and the 7 sub-libraries were then amplified by PCR. For each intron, we carried out 8 parallel PCR reactions with limited PCR cycling to reduce bias. We used the following primer pairs: RR273, RR249 for RPS14B; RR274, RR251 for RPS9B; RR275, RR253 for RPS9A; RR276, RR255 for QCR9; RR277, RR257 for RPL36B; RR279, RR261 for RPL28; and RR280, RR263 for RPL7A (Table S5). Forward primers for these PCR reactions included 12 nucleotide random barcodes and 5’ homology to the corresponding plasmid backbone (pRR1-pRR7), and reverse primers included 3’ homology to the plasmid backbone. We used the Q5 PCR amplification protocol from the previous section for each of the 8 reactions, but in this case with 30 ng template, 1 minute extension time, and 14 amplification cycles. The 8 PCR reactions for library inserts were concentrated together with the QIAQuick PCR Purification Kit (Qiagen) and gel purified with a 1% agarose gel and MinElute PCR Purification Kit (Qiagen). Plasmid backbones pRR1-pRR7 were linearized via PCR primers RR223 and RR239-RR245 using the Q5 protocol from the previous section with 3.5 minutes extension time, and then purified from 0.8% agarose gels with the MinElute PCR Purification kit, and treated with DpnI (NEB). Intron library inserts were assembled into these linearized backbones used NEBuilder HiFi DNA Assembly (NEB) with 100 ng backbone and 3X molar excess of library inserts.

For library cloning, we transformed NEB Stable Competent *E. coli* cells (NEB) with NEBuilder assembly reactions for each of our 7 sub-libraries using the high efficiency transformation protocol from the manufacturer. We used 2.5 μL ligation product and 32 μL competent cells per reaction. For each intron, we plated 100 μL of cells on each of 3 LB plates with carbenicillin. We obtained 6000-46000 clones per sub-library (30-230X library coverage), and we found correct designed variants in 50-100% of the clones from Sanger sequencing of 8-16 clones per sub-library. Plasmid DNA was extracted by scraping cells and then using the ZymoPURE II Plasmid Midiprep Kit (Zymo Research).

### Yeast genomic integration for structure variant library assay

The plasmid library was linearized by restriction digest and transformed into PLH001 for genomic integration. For the first digestion, we combined 3 μL of XcmI (NEB) with 10 μL 10X R2.1 buffer (NEB), 5 μg of each plasmid sub-library, and H_2_O to 100 μL. Reactions were left at 37 °C for 60 minutes, 65 °C for 20 minutes, and then on ice for 5 minutes. Then for the second digestion, we added 2.0 μL of 2.5 M NaCl, 4.16 μL of Tris HCl pH 7.9, and 1.5 μL of the BglII enzyme, followed by an incubation at 37 °C for 60 minutes. To increase efficiency of integration, the linearized plasmid library was transformed with Cas9 to encourage genomic double-stranded breaks at the integration site. We obtained 50 μg of plasmid p426^114^ with the ZymoPUREII Plasmid Maxiprep Kit (Zymo Research) and linearized with AhdI digest.

We used heat-shock transformation in PLH001 with the linearized Cas9 plasmid and library insert for genomic integration with the LiAc/ssDNA/PEG transformation protocol.^117^ For each sub-library transformation, we used 5 μg linearized plasmid library and 5 μg linearized p426 plasmid to transform 25 mL of OD600 0.5 cells. We plated transformed cells on SD-LEU selection plates, with 4 plates per sub-library. As negative controls, we transformed DNA from one sub-library into BY4741 (without the partial LEU2 landing pad in the genome), and we transformed p426 alone without a library insert into PLH001. We saw no colony growth for either of these negative control conditions on SD-LEU selection plates. We scraped colonies from all library transformation plates and stored in 15% filtered glycerol at -80 °C.

Dilution plates from the sub-library transformations were used for counting colonies and verifying genomic integration. We counted 9000-35000 colonies for each sub-library, yielding 45-175X library coverage for each intron. To verify genomic integration, we extracted genomic DNA from 4-8 colonies per sub-library with the YeaStar Genomic DNA Kit (Zymo Research) and sequenced PCR amplicons with Sanger sequencing, finding that 57-100% of clones were successfully integrated.

### Genomic DNA extraction and sequencing for structure variant library assay

We sequenced genomic DNA to assign barcodes to each designed variant sequence and to obtain transformation frequencies. For genomic DNA extraction, we started overnight cultures from 100 μL of yeast sub-library glycerol stocks. Cultures were diluted and grown to OD600 0.5-0.6 the following day. We used the YeaStar Genomic DNA Kit (Zymo Research) for DNA extraction.

To sequence genomic DNA, we first carried out an indexing PCR with limited cycles to amplify target intron library regions and add i5/i7 indices for Illumina sequencing. We used the primers RR281-RR294 to amplify our sub-libraries and add Illumina adapters (Table S5). These primers amplified the target sub-libraries including barcodes and complete variant sequences, with forward primers binding to the fixed 8-nucleotide sequences that distinguished the inserted intron library from the endogenous gene copy. Additionally, these primers included unique i5 and i7 index sequences for demultiplexing sub-libraries. For each indexing PCR, we used a 50 μL PCR reactions using Q5 (NEB) with 50 ng genomic DNA template and 7 amplification cycles. We purified PCR reactions with the DNA-5 Clean & Concentrator-5 kit (Zymo Research). Next, we ran qPCR reactions with two replicates per sub-library to determine the number of cycles of additional PCR required for sequencing. We then amplified our indexing PCR sample with P5/P7 primers, again using the Q5 protocol described earlier, now with 11-14 cycles and annealing at 66 °C. Samples were purified using RNACleanXP beads (Beckman Coulter) with a 0.8 Ampure bead to sample volume ratio for size selection, aiming to select amplicons and remove any primers or primer dimers from the sample. The sequencing libraries were quantified with a Qubit high sensitivity dsDNA Assay Kit (Invitrogen) and the libraries were sequenced using an Illumina MiSeq v3 600-cycle kit, providing paired-end reads of length 300 each.

### RNA extraction and targeted RNA sequencing for structure variant library assay

We aimed to sequence RNA to obtain levels of spliced and unspliced RNA. For RNA extraction, we again started overnight cultures from 100 μL of yeast sub-library glycerol stocks. For RNA extraction, when cultures reached OD600 0.5-0.6 the following day, we used 5 mL culture per sub-library in two columns from the YeaStar RNA Kit (Zymo Research), with 2.5 μL of Zymolase for every 2.5 mL starting cell culture.

For RNA sequencing, many library preparation steps mirrored our DMS-MaPseq library preparation described earlier. We pooled RNA extracted from each sub-library, combining 50 μg total RNA from each sub-library to make 350 μg total RNA. As described previously, we first concentrated extracted RNA with the RNA Clean and Concentrator-5 kit (Zymo Research), and then we depleted rRNA from our total RNA with RNAseH, using 50-mer oligos to tile rRNA sequences. We then carried out size selection with RNA Clean and Concentrator-5 kit (Zymo Research) to remove the smaller rRNA depletion oligos and small RNAs from the total RNA sample. We DNAse treated each reaction with 10 μL TURBO Dnase and a 30 minute incubation at 37 °C, and we purified reactions with the RNA Clean and Concentrator-5 kit (Zymo Research). After rRNA depletion, size selection, and DNAse treatment, we were left with 1.56 μg RNA.

To assess the removal of contaminating genomic DNA from our total RNA, we carried out a reverse transcription reaction with and without reverse transcriptase and checked for amplification of control RNA regions with PCR. For reverse transcription, we used the iScript Reverse Transcription Supermix (Bio-Rad), with 50 ng of our DNAse-treated RNA in samples with and without the reverse transcriptase. We purified samples with Oligo Clean and Concentrator columns (Zymo Research), eluting in 15 μL H_2_O. We then carried out PCR with the MATa1 and RPL36B primers as described in the section “RT-PCR for verifying splicing inhibition”. We visualized products on a 2% E-Gel EX Agarose Gel (Thermo Fisher).

We next generated cDNA that would be used for targeted PCR amplification and sequencing from our RNA sample. We chose to generate cDNA after fragmenting RNA and ligating on a universal cloning adapter as a handle for reverse transcription. Compared to library preparation approaches that would reverse transcribe RNA with targeted primers downstream of the intron, we expected that our approach would avoid large length differences between unspliced and spliced amplicons, which could lead to biases in reverse transcription and PCR amplification. Our approach mirrors the library preparation strategy described previously for DMS-MaPseq. Briefly, first, we fragmented our RNA and purified with the RNA Clean and Concentrator-5 kit (Zymo Research). We then removed 3’ phosphate groups from fragmentation with rSAP (NEB) treatment followed by rSAP inactivation. We prepared adenylated linker from RR118 (Table S5), and we then ligated this linker to our RNA sample with a T4 RNA ligase 2 truncated KQ (NEB) reaction, adding adenylated linker in 2X molar excess of the RNA sample. Excess linker was degraded with 5’ Deadenylase (NEB) and RecJf (NEB) exonuclease. The sample was cleaned with the RNA Clean and Concentrator-5 kit to prepare for reverse transcription. For reverse transcription, we used TGIRT enzyme (InGex) with the protocol described previously, using the reverse transcription primer RR301 which includes a 10 nucleotide UMI and Illumina adapters (Table S5). We degraded RNA and then purified cDNA with an Oligo Clean and Concentrator column (Zymo Research).

Next, we amplified cDNA with targeted PCR primers for each sub-library to generate final sequencing samples. We first added i5/i7 indices for Illumina sequencing along with Illumina adapters in a short indexing PCR for each sample. For these PCR reactions, we used the following primer pairs: RR281, RR302; RR283, RR303; RR285, RR304; RR287, RR305; RR289, RR306; RR291, RR307; and RR293, RR308 (Table S4). Our forward PCR primers were complimentary to cDNA just 5’ of our barcode sequences, and they included the 8-nucleotide fixed sequence that distinguished the inserted library of interest from the endogenous gene. These primers allowed for the amplification of cDNA including barcode sequences and the splice site junction. Additionally, these primers included unique i5 and i7 index sequences, allowing libraries to be demultiplexed after sequencing. For this indexing PCR, we used the Q5 reaction protocol described above, in this case with 7 amplification cycles. To obtain sufficient material for sequencing, we then carried out 24-28 additional PCR cycles with P5/P7 primers (Table S5), using Q5 with the same cycling parameters as the indexing PCR reaction. Finally, to obtain dsDNA for sequencing, we carried out two rounds of size-selection and purification with RNACleanXP beads (Beckman Coulter), using a 0.7X Ampure bead to sample volume ratio and eluting in 15 μL H_2_O. The sequencing libraries were quantified with a Qubit high sensitivity dsDNA Assay Kit (Invitrogen) and sequenced using a partial Illumina NovaSeq S4 lane, providing around 700 million paired-end reads of length 150.

### Genomic DNA sequencing data analysis for structure variant library

Genomic DNA sequencing provided links from barcodes to intron variants, a critical step for quantifying spliced reads for intron variants since the barcodes remain in spliced mRNA while variant sequences are spliced out. Additionally, genomic DNA sequencing provides quantification for the number of transformants for each variant, providing a normalization factor for mRNA levels. To prepare genomic DNA sequencing reads for analysis, we first used UMI-tools^77^ to extract barcodes by matching primer binding sites immediately upstream of the barcode. We then error corrected paired-end reads using bbmerge,^118^ removing Illumina adapter sequences and ensuring consistency between paired-end reads.

We used the following pipeline to assemble consensus intron variant sequences corresponding to each barcode, making use of functions within the fgbio toolkit (fulcrumgenomics.github.io/fgbio/). First, paired-end reads were aligned to introns using bwa^119^ with default settings. The resulting alignments were sorted using fgbio’s SortBam directive and mate-pairs were annotated with fgbio’s SetMateInformation. With reads aligned to the reference introns, we then grouped reads with the same barcode using fgbio’s GroupReadsByUmi, choosing the adjacency method to combine barcode sets. Consensus reads were called using fgbio’s CallMolecularConsensusReads with mostly default settings except the flags -m 20 -M 5, which required that only bases with Q score above 20 were included, and that at least 5 reads covered each consensus sequence. We then quality filtered these sequences using fgbio’s FilterConsensusReads with the flags -M 5 -q 20 -N 20 -E 0.01, again using the same quality and read depth scores and additionally requiring that consensus bases have quality at least 20 and reads have average error rates lower than 0.01. With samtools,^81^ we sorted the resulting alignment file and retrieved paired-end consensus reads. We then merged paired-end reads using bbmerge^118^ with default settings, obtaining consensus length distributions depicted in Fig. S20A. Finally, to ensure that barcodes are mapped primarily to single consensus reads, we kept the primary consensus sequence for each barcode and required that at most 5% of reads map to secondary consensus sequences. This filter removed 0.4-4.7% of barcodes from further analysis across all sub-libraries.

From our consensus sequences, we obtained barcode mappings for designed variants along with transformation frequencies. We collated barcodes that mapped to designed variant sequences, finding at least 10 unique barcodes for many of the designed variants in each sub-library (Fig. S20B). This indicates that each designed variant was transformed into at least 10 *E. coli* and *S. cerevisiae* colonies, with some variants having much higher transformation frequencies. We additionally found that many colonies had consensus sequence variants that were mutations of wildtype sequences or designed variants. From these sequences, we retrieved all barcodes corresponding to QCR9 intron variants with branch point mutations, providing a control sample for our RNA sequencing analysis. To obtain transformation frequencies, we found total read coverage for each barcode using the output from fgbio’s FilterConsensusReads.

### RNA sequencing data analysis for structure variant library

Our targeted RNA-sequencing data provided normalized mRNA levels and retained intron fractions for each intron variant. To process RNA-sequencing data, we first extracted barcodes (one for each transformant) and UMI’s (one for each cDNA molecule) from our sequencing reads with UMI-tools.^77^ We then removed Illumina adapters and the ligated linker for reverse transcription from the reads using cutadapt, additionally trimming bases with Q-score lower than 20. We then aligned reads to spliced isoform reference sequences to filter out reads from our RNA-sequencing run that did not correspond to the desired target sequence. For our reference sequence annotations, we collated all possible isoforms capturing spliced and unspliced transcripts (4 isoforms for two-intron RPL7A and 2 for all other genes), and we generated gff annotation files with gmap.^120^ We used TopHat2^85^ to align sequencing reads to these reference isoform annotations, using the following flags to enable potentially distant alignments between variants and the reference sequence: --no-novel-juncs -T –b2-mp 3,1 –b2-rdg 5,1 –b2-rfg 5,1 –segment-length 20 –segment-mismatches 3 –read-gap-length 7 –read-edit-dist 50 -m 1 –max-insertion-length 19 –max-deletion-length 19. We generated fasta files from the resulting alignment files with samtools.^81^

We classified each aligned sequence into one of three categories: unspliced, spliced at the expected junction, or other. We required an exact agreement with 14 nucleotides representing the unspliced junction (for unspliced sequences) or spliced junction (for spliced sequences). For each barcode, we computed the number of spliced and unspliced reads, deduplicating UMI’s at this stage to reduce the effects of PCR amplification biases on our quantification. We then computed two key metrics for each barcode: the retained intron fraction and the normalized mRNA level. The retained intron fraction was computed as the ratio between unspliced read counts and total read counts (sum of spliced and unspliced reads). The normalized mRNA levels were computed as the ratio between the spliced read count and the transformation frequency obtained from genomic DNA sequencing.

To understand the prevalence of alternative splicing events, we further analyzed the sequences that did not include an exact match with spliced or unspliced junctions. To identify a novel event as an alternative splicing event, we required that event appear in at least 10 UMI’s per barcode and at least 3 barcodes per variant. We computed the fraction of transcripts for each variant that were alternatively spliced.

For each intron sub-library, we used the barcode-variant correspondence from genomic DNA sequencing to collect the retained intron fractions and normalized mRNA levels for all the barcodes corresponding to each variant sequence. For all the stem and loop sets assessed in each intron sub-library, we compared RI fractions and normalized mRNA levels between the following sets of sequences: the wildtype sequence versus stem disruption variants, and the stem disruption variants versus the rescue sequences. Comparisons were made using two-sided permutation tests (scipy^110^) for the difference in mean statistic with 100,000 resamples of data labels per comparison.

### RT-qPCR from individual variant strains to validate VARS-seq

We first cloned intron variants of RPS9A and RPL36B individually into the corresponding plasmid backbones to generate complete gene constructs with individual variant introns. For RPS9A, we cloned in 8 intron variants, including the wildtype, 5’ mutant, 3’ mutant, and rescue sequences for two different barcodes (from gblock_RR15-18 with two barcodes subsequently added by PCR). For RPL36B, we cloned 8 intron variants including the wildtype, 5’ mutant, 3’ mutant, and rescue sequences for two different stem sets labeled “short” and “long” (cloning from gblock_RR19-26). For RPS9A, gblocks were amplified by PCR with primers RR253 and RR337 to include the barcode expected to have high retained intron levels, and with primers RR253 and RR338 to include the barcode expected to have low retained intron levels. For RPL36B, gblocks were amplified with primers RR343 and RR345 for the short stem set and with primers RR345 and RR344 for the long stem set. Intron library inserts were assembled into the corresponding linearized backbones (pRR5 and pRR7) using NEBuilder HiFi DNA Assembly (NEB) as described above. 2.5 μL ligation product was transformed into 15 μL NEB Stable Competent *E. coli* cells (NEB) for each sample and correct clones were verified by Sanger sequencing.

We next integrated these intron variant constructs into *S. cerevisiae* to analyze the effects of these variants on splicing. Plasmids extracted from correct clones were linearized by restriction digest, with a first restriction digest by XcmI (NEB) and a second digest with BglII (NEB) as described above. These linearized plasmids included complete gene constructs for RPS9A or RPL36B with intron variants, homology arms to the genomic integration site, and a partial LEU2 selection cassette that would be completed by genomic integration into PLH001. Linearized plasmids were transformed into PLH001 with the LiAc/ssDNA/PEG transformation protocol described above. For each transformation, we used 250 ng linearized plasmid to transform 2.5 mL of OD600 0.5 cells. We plated transformed cells on SD-LEU selection plates. We verified correct genomic by extracting genomic DNA with the YeaStar Genomic DNA Kit (Zymo Research) followed by Sanger sequencing.

We next extracted RNA from each strain and evaluated levels of spliced and unspliced transcripts with qPCR. To extract RNA, we used the YeaStar RNA Kit (Zymo Research) with 5 mL OD600 0.5-0.6 culture as described above, eluting in Rnase-free H_2_O. We dNase treated the samples by mixing 1 μL of TURBO Dnase (Thermo Fisher), 5 μL of TURBO dNase 10X buffer, 2 μg RNA, and Rnase-free H_2_O for a 50 μL reaction volume. We incubated the reactions at 37 °C for 1 hour, added 20 μL of 50 mM EDTA, incubated the reactions at 75 °C for 10 minutes, and placed them on ice. We cleaned up samples with Zymo5 RNA columns and eluted in 15 μL Rnase-free H_2_O. We then carried out reverse transcription with the iScript Reverse Transcription Supermix (Bio-Rad), using 7.5 μL of each RNA sample for a reaction that included the reverse transcriptase, and the remaining 7.5 μL for a control without this enzyme. 4 μL of the 5X RT Supermix was mixed with 7.5 μL of RNA sample and water to a reaction volume of 20 μL, and the reaction was incubated in a thermocycler for 5 minutes at 25 °C, 40 minutes at 46 °C, and 1 minute at 95 °C. The samples were purified with Oligo Clean and Concentrator columns (Zymo Research) and eluted in 15 μL.

With this reverse transcription product, we then carried out qPCR with the iTaq Universal SYBR Green Supermix (Bio-Rad), using 1 μL of forward and reverse primer, 7 μL of Rnase-free H_2_O, 1 μL of the reverse transcription reaction, and 10 μL of the iTaq mix. Forward primers were designed to avoid the endogenous gene locus by including the fixed 8 nucleotide sequence inserted into the constructs (see the section “Designing gene context for intron variants” above). Reverse primers were designed for the RPS9A and RPL36B constructs to amplify only the spliced construct (binding to the exon junction) or only the unspliced construct (binding within the intron). For RPS9A we used primers RR313 and RR314 to amplify the spliced product, and RR317 and RR319 to amplify the unspliced product. For RPL36B we used primers RR346 and RR347 to amplify the spliced product, and RR346 and RR348 to amplify the unspliced product. A region of ACT1 was used as a control interval, with primers RR341 and RR342. All primers were verified by checking for the formation of a single band for each amplicon on agarose gels.

### *De novo* computational structure prediction and structure metrics

We predicted secondary structure ensembles for *S. cerevisiae* introns and control sequence sets with *de novo* structure prediction. For direct comparison between DMS-guided RNAstructure prediction and *de novo* RNAstructure prediction, we computed features in both cases only for the 161 introns with sufficient coverage from DMS-MaPseq; for other cases, we made predictions for the complete set of introns filtered to exclude sequences smaller than 50 nucleotides or larger than 600 nucleotides, leaving 288 of 297 introns. Three control sequence sets were assembled, each with length-matched control sequences for every intron. To construct the first set, the “shifted control”, the genome coordinates for each intron were shifted 500 nucleotides downstream of the intron’s 5’ start site and a sequence of the same length as the intron was taken from these shifted coordinates. We constructed the second control set, the “sequence-matched control”, by replacing the shifted control’s sequences at the positions of splice sites in the matched intron, substituting the 5’ splice site, branch point, and 3’ splice site sequences from the intron at these positions. This set was constructed to control for structural differences caused primarily by sequence preferences at these splice sites. The final set, the “shuffled control”, was obtained by shuffling each intron’s sequence randomly.

Secondary structure ensemble predictions for each set of sequences were obtained by either making minimum free energy predictions or by sampling suboptimal structures using computational secondary structure prediction packages. For suboptimal structure sampling, 1000 structures were sampled for each sequence using the Arnie package (https://github.com/DasLab/arnie) to call structure prediction executables from Vienna 2.0^50^ and RNAstructure^33^ with default parameters. We additionally made structure predictions for intron sequences with surrounding exon context, extending each intron to include 50 bases upstream and downstream of the intron. For each intron and matched control, we computed average values across these structure ensembles for the zipper stem free energy, downstream stem free energy, length of the longest stem, maximum extrusion from ends, and distance between the 5’ splice site and branch point. As described previously, we compared the intron and control values using the non-parametric Wilcoxon ranked sum test. When we compared feature values using a t-test for related samples, we identified the same comparisons as significant in all cases.

### Structure prediction for *Saccharomyces* genus

We obtained multiple sequence alignments for introns in 20 species in the *Saccharomyces* genus from Hooks, et al. 2014.^51^ We computed statistics on the alignments for each intron, finding its number of orthologs and the average percent sequence conservation. To count the number of orthologs for each intron, we found all the species with non-empty sequences aligned to the *S. cerevisiae* intron. We additionally counted the number of zipper stem orthologs by finding all the non-empty regions aligned to the longest zipper stem in each *S. cerevisiae* intron. Finally, we computed the average percent sequence conservation across each complete intron and across the intron’s zipper stem if one was present.

We next evaluated secondary structure feature enrichment for the introns across the *Saccharomyces* genus. We tested each intron in these 20 species against matched control sets, with each intron having a matching shuffled sequence control. We additionally compared the secondary structure ensembles of all the introns in each of these species’ genomes to phylogenetic controls, assessing whether they preserved structural features more than expected compared to sequences altered at similar levels from the *S. cerevisiae* intron homolog. To construct phylogenetic control sequence sets, we measured mutation and indel rates between each intron and its homologous *S. cerevisiae* intron, and we created an intron’s matched control sequence by inserting mutations and indels randomly into the homolog *S. cerevisiae* intron at this measured average frequency. As described earlier for the analysis of *S. cerevisiae* introns, for each of these comparisons, we predicted secondary structure ensembles for the intron set and control set, computed secondary structure features, and compared between sequence sets with a Wilcoxon ranked sum test.

### Reagent and resource sharing

Yeast strains are available upon request from the authors. Requests for resources and reagents should be directed to the corresponding author, Rhiju Das.

## Code availability

Code for assessing DMS-MaPseq reactivity values, evaluating control RNA structures, and analyzing DMS-derived secondary structures is included here: https://github.com/ramyarangan/DMS_intron_analysis. Code for computational secondary structure prediction, secondary structure feature calculation, statistical comparisons, and evolutionary analyses are included at https://github.com/ramyarangan/pre-mRNA_Secstruct. Code for designing VARS-seq sequences and analyzing gDNA and RNA sequencing data from VARS-seq is included in https://github.com/ramyarangan/VARSseq.

## Data availability

All sequencing data are available at the Gene Expression Omnibus (GEO) accession number <u>GSE209857</u>. DMS-derived secondary structures are included in Table S2. Processed DMS reactivity data, control RNA secondary structures, and base-pairing probabilities from structure prediction with bootstrapping are included in https://github.com/ramyarangan/DMS_intron_analysis. VARS-seq intron variant sequences along with spliced and unspliced read counts are included in Tables S3-S4.

## Acknowledgments

We thank K. B. Hooks and D. Delneri for providing secondary structure data upon request. We thank A. Hoskins, J. Puglisi, T. Lan, and S. Rouskin for illuminating discussions on this project, and we thank V. V. Topkar and I. Zheludev for advice on experimental protocols. We thank Stanford University for providing computational resources and support for the Sherlock 2.0 cluster that contributed to these results. This work was supported by the National Science Foundation Graduate Research Fellowship Program grant no. 1650114 (R.R.); the Gerald J. Lieberman Fellowship (R.R.); the National Institutes of Health grant R35GM145266 (M.A.), a Stanford Bio-X Interdisciplinary Initiative Award (R.D.); the National Institutes of Health grant R35GM122579 (R.D.); and the Howard Hughes Medical Institute (HHMI) Investigator Program (R.D.). This article is subject to HHMI’s Open Access to Publications policy. HHMI lab heads have previously granted a nonexclusive CC BY 4.0 license to the public and a sublicensable license to HHMI in their research articles. Pursuant to those licenses, the author-accepted manuscript of this article can be made freely available under a CC BY 4.0 license immediately upon publication.

## Author information

R.R. and R.D. conceived the project. R.R. designed and performed experiments and carried out computational analyses. O.H. and M.A. developed the *S. cerevisiae* strain OHY001 used in the project. R.H. conducted *in vitro* DMS probing experiments. P.P. tested alternate structure prediction approaches. R.R., M.A., and R.D. interpreted the results. R.R. wrote the initial draft, and R.R., M.A., and R.D. wrote the final manuscript with input from all authors.

