## Supplementary Material for "RNA structure landscape of *S. cerevisiae* introns"

Supplemental Text

**Alternative DMS-guided structure prediction approaches.** We explored additional structure prediction approaches to identify potential pseudoknots and alternative conformations in introns using the DMS data. To find potential pseudoknots, structural motifs that often participate in stable three-dimensional RNA folds, we predicted structures using ShapeKnots^1^ guided by DMS data for each intron. No introns included predicted pseudoknots with helix confidence estimates exceeding the 70% confidence threshold. As a control for our pseudoknot predictions, we predicted the secondary structure for the RNase P RNA with ShapeKnots^1^ guided by DMS data. The known pseudoknot in RNAse P^2^ was recovered with this approach but only with 21% helix confidence estimate, suggesting that some pseudoknots in introns may be missed in our scan due to low sensitivity when using the 70% helix confidence threshold.

Additionally, for eight intron regions with high coverage (see Methods), we generated structure predictions using DREEM^3^ to test for alternate conformations represented by the DMS data. However, reactivity data for all eight regions were best explained by a single structure based on the Bayesian Information Criteria (BIC) test statistic reported by DREEM. We note that it is possible these introns have significant alternative conformations that would only become apparent when probing with higher DMS concentrations or sequencing more deeply.

**Detailed evaluation of proposed functional structures in *S. cerevisiae* introns.** Our DMS-MaPseq data after splicing inhibition allowed us to assess classes of previously proposed intron structures in *S. cerevisiae*. These classes include intron structures that were identified through mutational studies and functional experiments, along with introns identified through computational structure prediction and evolutionary analysis. We find that structures identified with functional experiments or scans for covariation were largely supported by our *in vivo* chemical probing data, whereas structures identified through other computational prediction approaches had limited support. Here, we include a detailed description of these classes of structures along with their support from our DMS data.

Prior studies have used experiments assessing the role of structure using mutants and compensatory mutants to pinpoint to potential regulatory structures in some *S. cerevisiae* introns. For instance, in the case of RPS17B, a stem linking the 5’ splice site to the branch point, termed a “zipper stem”, enables efficient splicing despite this intron’s weak 5’ splice site.^4,5^ A stem between the branch point and 3’ splice site in RPS23B hides a more proximal cryptic 3’ splice site and enables thermosensitive regulation of 3’ splice site selection for this intron.^6^ In the case of the introns in RPL32, RPS9A, and RPS14B, portions of the intron have been implicated in regulating gene expression by binding of excess protein product to pre-mRNA, leading to gene-specific reduced splicing efficiency.^7-10^ Finally, structures in RPL18A and RPS22B have been found to mediate the degradation of unspliced pre-mRNA.^11^

In four out of six cases, structures from prior functional experiments received medium or high support from our DMS data, with high loop reactivity, low stem reactivity, and high helix confidence estimates (Fig. 2). For instance, the structure in RPL18A involved in pre-mRNA degradation includes high confidence stems from DMS probing (Fig. 2A). DMS data additionally support the stem in RPS23B co-localizing the branch point and 3’ splice site (Fig. 2B), and high confidence stems are identified in the secondary structure proposed in RPS14B to bind the RPS14B protein (Fig. 2C). Finally, high confidence stems are predicted in the RPS9A intron structure proposed to regulate gene expression levels of RPS9A and RPS9B (Fig. 2E). In other cases, proposed structures are not supported by the probing data, with DMS accessible residues present in these structures’ stems. These lower support structures include the zipper stem in RPS17B (Fig. S7A), and the RPL30 structure proposed to regulate gene expression (Fig. S7B). The structure in RPS22B found to trigger pre-mRNA degradation was not assessed, as this intron did not pass our DMS-MaPseq coverage threshold. With most structures in this class showing medium or high DMS support, proposed structures from functional experiments assessing mutants and compensatory mutants are in general validated by our data (Fig. 2G).

We next evaluated structures identified by Hooks, et al. (2016)^12^ through predictions from CMfinder^13^, RNAz^14^, and Evofold^15^, approaches that use sequence alignments to identify structures. These structures showed mixed support from DMS data, with introns in YRA1, RPL18B, RPL7A, and RPL28 including some stems with high confidence and other stems with low DMS support (Fig. S8A-D). The intron structures in RPL22B and GLC7 were not predicted from the DMS data (Fig. S8E-F). Proposed intron structures in MPT5 and RPS22B could not be evaluated due to low sequencing coverage. Unlike R-scape, RNAz uses thermodynamics calculations and conservation alone rather than covariation, and CMfinder and Evofold do not use phylogeny sequence backgrounds to identify significant covariation. With a higher fraction of predicted structures that were not validated by DMS data from this set (Fig. 2G), these approaches appear to less reliably identify structures that form *in vivo* compared to functional experiments.

We evaluated intron structures that included covarying residues using DMS data, finding that in the majority of cases, covarying base-pairs were supported by the DMS data. Six of the seven snoRNA-containing introns exhibited sufficient coverage from DMS-MaPseq for evaluation. In these six introns, a majority of covarying residues were supported by DMS data, with 20 of the 27 covarying residues present in high-confidence stems from DMS-guided structure prediction (Fig. S10). Beyond snoRNA-containing introns, covarying residues were identified in four introns from our covariation scan and in two introns from Gao, et al. (2021).^16^ First, we identified covariation in stems of RPL7A’s first intron, and these covarying residues were included in high confidence stems forming a 3-way junction (Fig. 2D). Next, as noted by sequence analysis in Plocik and Guthrie (2012),^9^ we found signals for covariation in a hairpin shared by the introns in RPS9A and RPS9B, and covarying base pairs in these two introns align with the secondary structure from DMS-MaPseq (Fig. 2E-F). Only the structure in RPS13 noted to have covarying residues in Gao, et al. (2021)^16^ was assigned low helix confidence estimates (22%), suggesting that these residues are not paired in the dominant *in vivo* structure (Fig. S7C). The first intron in RPL7B, found to have covarying residues by Gao, et al. (2021)^16^ and in our covariation scan, did not result in enough sequencing coverage from DMS-MaPseq for analysis. Overall, the prevalence of DMS-validated covarying residues across these cases suggests that covariation from R-scape^17^ can reliably identify structures that form *in vivo* (Fig. 2G).

**Intron RNA folding with *in vitro* M2-seq.** Our DMS-guided intron secondary structure predictions suggested that *in vivo*, *S. cerevisiae* introns harbor extended secondary structure, with longer, high-confidence stems compared to coding regions. We explored the structure of introns outside the nuclear environment through *in vitro* structure probing. Here we discuss in detail our findings from *in vitro* probing with M2-seq for a set of *in vitro* transcribed candidate introns.

To assess the *in vitro* folding of individual intron RNAs, we turned to mutate-and-map readout through next-generation sequencing (M2-seq^18^) on introns that were *in vitro* transcribed and probed separate from other RNA. M2-seq allowed us to assess the formation of base-pairs found from DMS-guided secondary structure prediction *in vitro*, identifying base-pairing partners in addition to providing average per-residue accessibility data. For each intron, we generated a pool of RNA *in vitro* transcribed with sparse errors that were installed via error-prone PCR. These mutations can lead RNA molecules to adopt altered secondary structure ensembles, with a mutated residue in a stem exposing its base-pairing partner in solution and potentially leading to unfolding of the stem. Chemical probing of this RNA pool by DMS, in addition to modifying positions that are accessible in the wildtype RNA, also modifies base-pairing partners for mutated stem residues and any other newly accessible positions in the mutated RNA. Off-diagonal signals in the resulting background-subtracted Z-score plots indicate the presence of a stem, as these appear when one residue’s mutation leads to an increase in another residue’s accessibility. In the case of the introns in QCR9 and RPL36B, the Z-score plots from *in vitro* M2-seq included these off-diagonal signals, with mutations in some positions resulting in increased DMS reactivity at other distal positions (Fig. 4A, Fig. S17A). When using RNAstructure to predict secondary structures for these constructs using both 1D and 2D reactivity data, the resulting helix confidence estimates from bootstrapping confirm the formation of stable stems in both cases (Fig. 4C, Fig. S17C). The Z-scores include two-dimensional reactivity signals for many of these stems (Fig. 4A, Fig. S17A), and the resulting base-pairing probabilities support the formation of these stems as these introns’ primary structure (Fig. 4B, Fig. S17B). These secondary structures agree with the stems observed *in vivo*, with high confidence stems *in vivo* also appearing in the *in vitro* M2-seq structures (Fig. 4C-D, Fig. S17C-D).

**VARS-seq experimental design and validation.** To evaluate the effects of intron structure variants with VARS-seq, we integrated genes containing intron libraries into the yeast genome^19^ (Fig. 6C). Reasoning that exogenous reporter genes can alter pre-mRNA secondary structure and splicing patterns, we instead integrated them into the genome in their full native gene context with CRISPR/Cas9 (Fig. S19C). Random barcodes (12N) were installed in the 5′ UTR upstream of the intron to serve two roles. First, these sequences helped link spliced reads to the pre-mRNA variant they originated from. Second, these randomized sequences provided perturbations for the efficiency of splicing and mRNA decay; these perturbations are especially useful to identify effects in cases where wildtype retained intron levels are beyond the dynamic range for our assay. For instance, in the case that retained intron levels are low with highly efficient splicing for a wildtype intron, this barcoding strategy can provide additional dynamic range for our assay because some randomly designed barcode sequences will attenuate splicing or pre-mRNA decay rates. In particular, by measuring the effects of many unique barcode sequences per intron variant, we can observe the distribution of retained intron levels for wildtype and variant sequences across barcodes, robustly identifying variants that shift this distribution and alter gene expression.

We sequenced genomic DNA (gDNA) to link randomized barcodes to intron variant sequences and obtain transformation frequencies, and we measured spliced and unspliced RNA levels with targeted RNA sequencing (Fig. 6C). Reassuringly, most consensus variant sequences obtained from gDNA sequencing were the expected library length (between 71.5% and 84.8% across sub-libraries, Fig. S20A), and most designed variants were assigned to at least 10 unique barcodes (92.1%, Fig. S20B). Our targeted RNA sequencing measurements were free from genomic DNA contamination (Fig. S20C), and as expected, variants that disrupted key splice site sequences (branch point mutants in QCR9) significantly increased retain intron levels (Fig. S20D). Though we did observe alternative splicing events from our RNA sequencing data, all 6 observed events represented a minor population of transcripts produced from variants including these events (Fig. S20E). Using our DNA and RNA sequencing data, we computed the following two metrics for each barcode and variant sequence: the retained intron (RI) fraction (fraction of RNA-seq reads that were unspliced) and the normalized mRNA level (spliced mRNA read counts normalized by the gDNA read counts). Though many variants led to small changes in these two readouts compared to wildtype sequences, barcode sequences enabled us to robustly identify even subtle effects by providing perturbations that altered baseline splicing and decay rates.

**Comparing intron DMS-guided structure prediction with *de novo* structure prediction.** We found that many structural features that were enriched in introns when using DMS-guided structure prediction remained enriched when using *de novo* structure prediction. Zipper stems and downstream stems in introns are more stable (lower dG values) than those in control sequences, whether using DMS-guided structure prediction, *de novo* minimum-free energy structure prediction, or *de novo* secondary structure ensemble prediction (p-values < 0.01, blue and orange in Fig. 7B). Introns have longer stems than length-matched control sequences and have higher maximum extrusion from ends, again both from *de novo* structure predictions and DMS-guided structure predictions (p-values < 0.01, green and red in Fig. 7B). We additionally measured the secondary structure graph distance between the 5’ splice site and branch point sequence positions, comparing between introns and the control sequences (purple in Fig. 7B). From DMS-guided structures, the 5’ splice site and branch point were more distant in introns than controls (p-value < 0.01), perhaps due to the requirement for single-stranded nucleotides proximal to these sequences in the A-complex spliceosome. On the other hand, *de novo* structure prediction suggests that introns have lower secondary structure graph distances between the 5’ splice site and branch point without the context of the spliceosome (p-values < 0.01), as found by Rogic, et al. (2008),^5^ suggesting that introns co-localize key splicing sequences. We tested other structure prediction approaches and found similar structural feature enrichment, comparing against other control sets, using an alternate folding package, and making predictions including exonic context (Fig. S25). In particular, the enrichment for stable zipper stems, higher maximum extrusion from ends, and lower 5’ splice site to branch point distances remained significant across all tested *de novo* prediction approaches.

Supplemental Figures

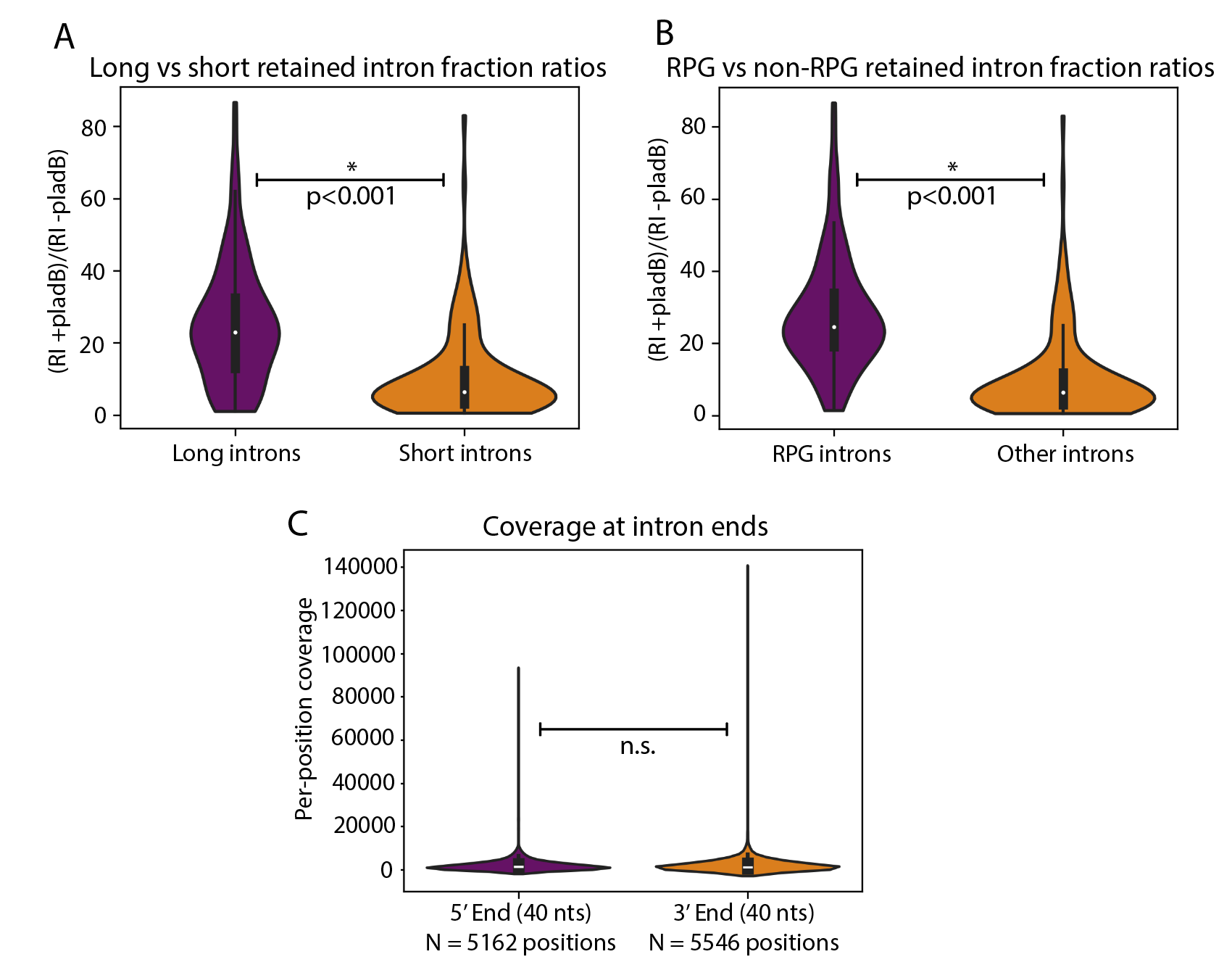
**Figure S1**: Analyzing effects of pladB treatment on intron retention and intron degradation. Comparing the ratio of the retained intron (RI) fraction between A) long (> 200 nucleotides) and short (< 200 nucleotides) introns, and between B) introns in ribosomal protein coding genes (RPGs) vs other introns. C) Comparing per-position coverage at the 5’ end and 3’ end of all introns after pladB treatment (no significant difference at p=0.05 threshold). p-values were computed using Wilcoxon ranked sum tests to compare classes.

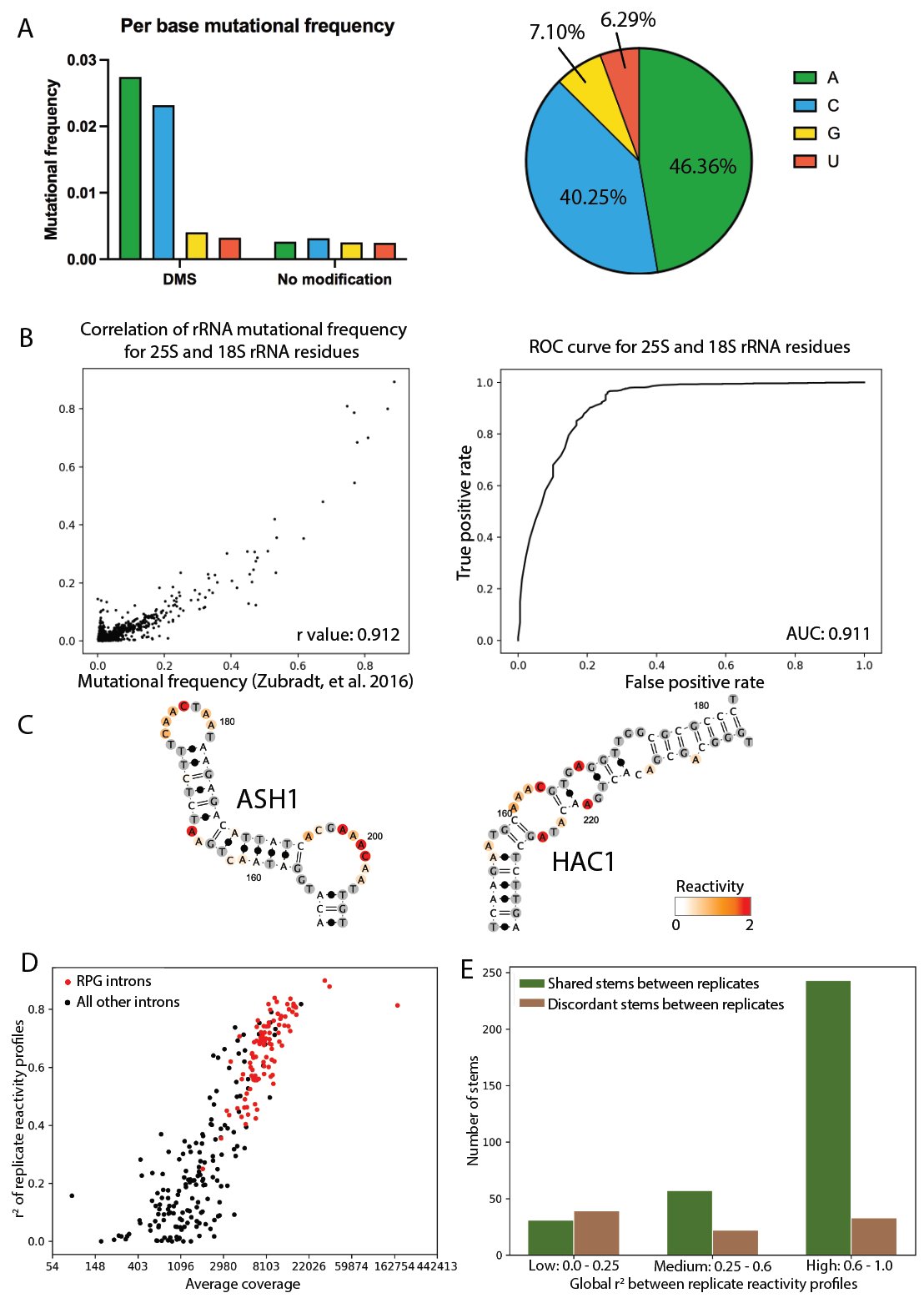

**Figure S2**: DMS-MaPseq data quality. A) Per-base mutational frequencies. B) Accuracy of mutational frequency values for rRNA residues. C) HAC1 and ASH1 positive control structures with overlaid reactivity profiles.

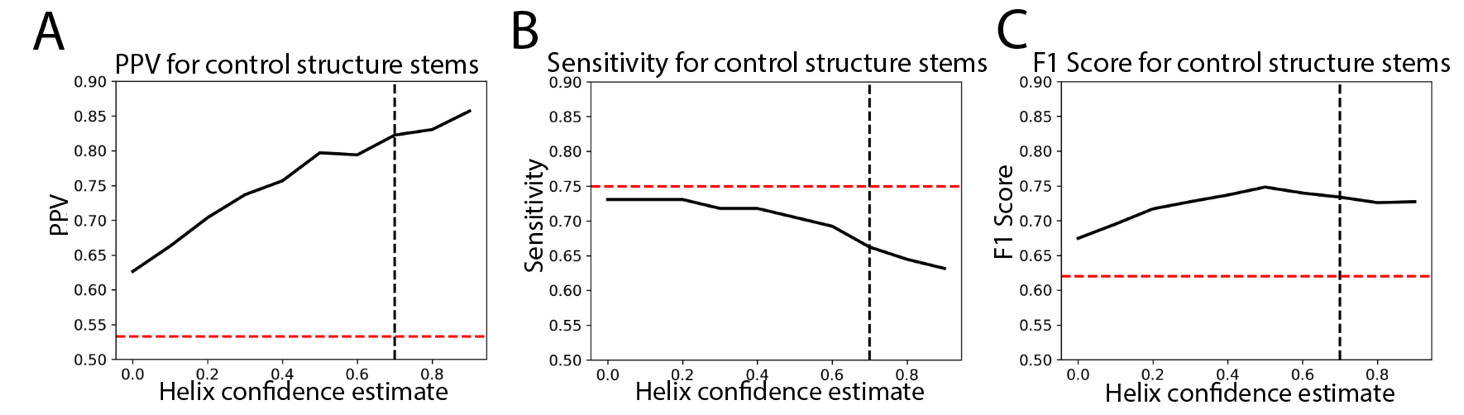

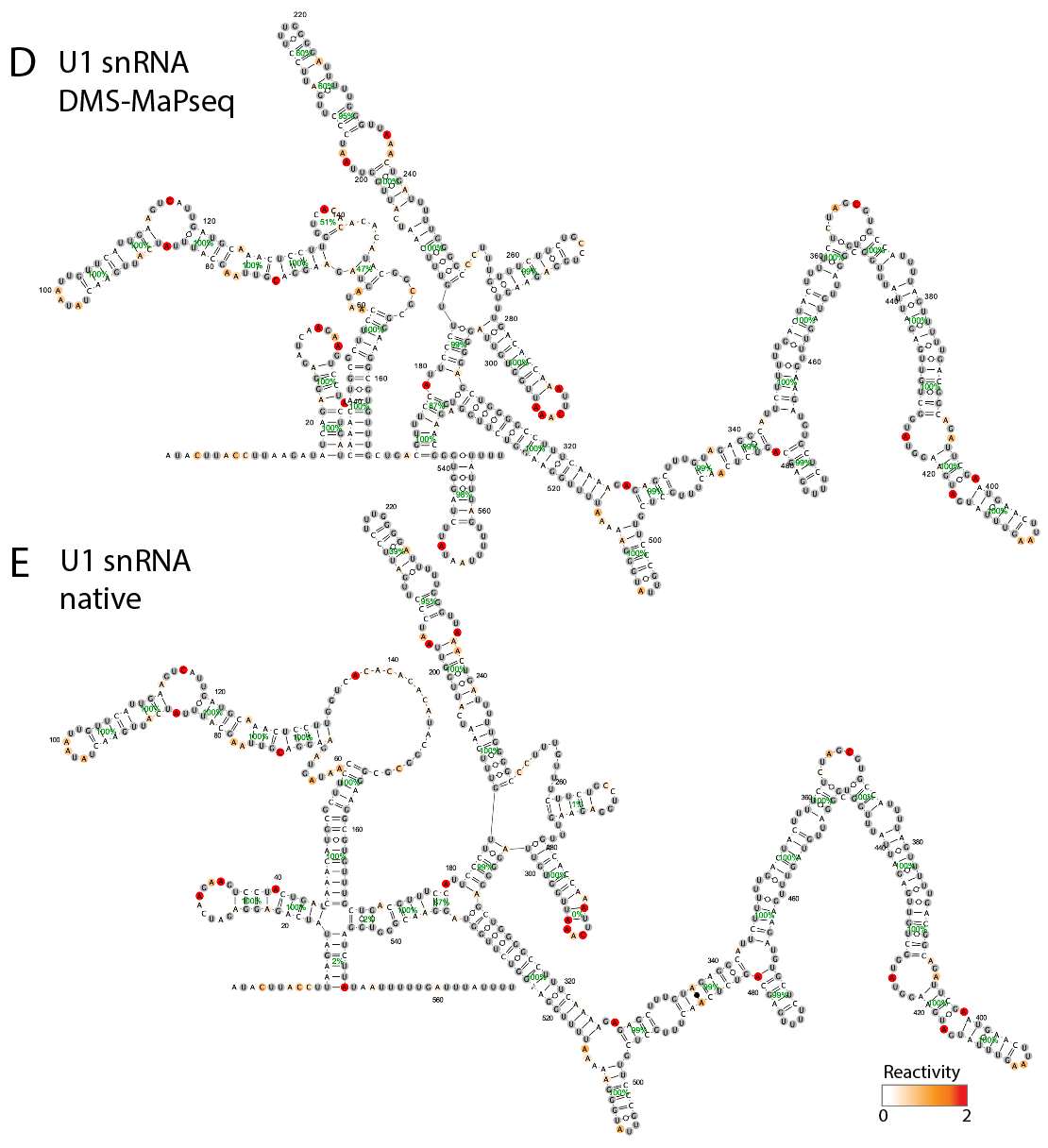

**Figure S3**: Support from DMS reactivity for *in vivo* formation of control structures. A) PPV, B) sensitivity, and C) F1 score for structure prediction of a set of control RNA structures (rRNAs, tRNAs, snRNAs, and mRNAs; Table S2), using RNAstructure guided by DMS with varying helix confidence estimate cutoffs for calling stems. The black dotted line represents the helix confidence estimate 0.7 chosen in this paper. The red dotted line represents the PPV, sensitivity, and F1 score for Vienna RNA structure prediction without using DMS data. D)-E) DMS-MaPseq structure prediction for the U1 snRNA compared to the native secondary structure obtained from Li, et al. (2017).^20^ D) DMS-guided secondary structure prediction for the U1 snRNA, with reactivity values overlaid and helix confidence estimates indicated in green percentages. E) Native secondary structure for the U1 snRNA, with DMS reactivity overlaid along with helix confidence estimates. For the native structure, helix confidence estimates were computed as the percent of bootstrapping iterations where the helix was recovered when sampling DMS reactivity values and making structure predictions.

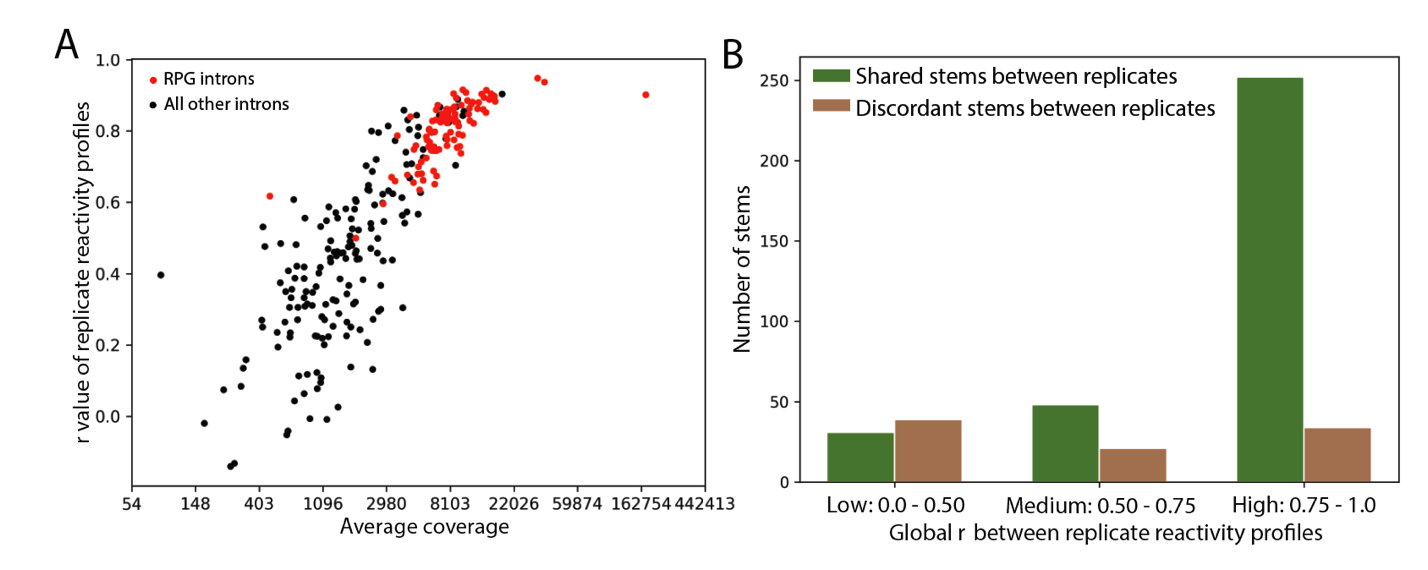

**Figure S4**: DMS-MaPseq reproducibility between replicates. A) Pearson’s correlation coefficient r between replicates for each intron in *S. cerevisiae* versus the average sequencing coverage between replicates. B) Bar graph comparing the number of stems that are shared vs discordant between replicates for introns in different correlation ranges.

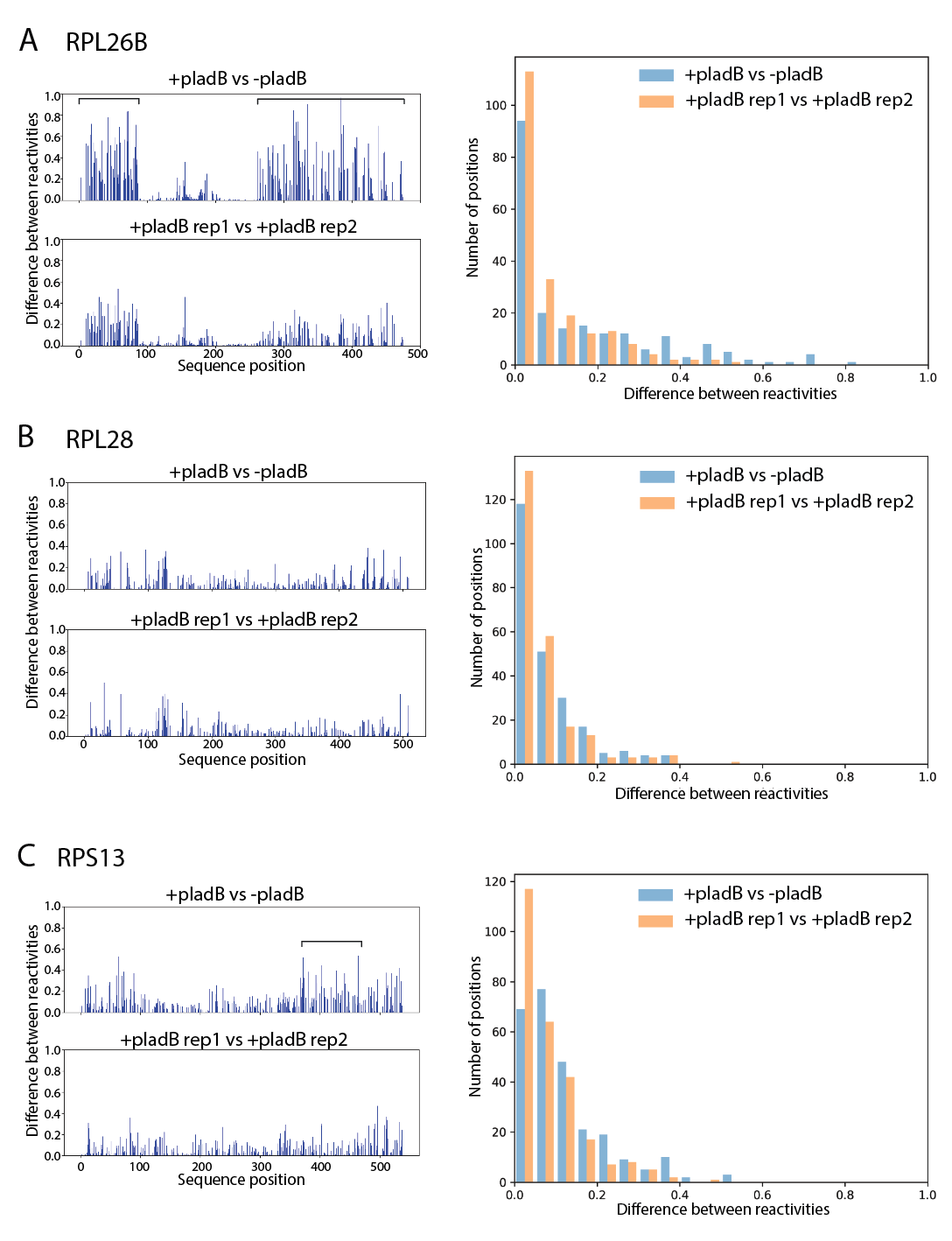

**Figure S5:** Comparing reactivity values with and without pladB treatment for A) RPL26B, B) RPL28, and C) RPS13. Left: the absolute value of the difference between reactivities (top row: comparing with and without pladB treatment; bottom row: comparing two replicates with pladB treatment.) Brackets highlight intervals including at least one position with > 0.5 absolute difference between reactivity values with vs without pladB treatment. Right: histograms summarizing the number of sequence positions with specified absolute differences between reactivity values.

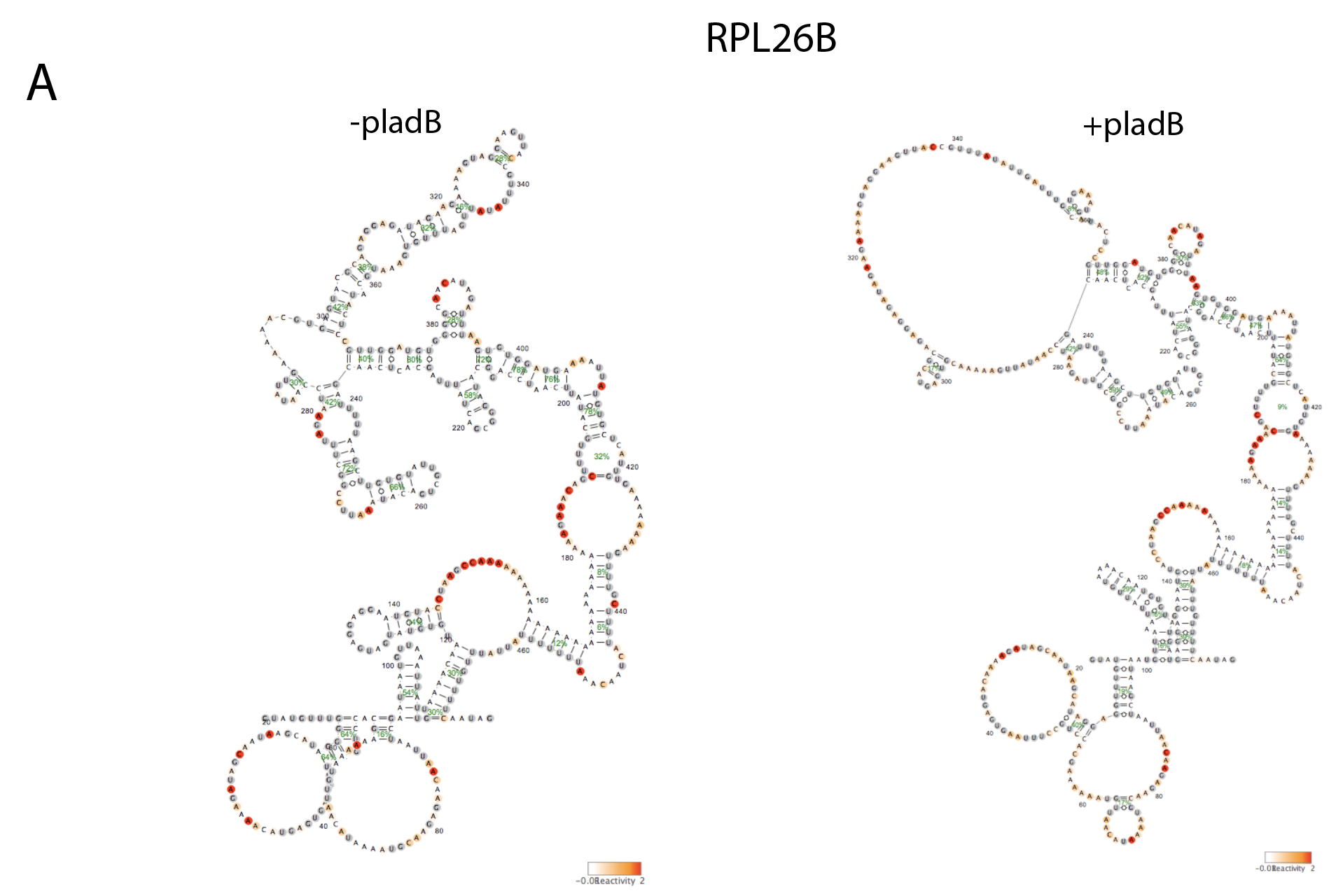

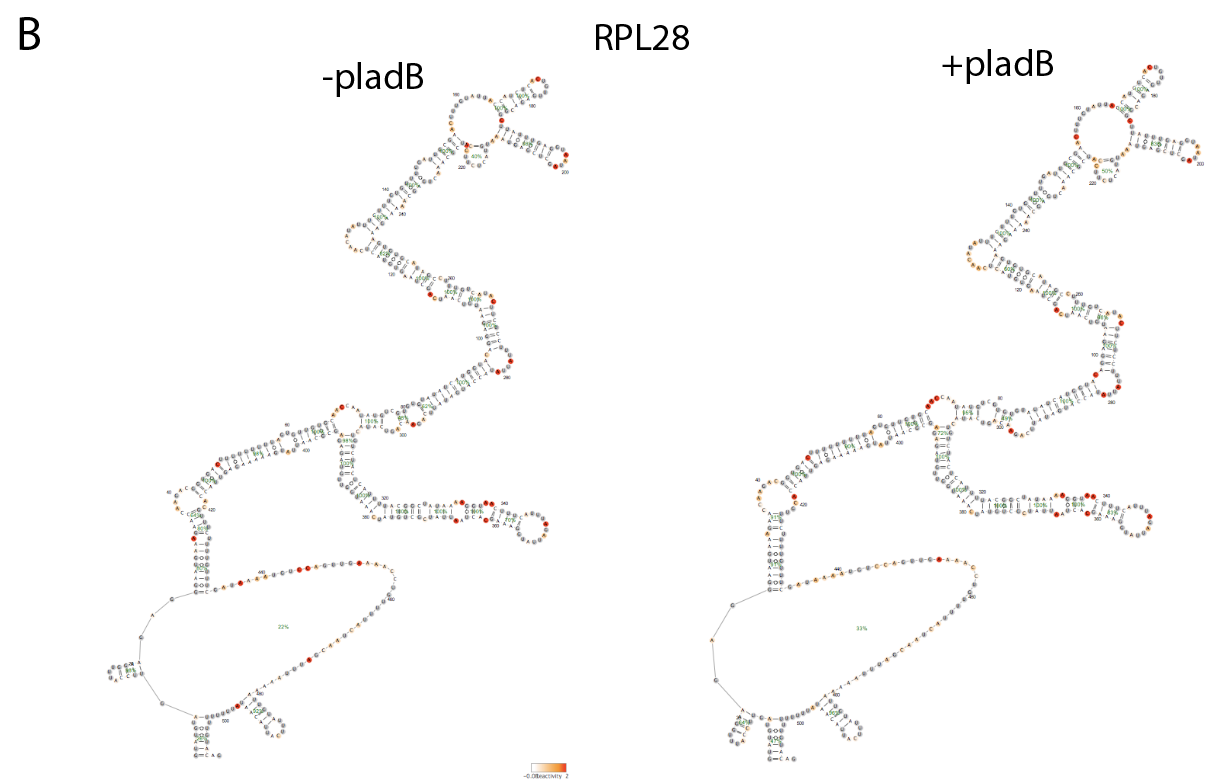

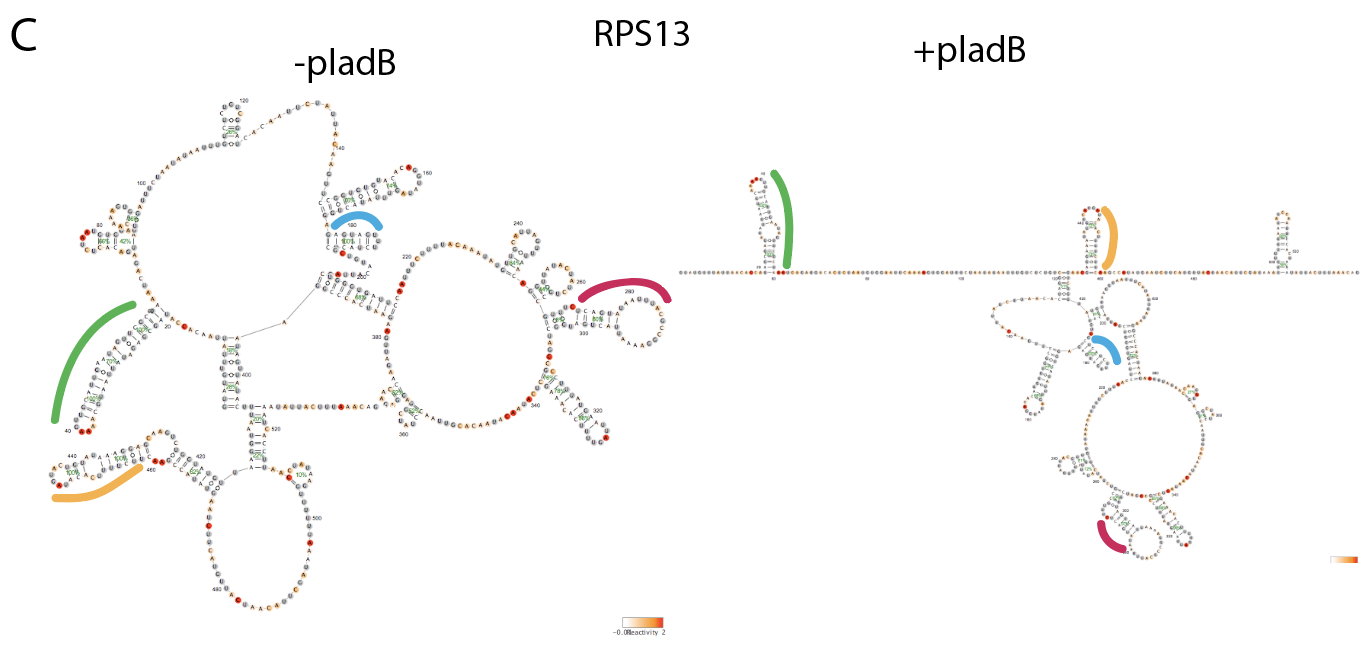

**Figure S6**: Reactivity values, structure prediction, and bootstrapping probabilities with and without pladB treatment for: A) RPL26B, B) RPL28, and C) RPS13. Common stems are highlighted in matching colors for RPS13.

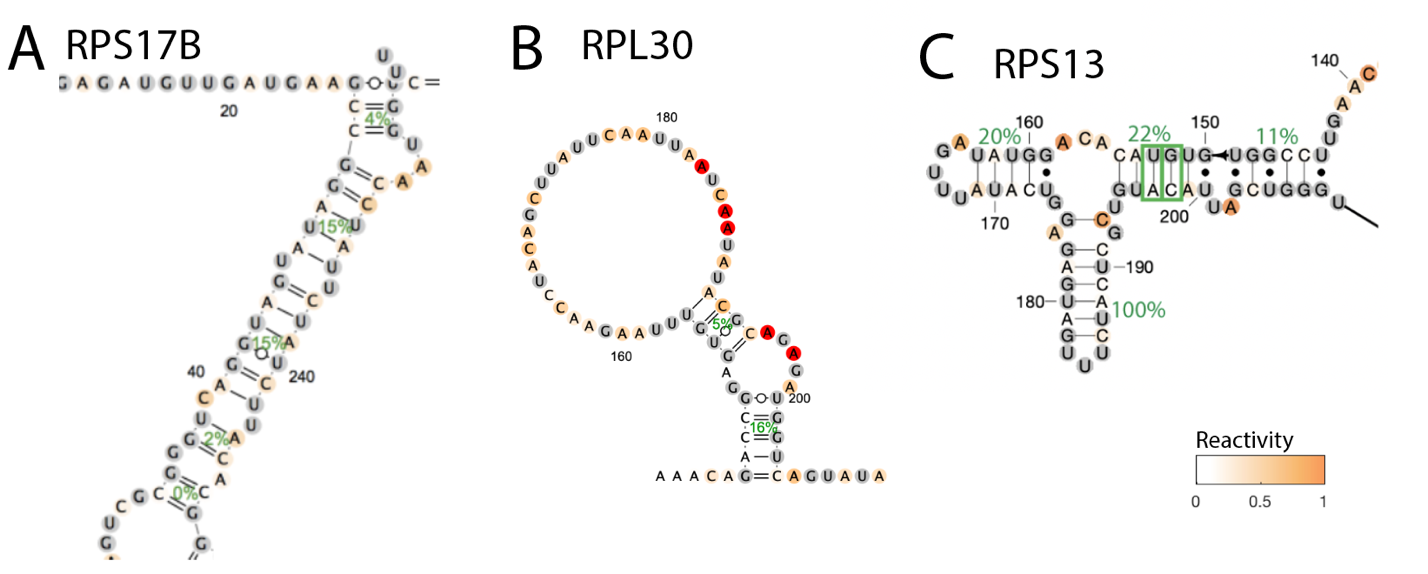

**Figure S7:** Structures previously proposed from functional experiments or covariation scans with low DMS reactivity support. Secondary structures are colored by DMS reactivity and helix confidence estimates are depicted as green percentages. Covarying base pairs in RPS13 are indicated as green boxes.

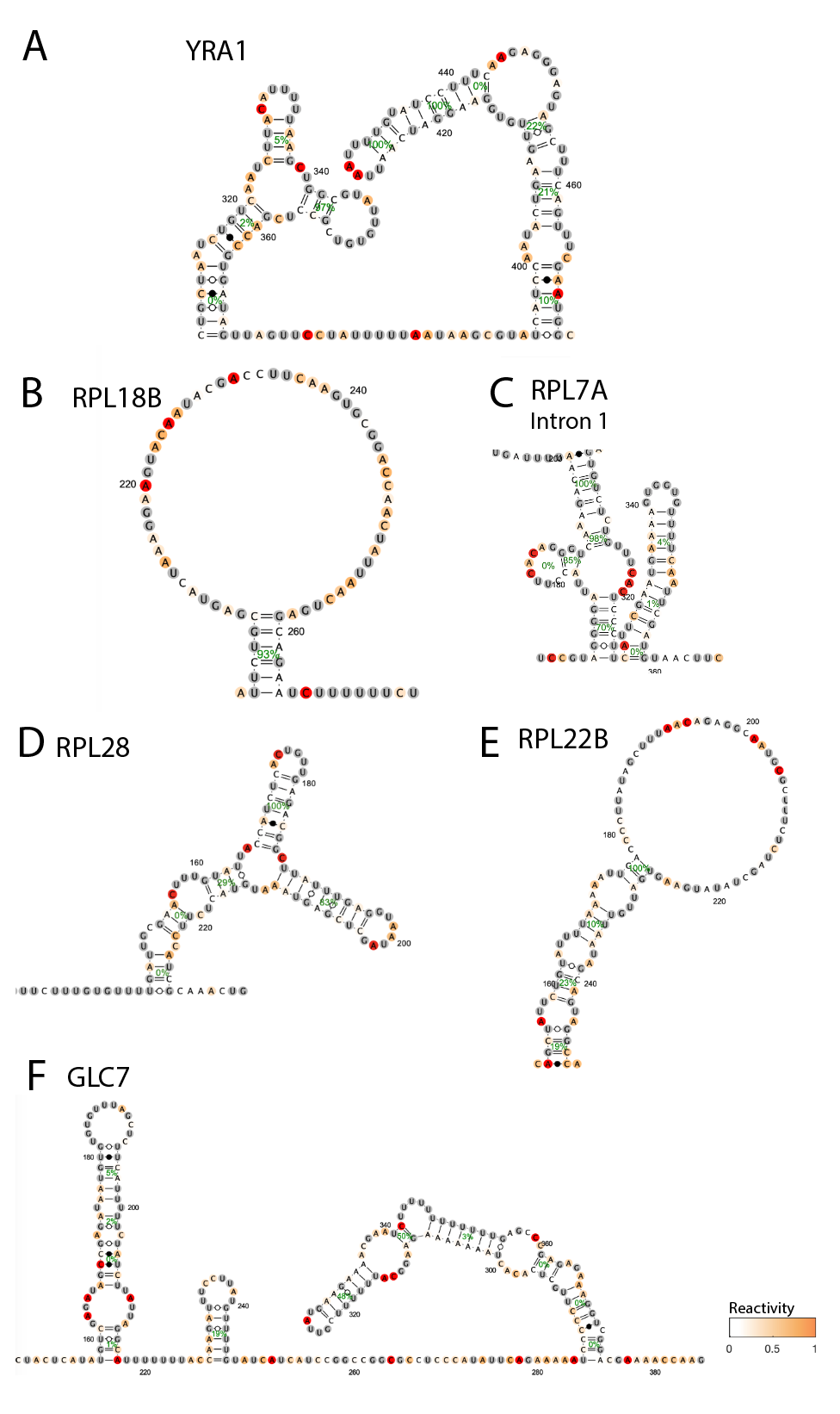

**Figure S8:** DMS reactivity support for *in vivo* formation of structures from computational prediction based on evolutionary sequence alignments.

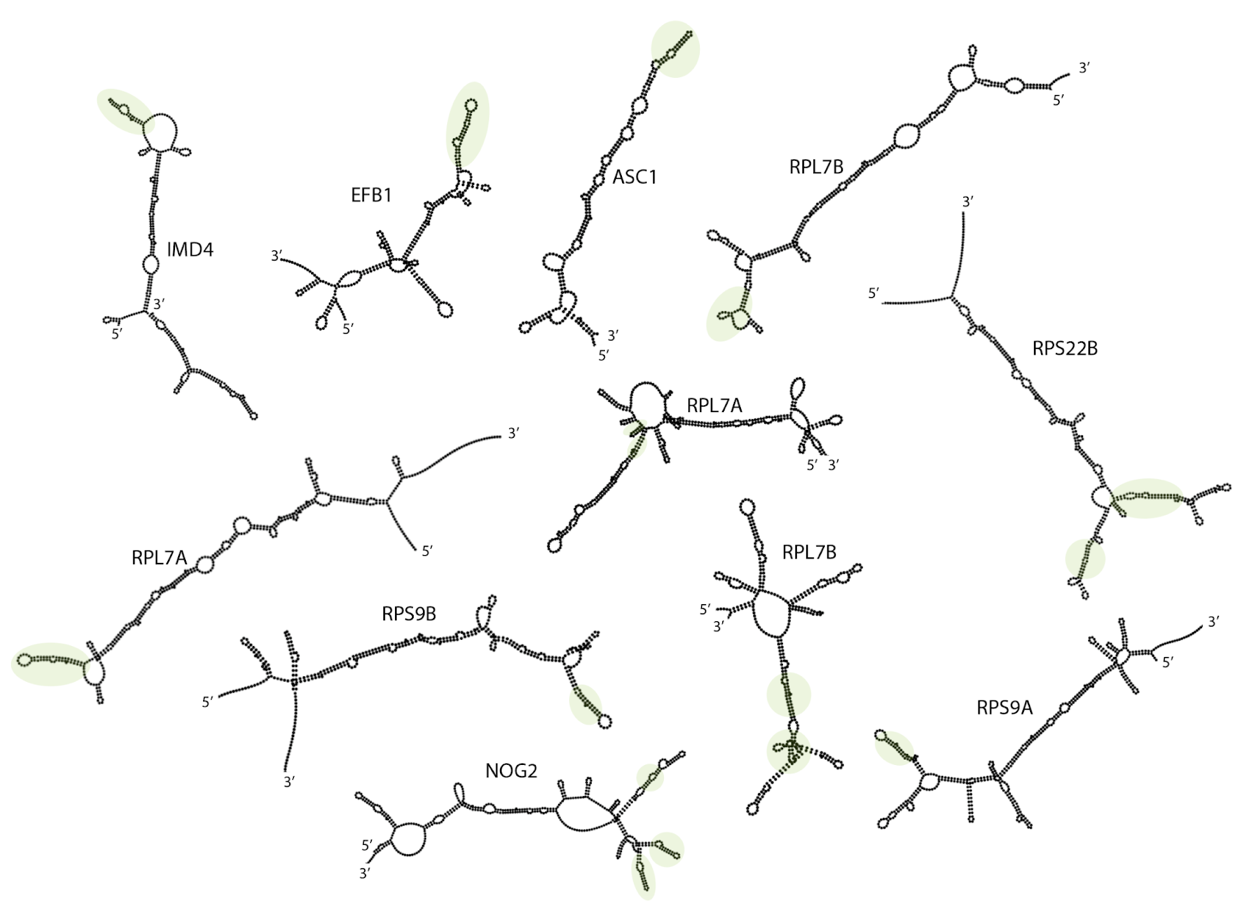

**Figure S9**: Full gallery of introns longer than 200 nucleotides with at least one covarying base-pair in a stem of at least 3 base-pairs, only showing cases where the covariation data was consistent with the predicted minimum free energy structure.

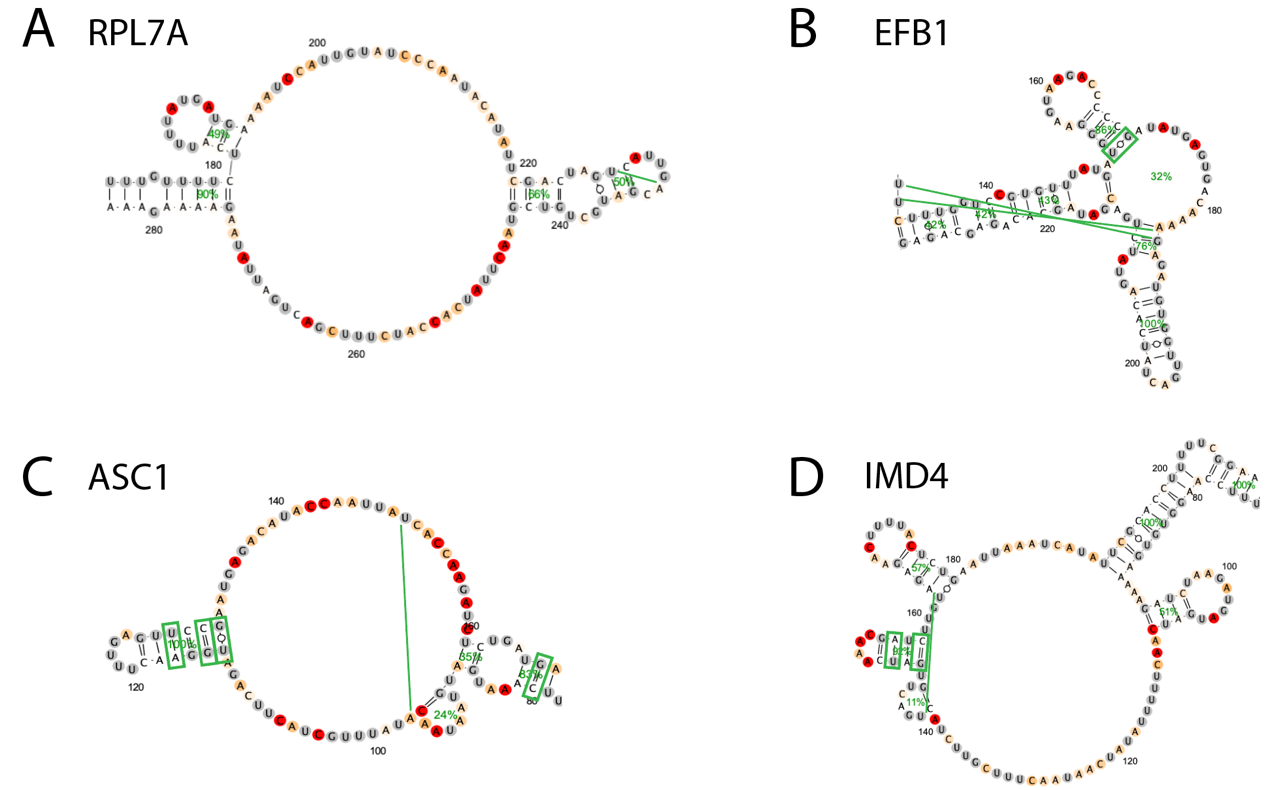

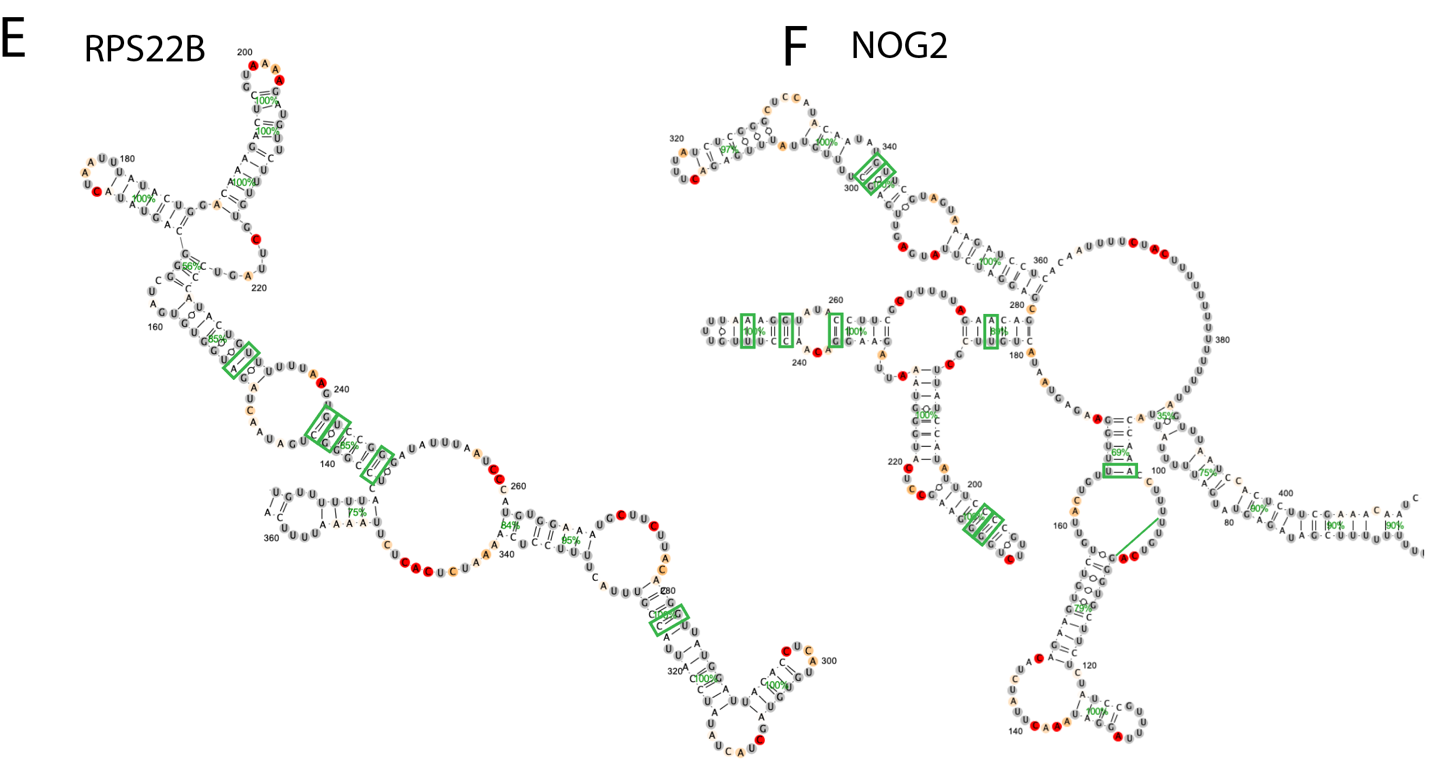

**Figure S10**: DMS reactivity support for the *in vivo* formation of intron structures that include covariation in snoRNA regions. Covarying base-pairs are annotated in green boxes when they agree with base-pairs from the DMS-guided structure prediction, and lines when they include residues that are not base-paired in the DMS-guided structure prediction.

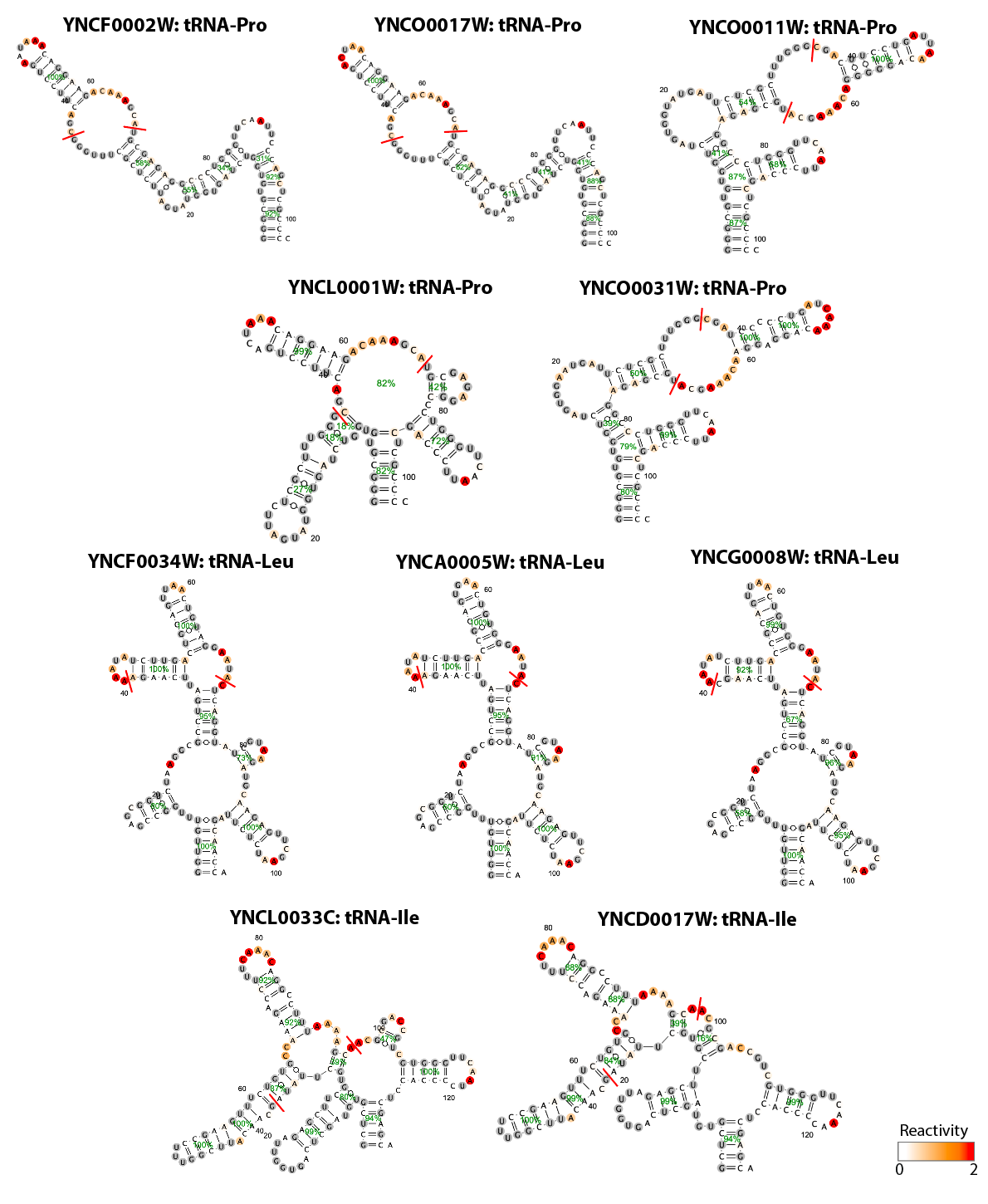

**Figure S11:** DMS-guided structure predictions for tRNAs containing introns with length at least 30 nucleotides. Red lines indicate the excised introns’ start and end positions.

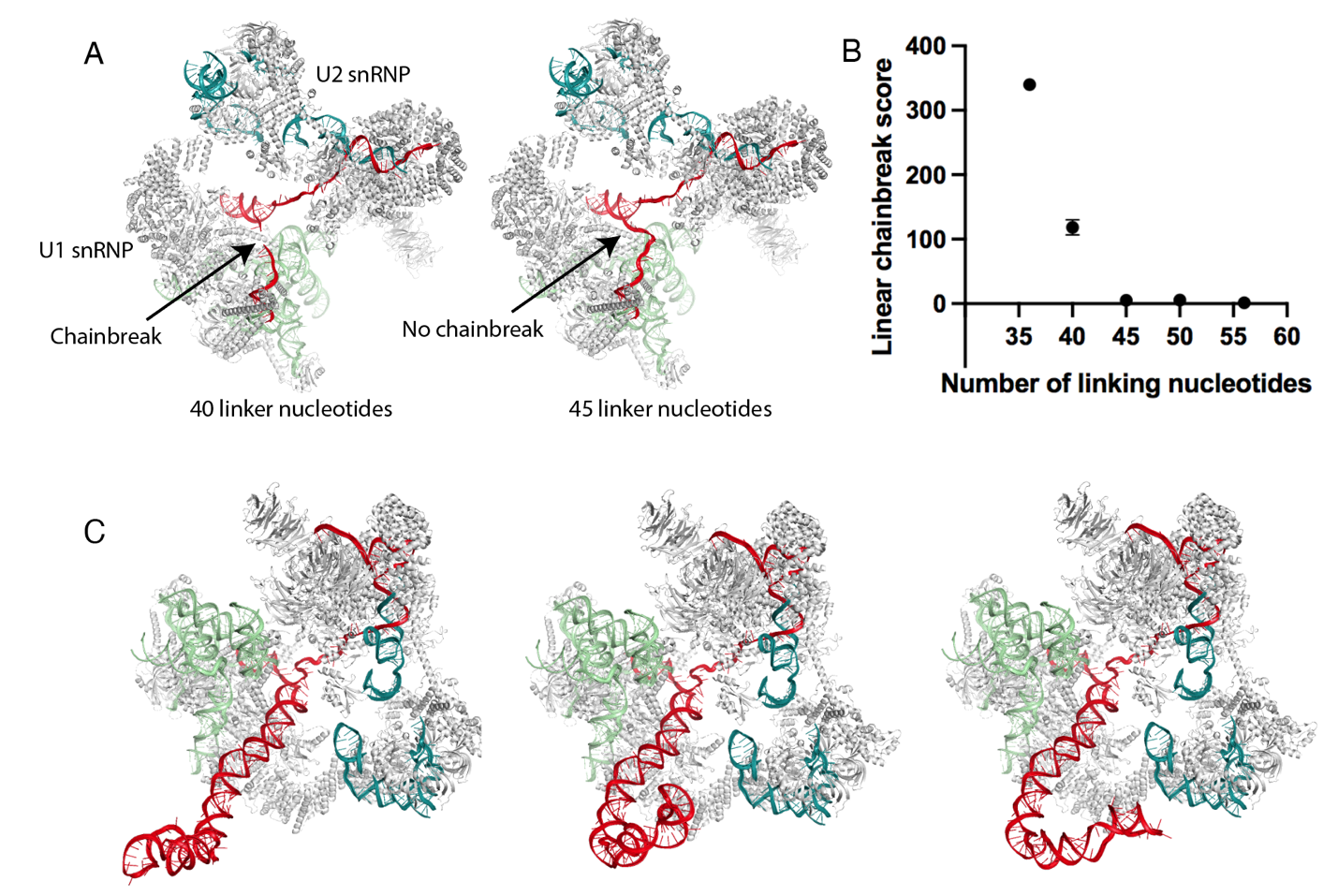

**Figure S12**: Structural modeling with the A state spliceosome. A) Sample Rosetta models of introns with varying linker lengths to identify linker lengths for zipper stems compatible with the A state spliceosome structure. B) Penalty for chain breaks as modeled linker length increases. C) Top 3 models for RPL36B intron in the context of the A state spliceosome, with the RPL36B intron secondary structure specified from DMS-MaPseq.

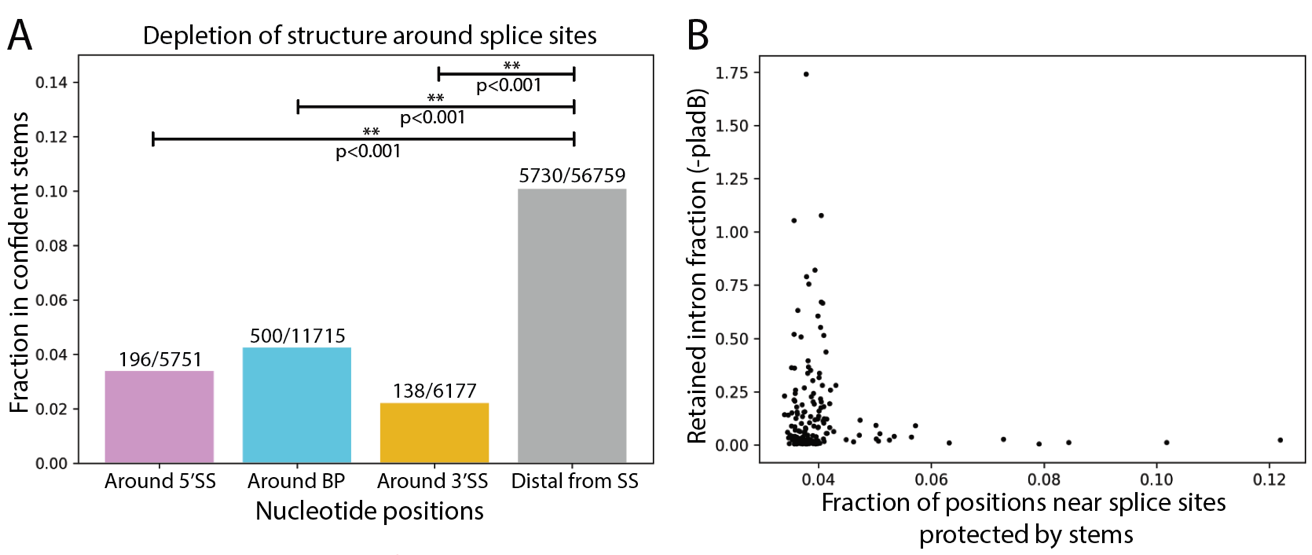

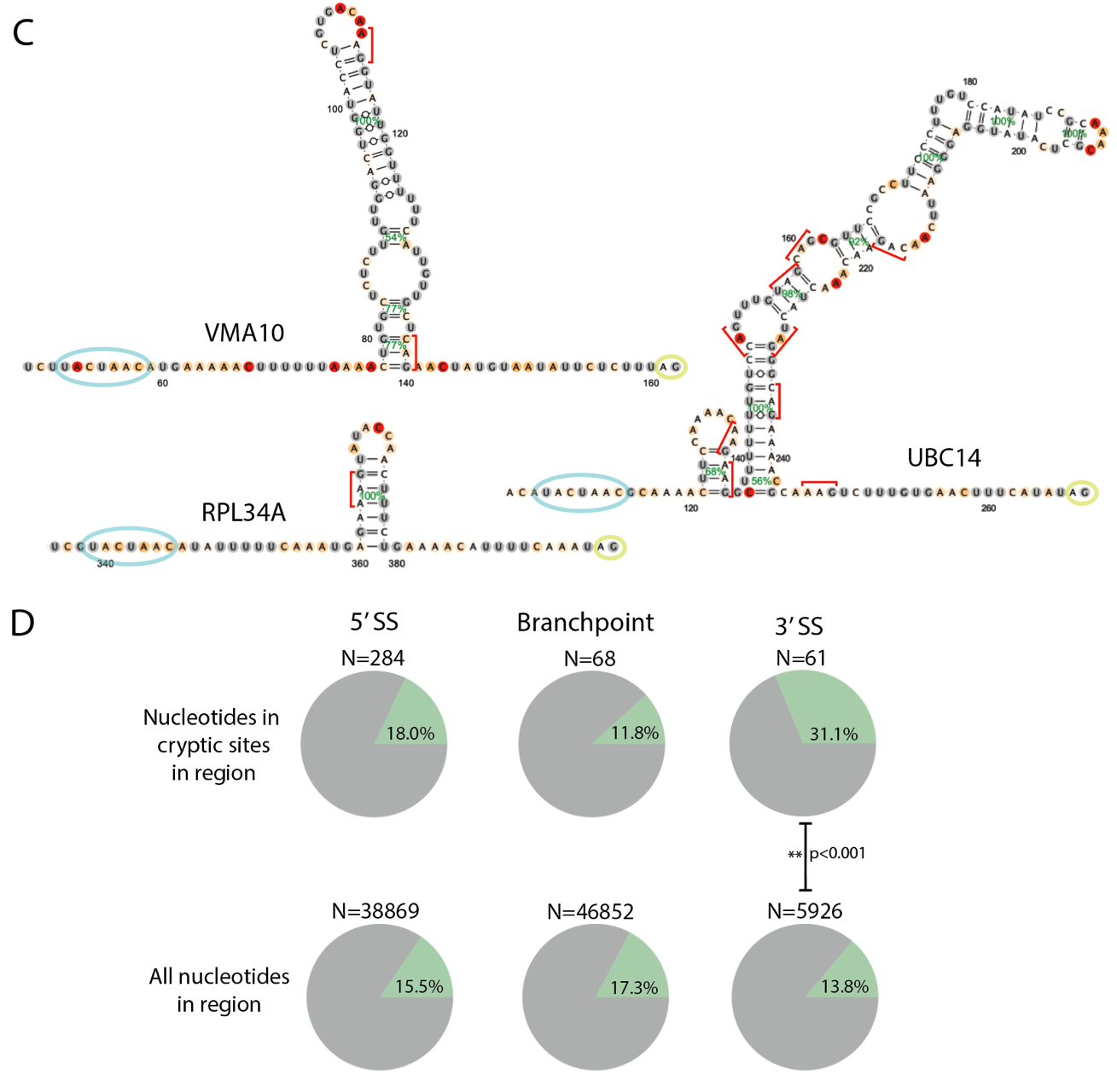

**Figure S13:** Intron structures surrounding canonical and cryptic splice sites. A) Proportion of nucleotides in high confidence stems from sequence intervals surrounding the canonical 5’ splice site, branch point, and 3’ splice site sequences across all introns. Intervals around 5’ splice site, branch point, and 3’ splice site sequences were identified by inspecting spliceosome structures and finding sequence regions that thread through the spliceosome (details in Methods). All intron sequence positions external to these intervals were included in the “distal from SS” set. High confidence stems were identified using structure predictions for introns with 50 nucleotides of surrounding pre-mRNA sequence context. P-values are computed with Chi-squared tests on 2x2 contingency tables. B) The relationship between retained intron fraction and the fraction of positions surrounding the 5’ splice site, branch point, and 3’ splice site sequences that are occluded by high confidence stems. C) Example structures with downstream stems between the branch point (blue circle) and 3’ splice site (yellow circle) that occlude cryptic 3’ splice sites (red brackets). Secondary structures are colored by DMS reactivity and helix confidence estimates are depicted as green percentages. D) Comparison of the proportion of nucleotides protected by high confidence stems between nucleotides in cryptic splice sites versus nucleotides surrounding these sequences. Cryptic splice sites were identified by searching for sequences that matched other introns’ splice site sequences in defined sequence intervals (details in Methods.) P-values are computed with Chi-squared tests on 2x2 contingency tables.

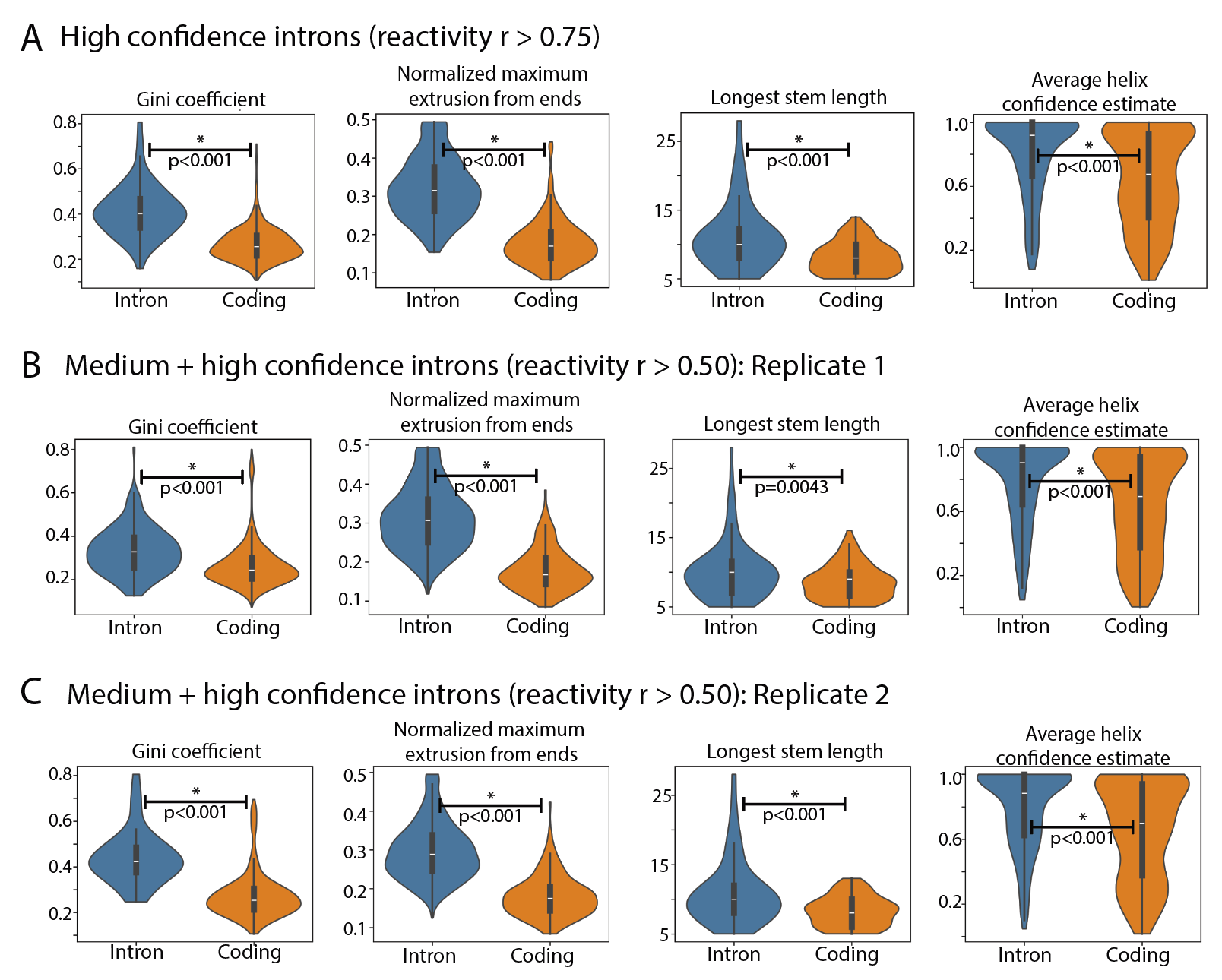

**Figure S14:** Comparing intron and coding structural properties across replicate experiments. Comparisons are shown for A) high confidence introns (between-replicate reactivity r > 0.75) combining data from both replicate DMS-MaPseq experiments, B) medium and high confidence introns (between-replicate reactivity r > 0.5) using data from the first DMS-MaPseq replicate, and C) medium and high confidence introns (between-replicate reactivity r > 0.5) using data from the second DMS-MaPseq replicate. P-values for comparisons of secondary structure features between introns and coding regions were computed using Wilcoxon ranked sum tests.

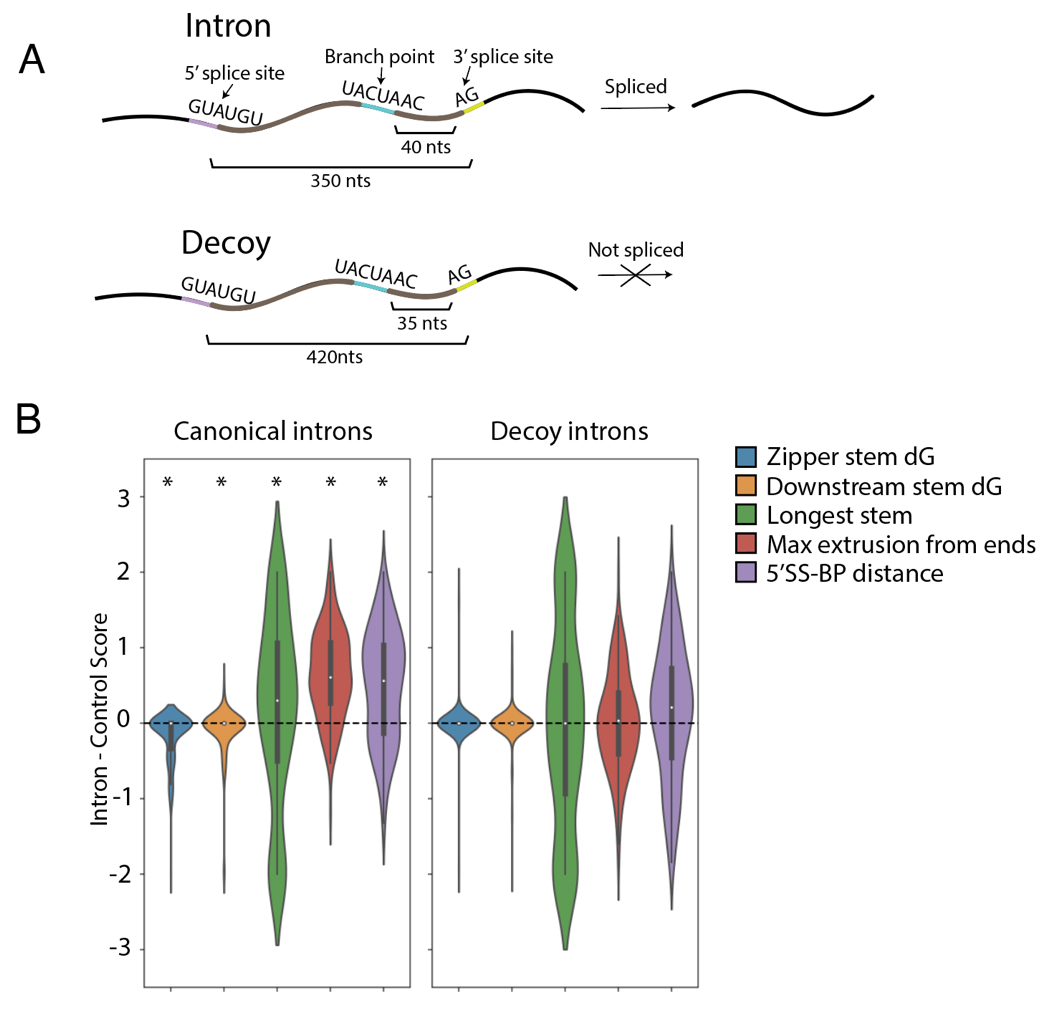

**Figure S15**: Secondary structure features for introns and non-splicing decoy sequences. A) Intron and decoy sequence schematic. Nucleotide lengths depicted are representative lengths for introns and decoy sequences, with decoy sequences chosen to have 5’ splice site, branch point, and 3’ splice site placements matching length distributions from canonical introns (see Methods). B) Secondary structure features are enriched in standard canonical spliced introns in yeast, but not in decoy sequences (genomic intervals which match splice site sequences and yet do not splice). *p-value < 0.01 by Wilcoxon ranked-sum test.

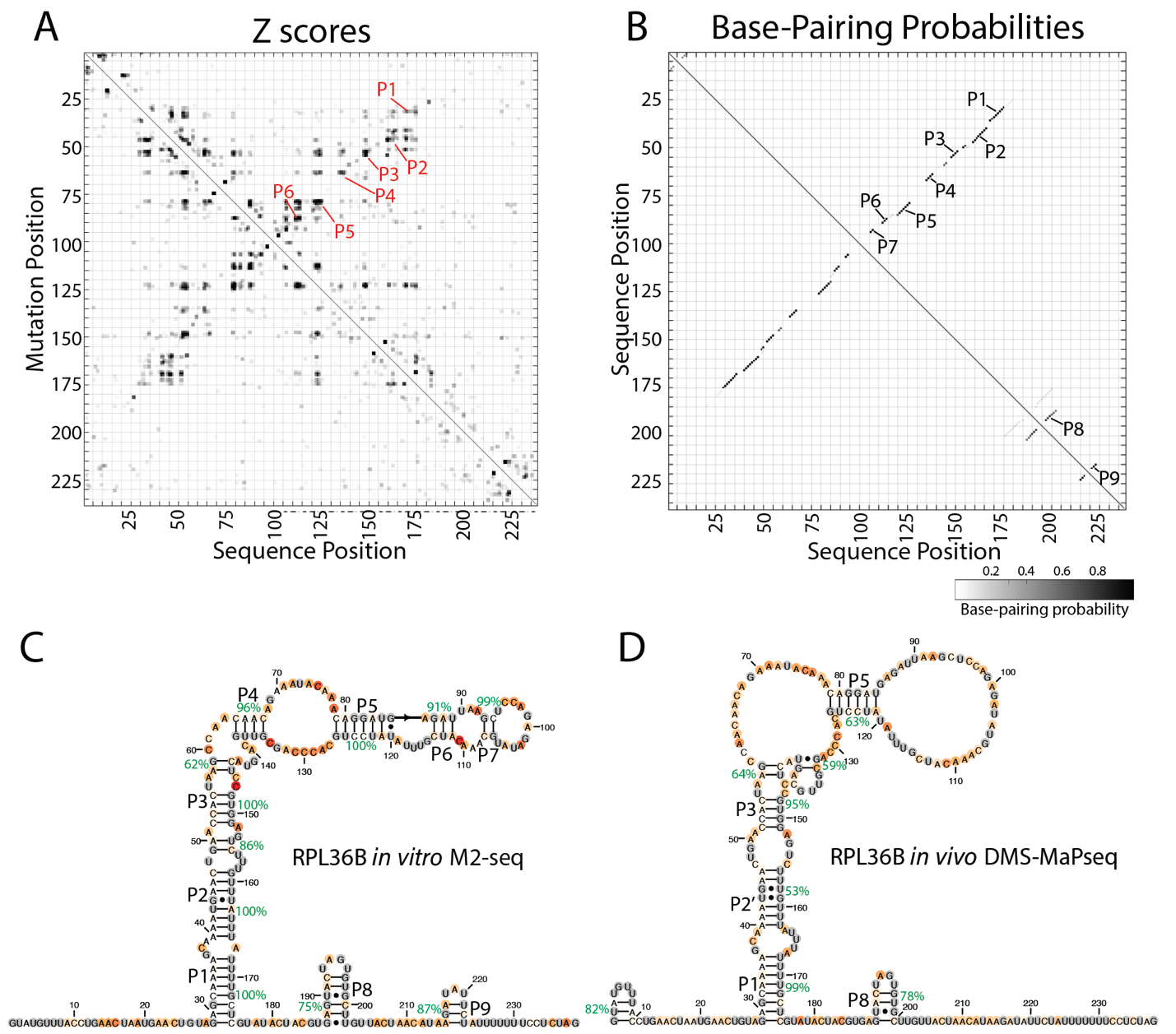

**Figure S16:** Multidimensional chemical mapping for RPL36B. A) *In vitro* M2-seq Z scores for the intron in RPL36B, with peaks representing helices annotated in red. B) *In vitro* chemical reactivity base-pairing probabilities for RPL36B using 1D and 2D chemical reactivity from M2-seq. Secondary structure predictions guided by 1D and 2D DMS probing data for the intron in RPL36B C) from *in vitro* M2-seq, and D) from *in vivo* DMS-MaPseq.

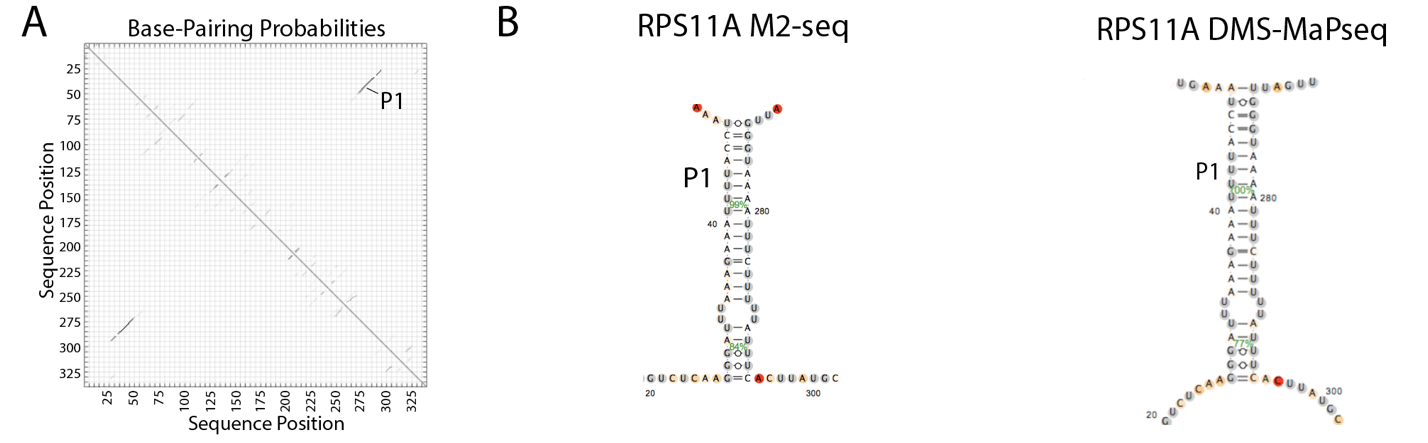

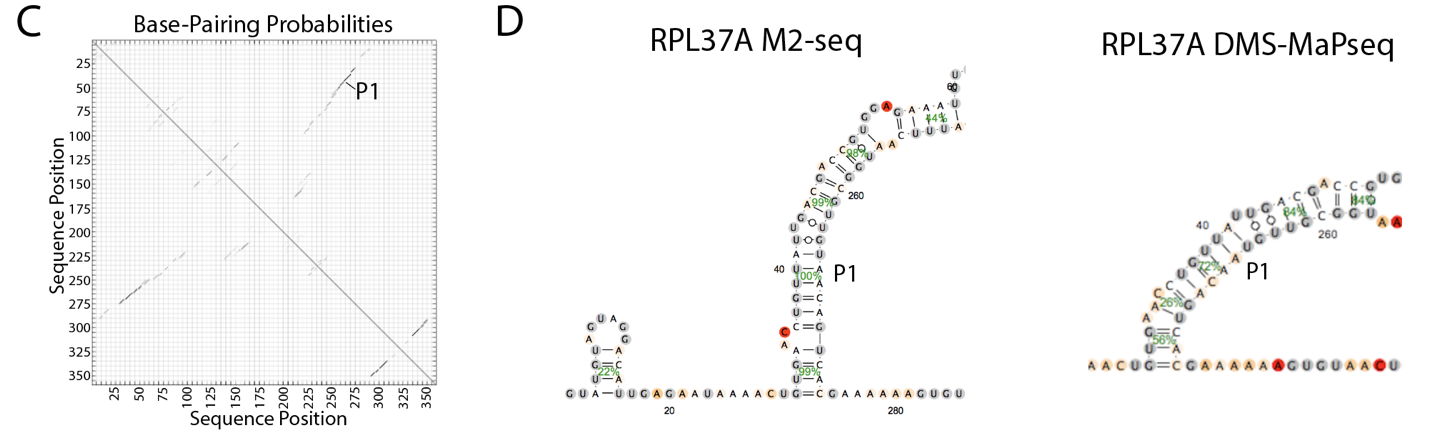
repl
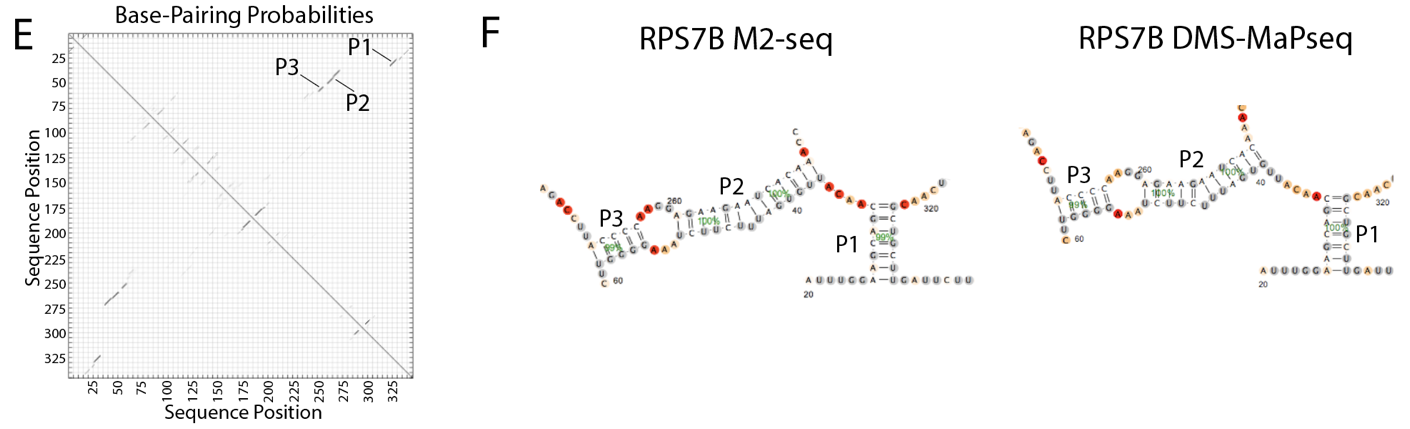

**Figure S17**: Base-pairing probabilities and secondary structure predictions from *in vitro* M2-seq and *in vivo* DMS-MaPseq for RPS11A (A, B), RPL37A (C, D), RPS7B (E, F).

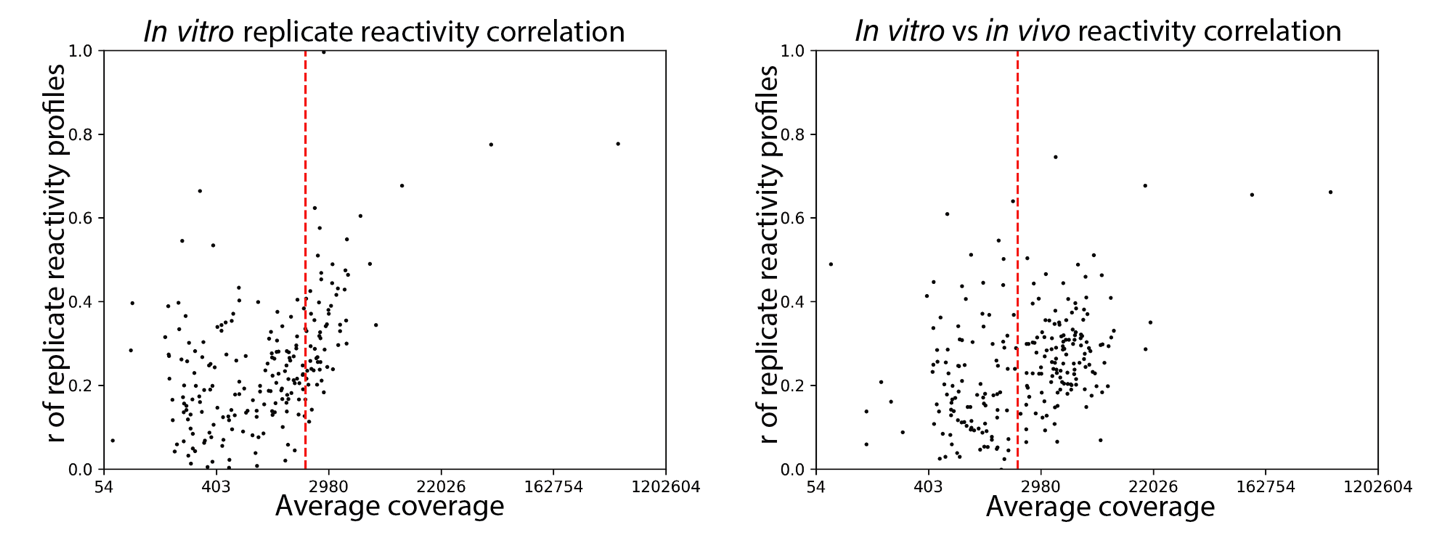

**Figure S18**: Probing *in vitro* refolded RNA. Left: Pearson’s correlation coefficient between *in vitro* DMS-MaPseq replicates for each intron in *S. cerevisiae* versus the average sequencing coverage between replicates. Right: correlation between reactivity values for introns probed in *in vivo* and *in vitro* DMS-MaPseq, versus the average coverage in these experiments. The vertical red line indicates the coverage cutoff used for analysis of *in vivo* samples.

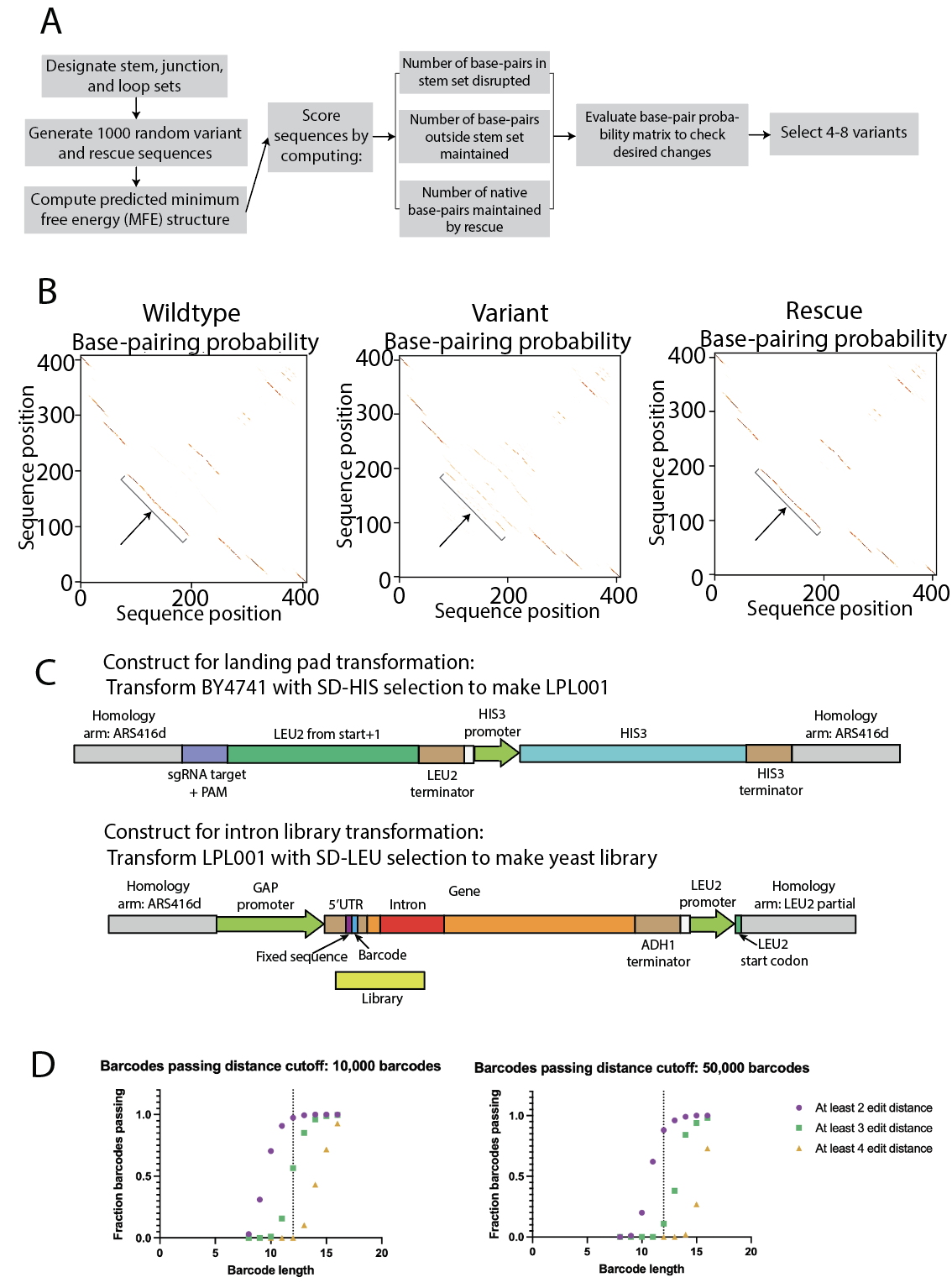

**Figure S19**: Structure variant library design to measure spliced and unspliced RNA levels for intron variants. A) Computational pipeline for designing variant and rescue sequences to assess stem and loop sets in an intron. B) Sample base-pair probability matrix comparison between a set of wildtype, variant, and rescue sequences. The stem set targeted by this variant and rescue sequence are bracketed. C) Constructs used for constructing the landing pad strain LPL001 (top) and for transforming the intron library via genomic integration into LPL001 (bottom). D) Based on simulations sampling 10,000 or 50,000 random barcodes, the number of barcodes within 2, 3, or 4 edit distance of another barcode. The vertical line indicates the chosen barcode length (12).

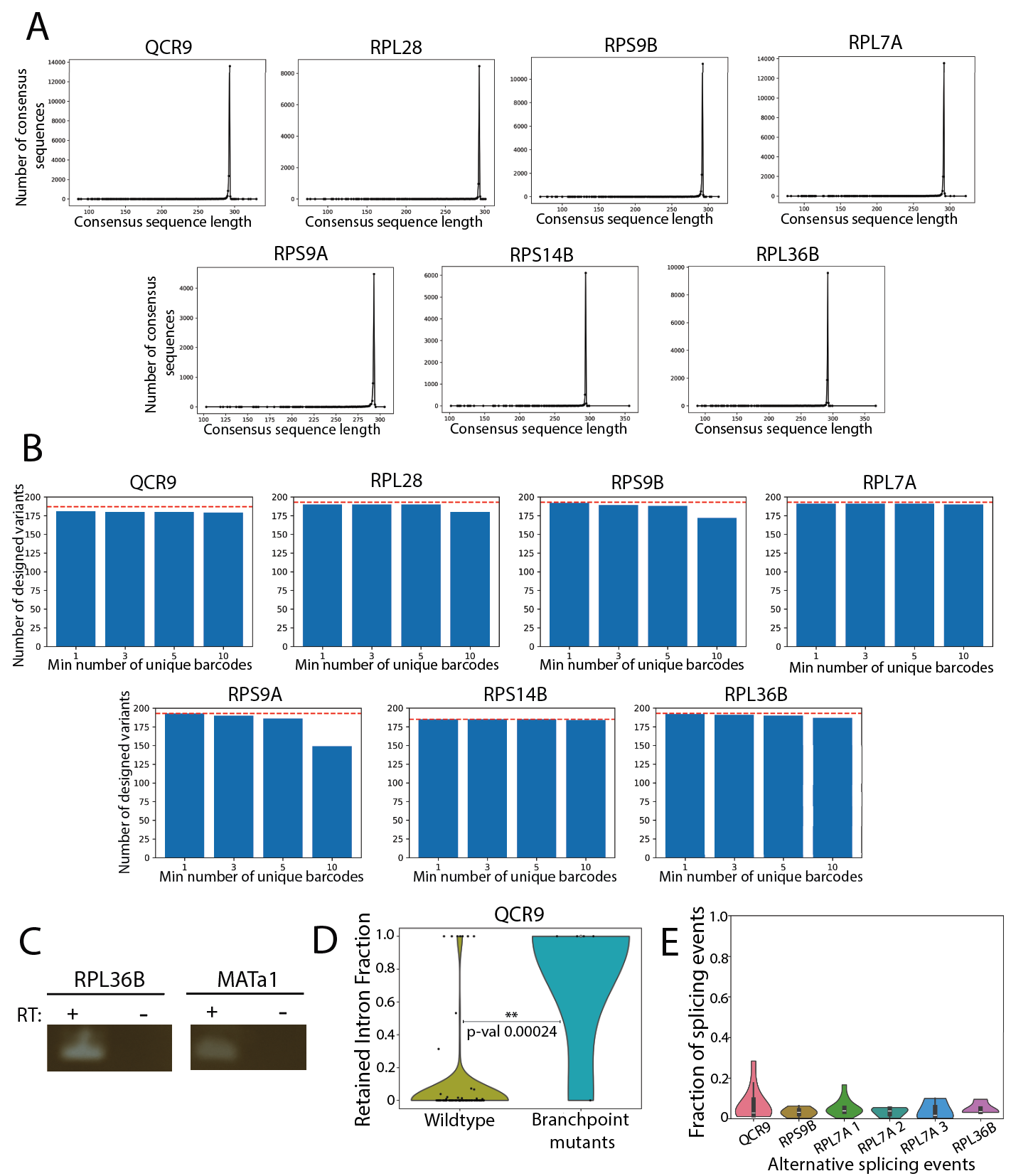

**Figure S20**: Structure variant library gDNA and RNA sequencing validation. A) Histogram of consensus sequence lengths determined for each barcode from gDNA sequencing. B) Number of designed variants assigned to at least 1, 3, 5, or 10 unique barcodes from gDNA sequencing. Red dashed line indicates total number of designed variants per construct. C) RT-PCR for control regions (RNA intervals in RPL36B and MATa1) demonstrating depletion of gDNA from targeted RNA-sequencing library preparation. D) Accumulation of retained introns for QCR9 variants with branch point mutations. P-value from a permutation test for the mean statistic. E) For each of six recurring alternative splicing events across 4 introns, we identified the variant sequences for which these events were observed. These violin plots depict the fraction of splicing events that were alternatively spliced for transcripts from these variant sequences.

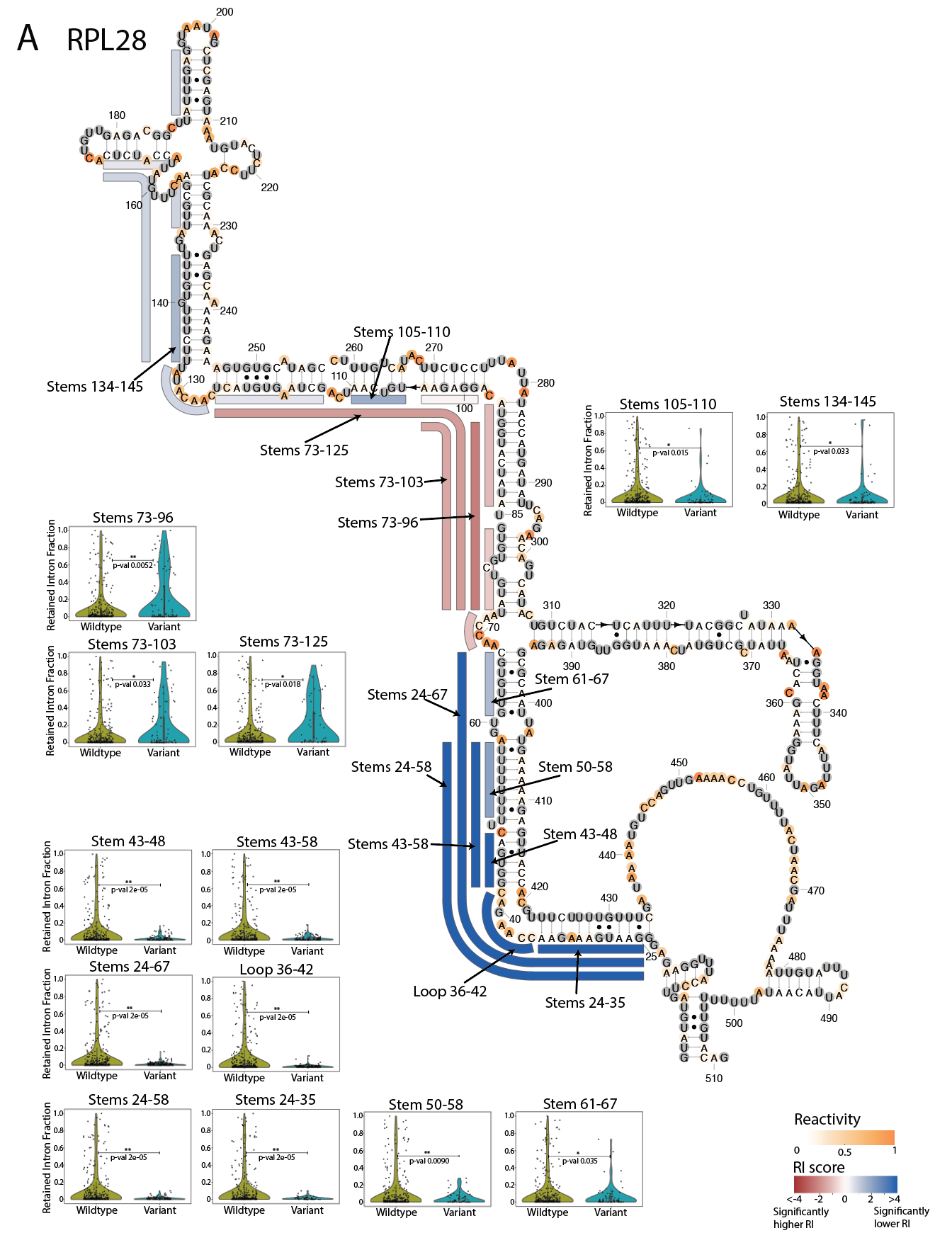

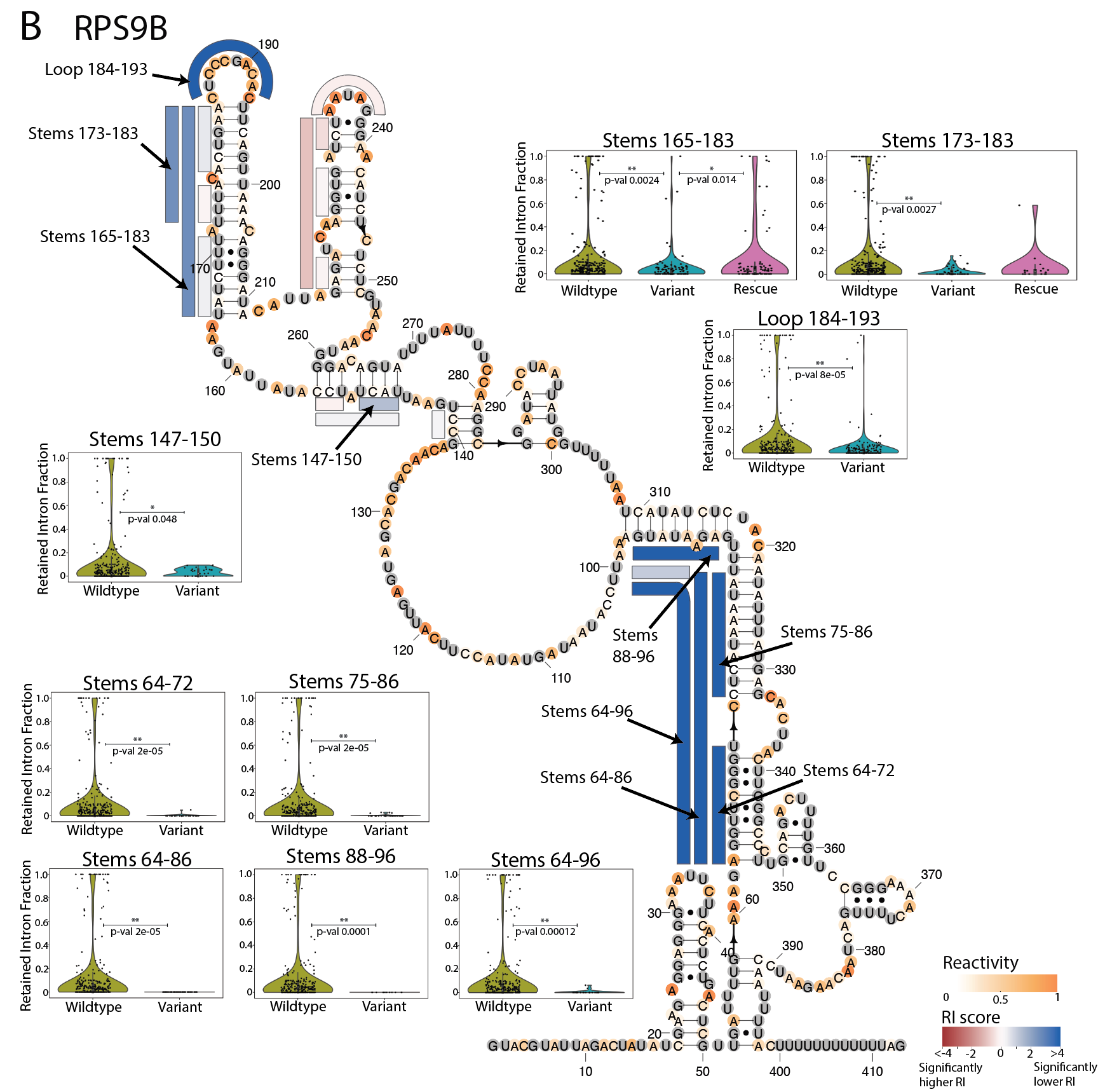

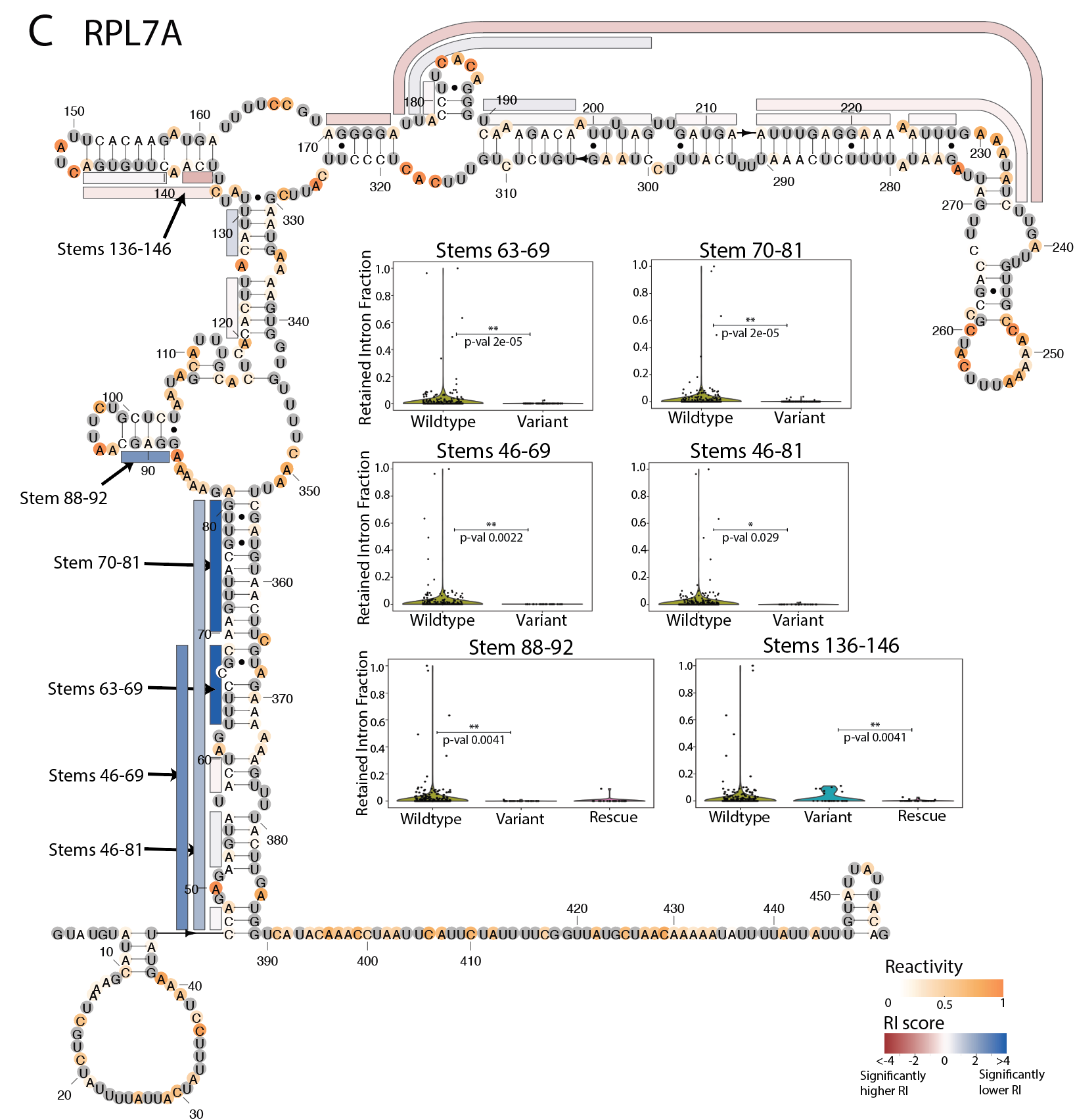

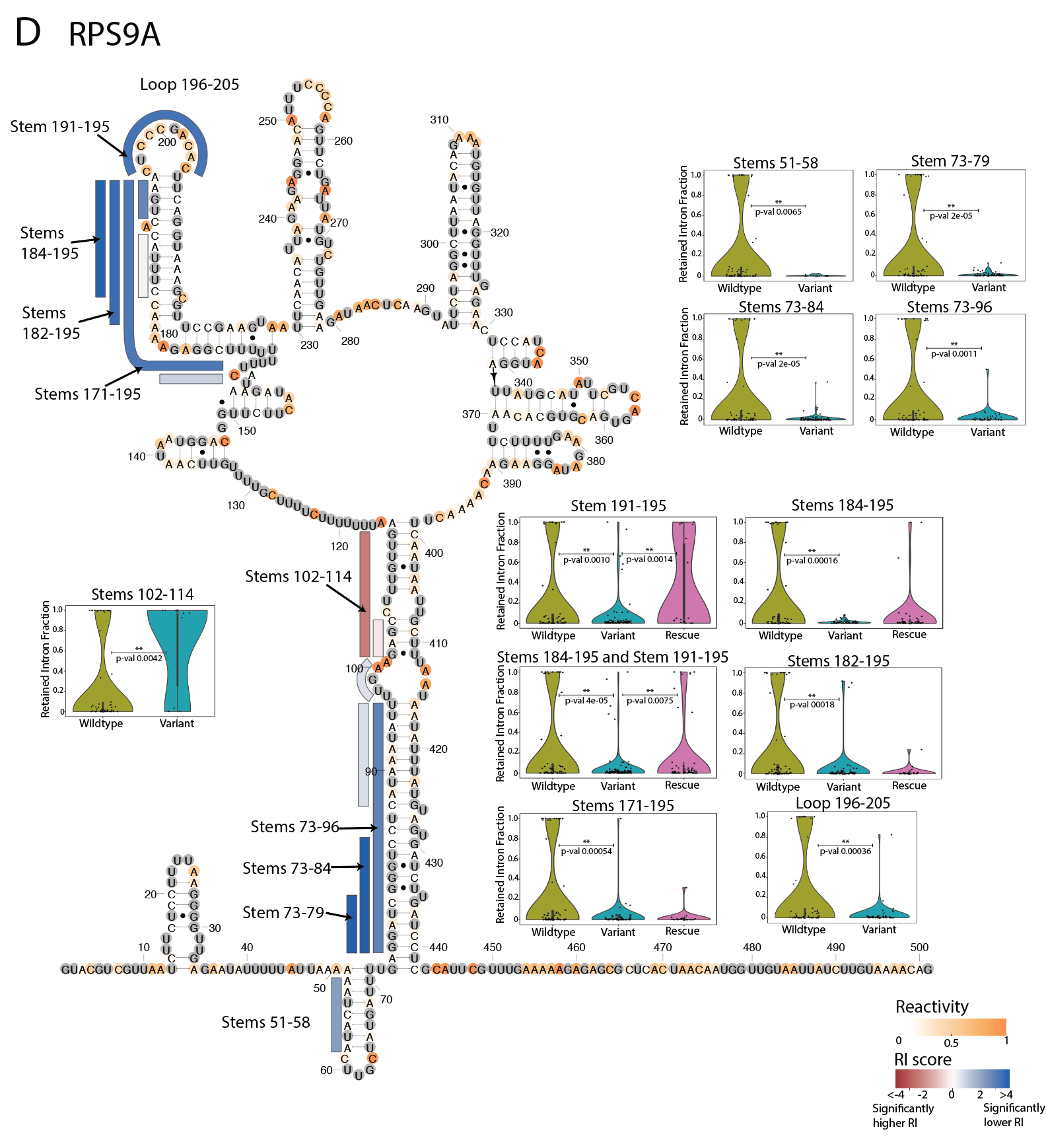

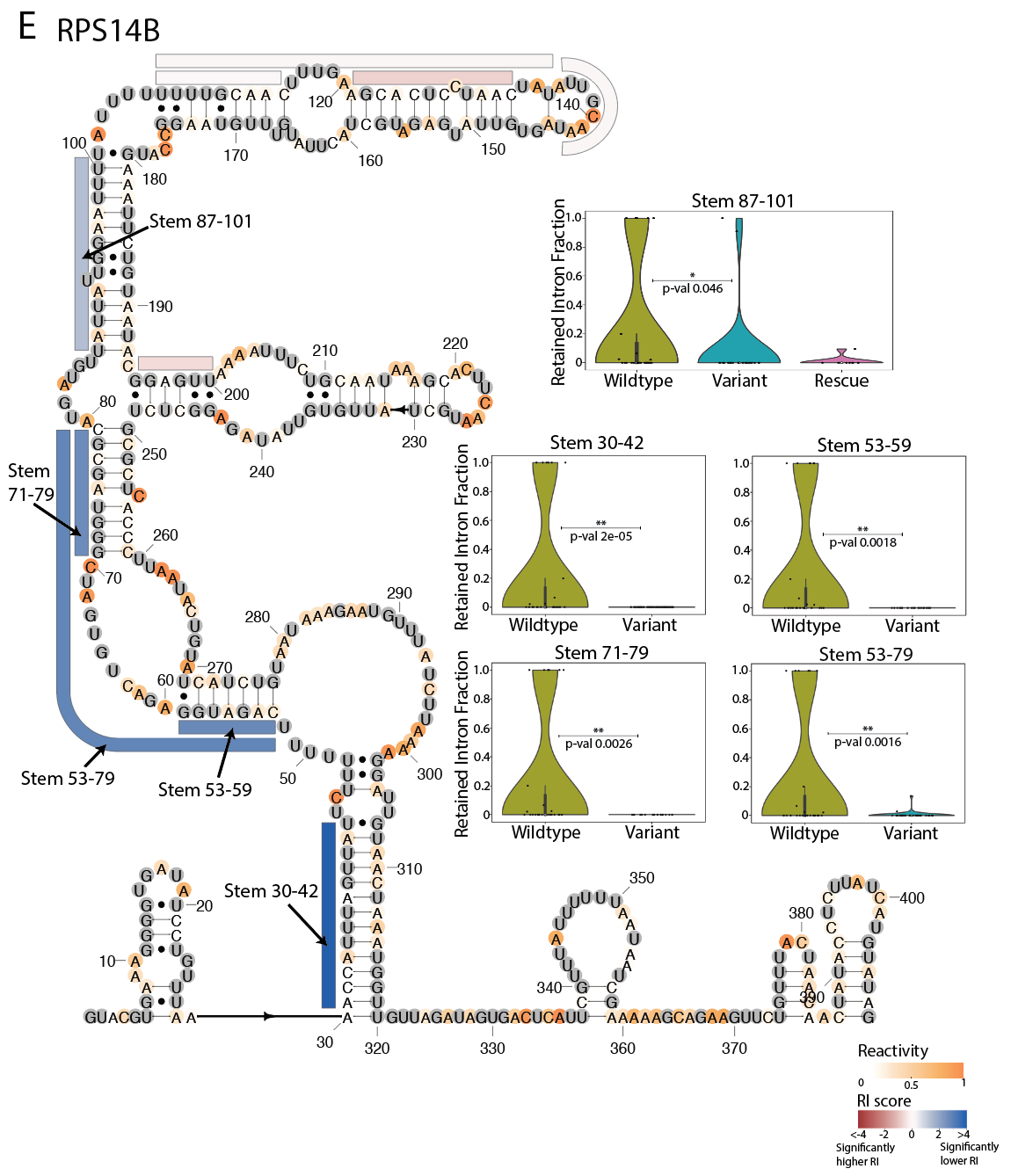

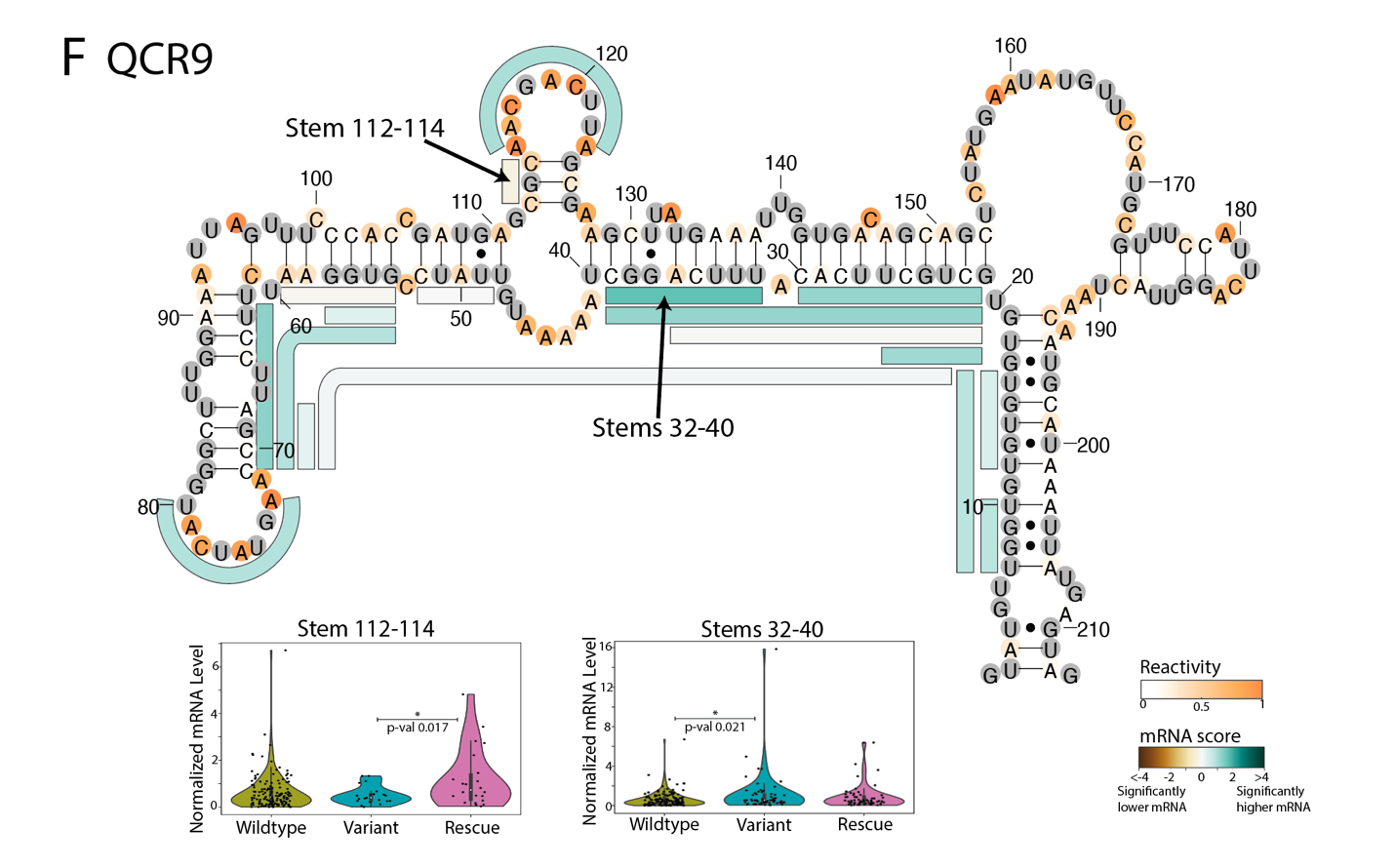

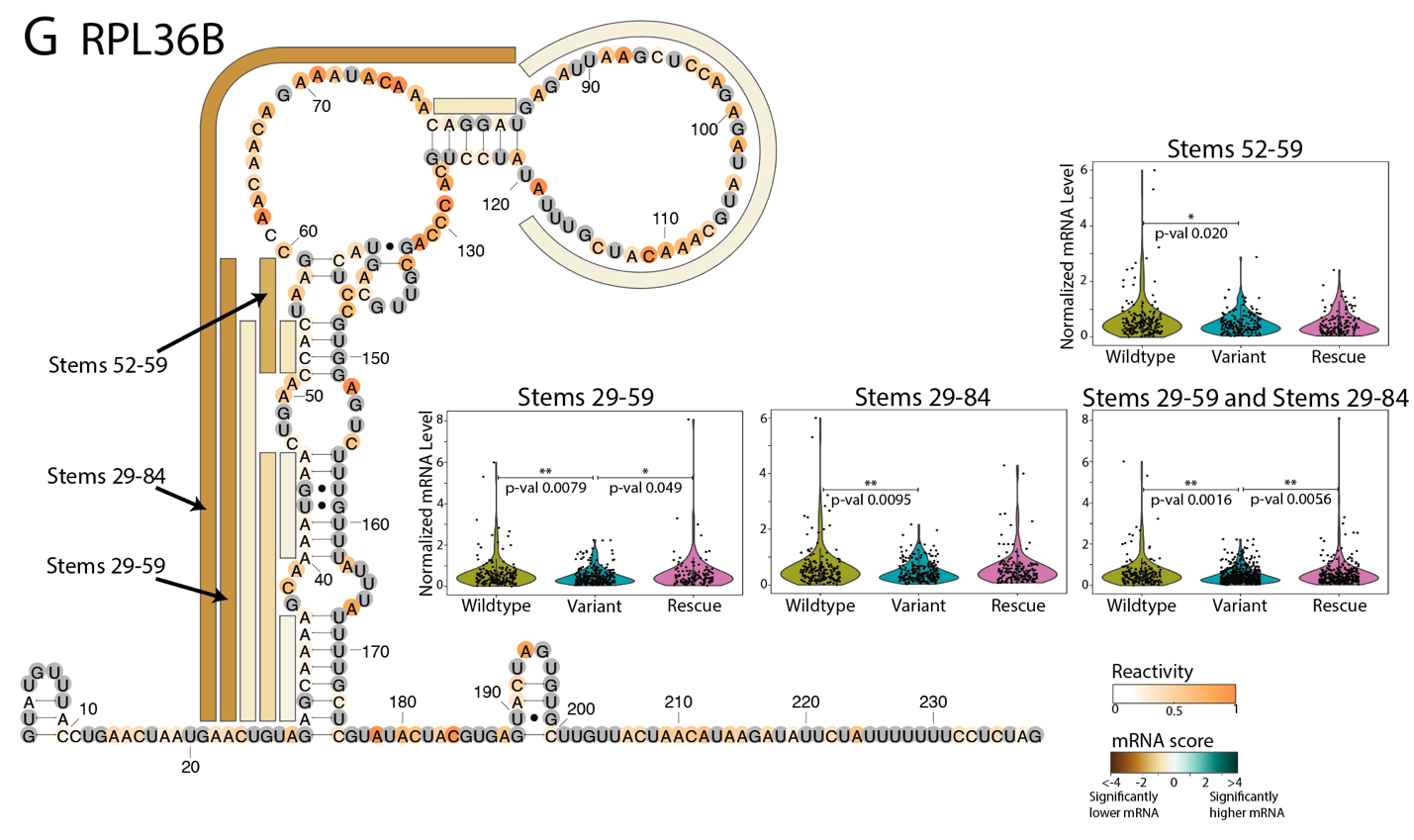

**Figure S21**: Effects of structure variants on retained intron levels for A) RPL28, B) RPS9B, C) RPL7A, D) RPS9A, E) RPS14B, and effects of structure variants on normalized mRNA levels in F) QCR9, and G) RPL36B. For a given stem set or loop, violin plots depict data for the wildtype sequence and all variant sequences, with points for each unique barcode. Data for rescue sequences are shown when included in the intron library. p-values (computed by permutation tests) are indicated for comparisons between wildtype and variant sequences, and between variant and rescue sequences. Secondary structures are colored by reactivity data, and bars alongside the secondary structure indicate stem and loop disruption sets, with each bar representing variant sequences mutating nucleotides across the full extent of the bar. These bars are colored by the retained intron (RI) score (A-E) and mRNA scores (F-G) for the interval. The RI score is the negative log(p-value) comparing RI values between wildtype and variant sequences, and the sign indicates the effect direction, with positive values (shown as blue) for lower variant RI compared to wildtype, and negative values (shown as red) for higher variant RI. Similarly, the mRNA score is computed as the negative log(p-value) comparing normalized mRNA levels between wildtype and variant sequences, with positive (green) values indicating higher variant mRNA levels, and negative (brown) values indicating lower variant mRNA levels.

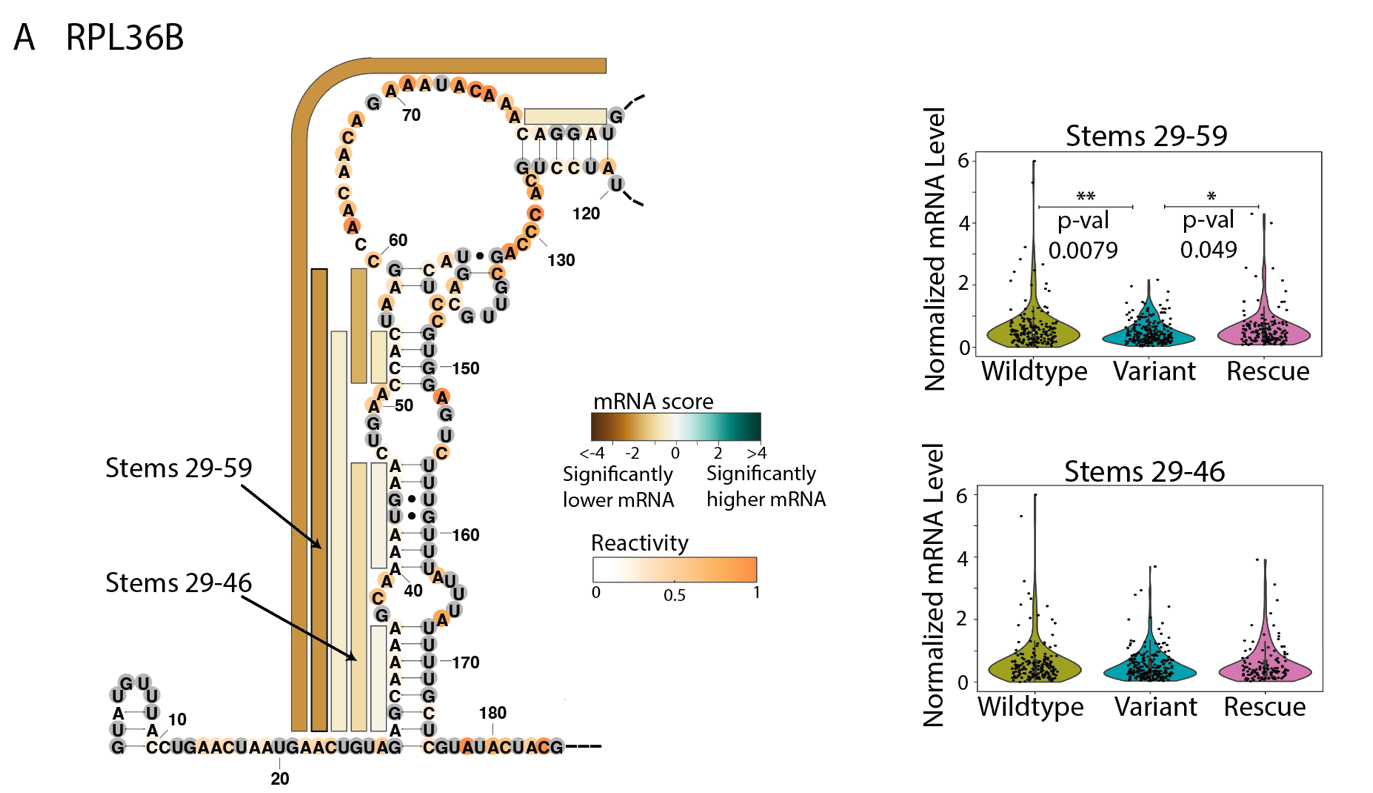

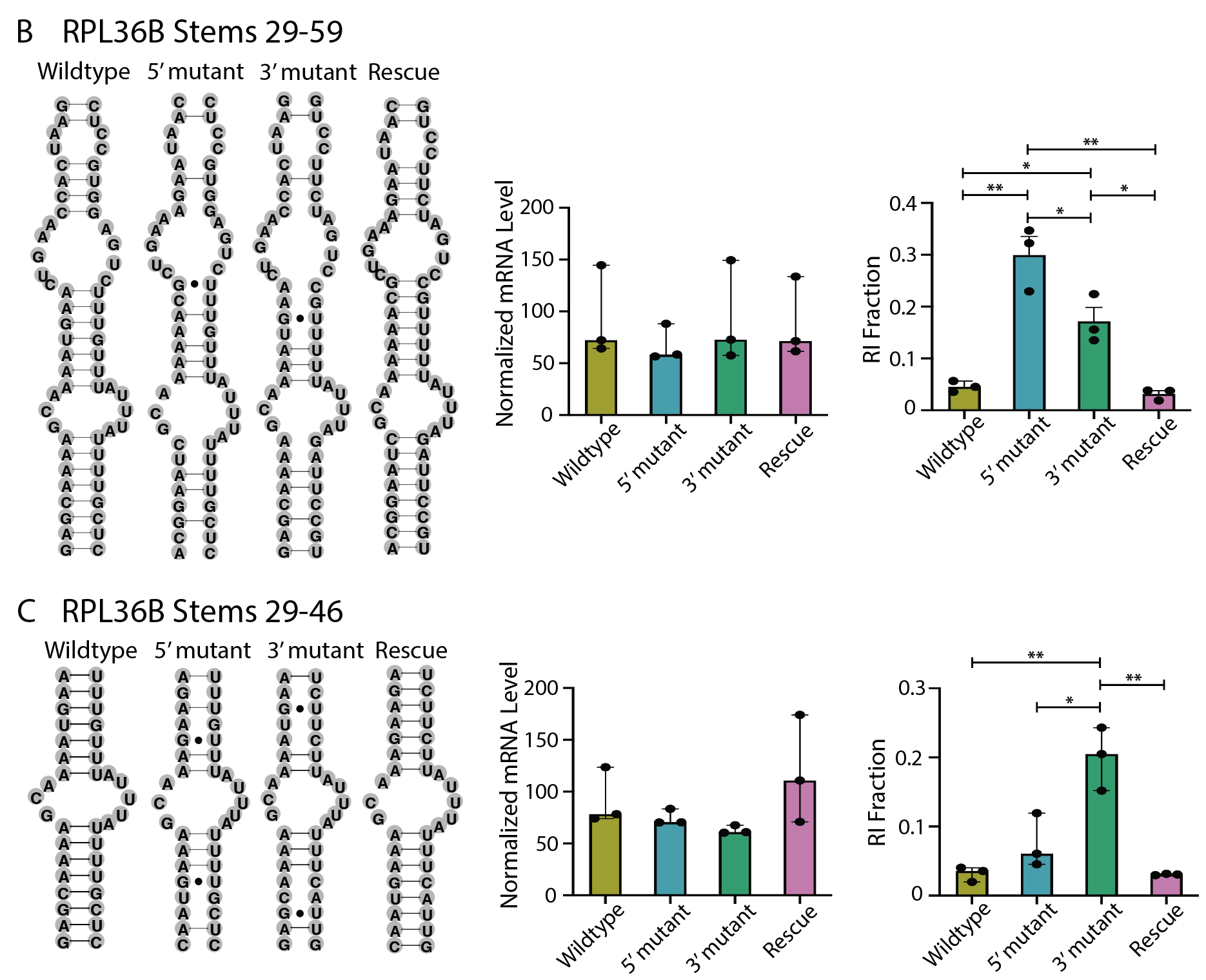

**Figure S22:** Assessing the effects of stem variant and rescue sequences on stem sets in RPL36B with VARS-seq and RT-qPCR. A) The effects of structure variants on normalized mRNA levels for two regions of of the RPL36B intron. For a given stem set or loop, violin plots depict data for the wildtype sequence and all variant sequences, with black points for each unique barcode. p-values are indicated for comparisons between wildtype and variant sequence sets, and between variant and rescue sequence sets. p-values are computed using permutation tests for the difference in mean statistic. Secondary structures are colored by reactivity data. Bars alongside the secondary structure indicate stem and loop disruption sets, with each bar representing a set of variant sequences mutating nucleotides across the full extent of the bar. These bars are colored by the mRNA score for the corresponding stem or loop disruption set. The mRNA score is computed as the negative log(p-value) when comparing normalized mRNA levels between wildtype and variant sequences, with positive (green) values indicating higher variant mRNA levels, and negative (brown) values indicating lower variant mRNA levels. B, C) RI fractions as measured by RT-qPCR for individual strains representing two sets of wildtype, variant, and rescue sequences for RPL36B stem sets. Stem variants are shown on the left, and RT-qPCR data are shown for 3 biological replicates on the right. p-values are computed with 2-way ANOVA tests with multiple comparisons.

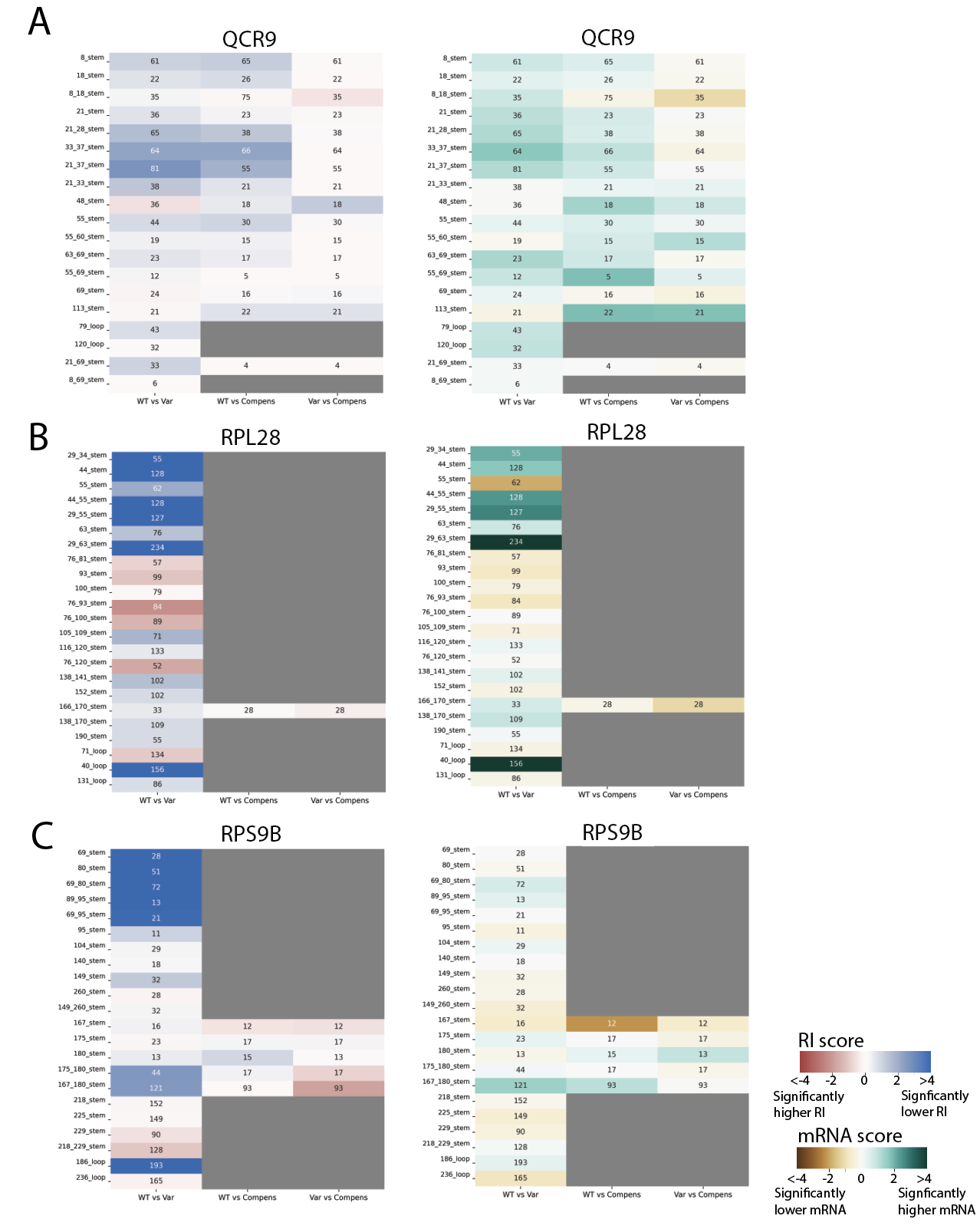

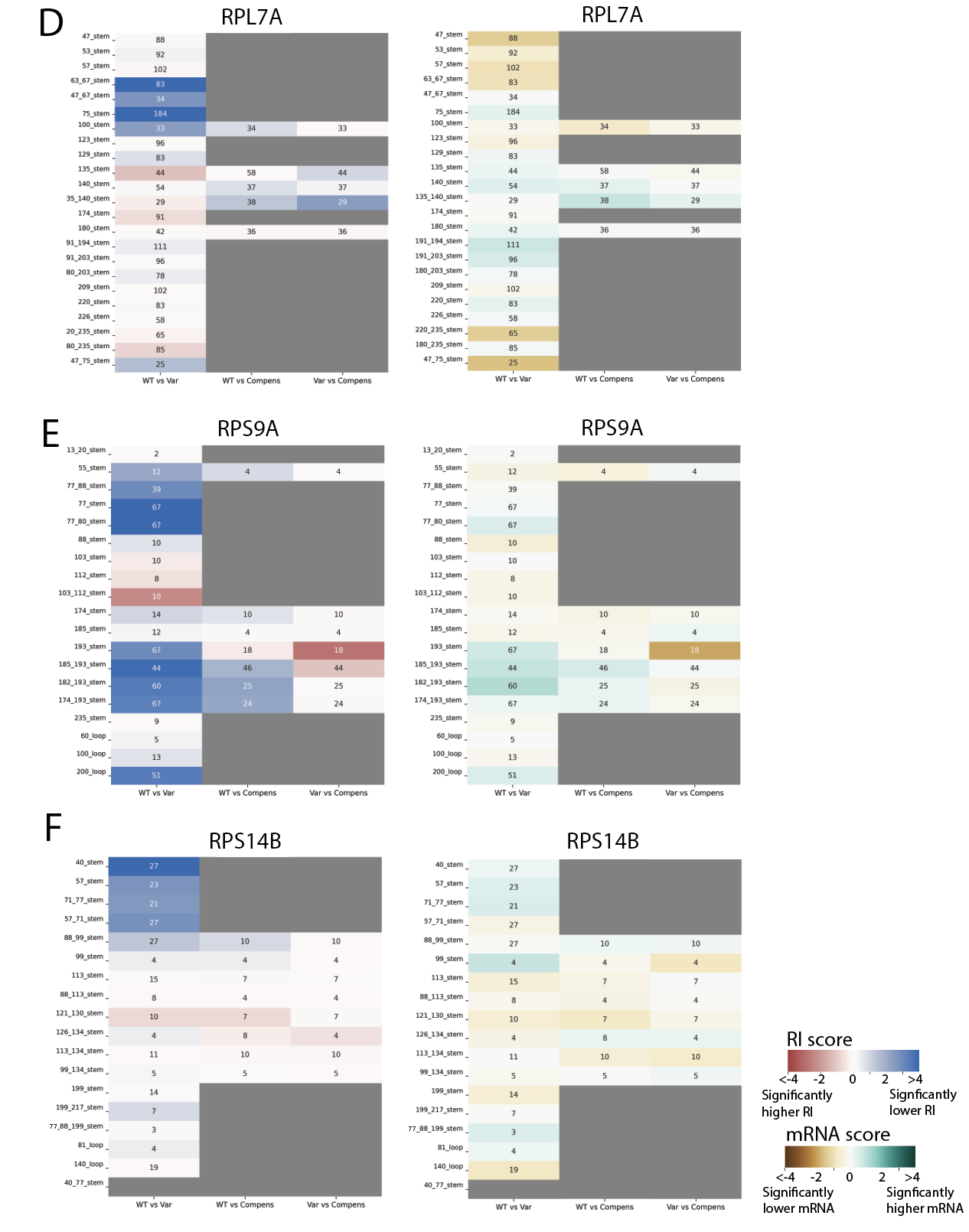

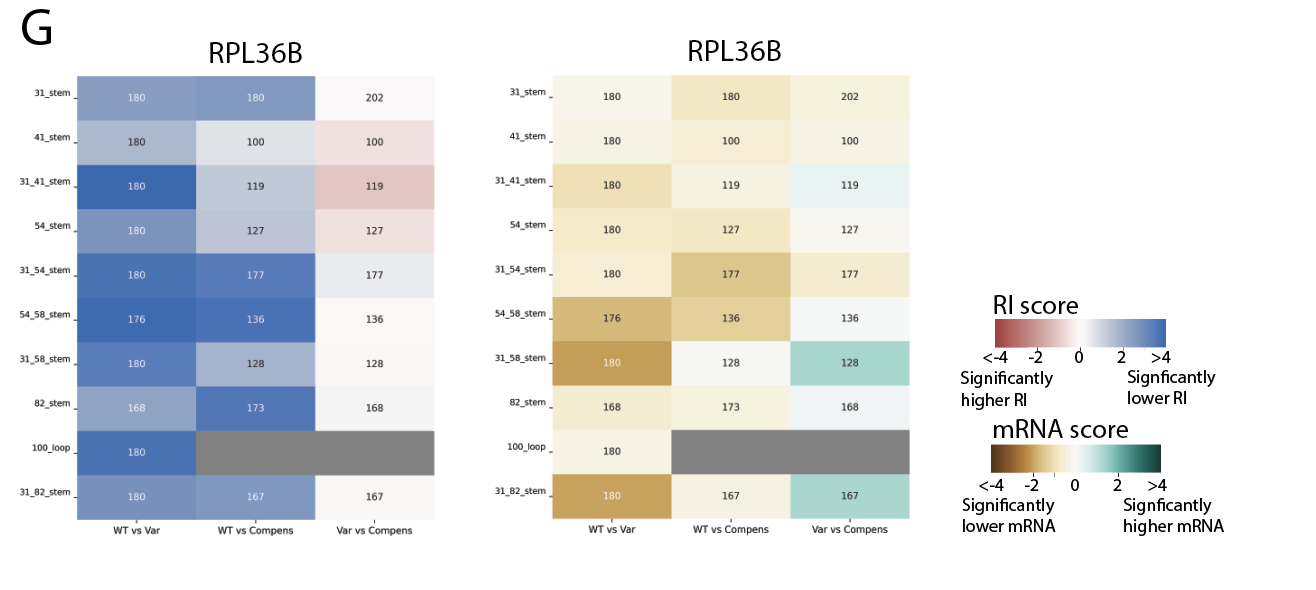

**Figure S23**: Heatmaps depicting the effects of structure variants on retained intron fractions and normalized mRNA levels for A) QCR9, B) RPL28, C) RPS9B, D) RPL7A, E) RPS9A, F) RPS14B, and G) RPL36B. Each row corresponds to a stem or loop disruption set, and the nucleotides these labels correspond to are noted in Table S3. Boxes are colored based on the significance of the difference in RI fraction or mRNA score between sequence sets, and numbers in boxes indicate the minimum number of sequences used for each comparison. Boxes are gray if comparisons involve 2 or fewer sequences, either due to low coverage or because the sequence set was not included in the library. The heatmap compares the following sets of sequences: 1) left column: wildtype sequences are the base set and variant sequences are the comparison set; 2) middle column: wildtype sequences are again the base set and compensatory rescue sequences are the comparison set; 3) right column: variant sequences are the base set and compensatory rescue sequences are the comparison set. The heatmaps are colored by the retained intron (RI) scores (left) and mRNA scores (right). The RI score is the negative log(p-value) comparing RI values between sets, and the sign indicates the effect direction, with positive values (shown as blue) for lower RI values in the comparison set compared to the base set, and negative values (shown as red) for higher RI values in the comparison set. Similarly, the mRNA score is computed as the negative log(p-value) comparing normalized mRNA levels between sets, with positive (green) values indicating higher comparison set mRNA levels, and negative (brown) values indicating lower comparison set mRNA levels. P-values are computed by permutation tests for the difference in mean statistic.

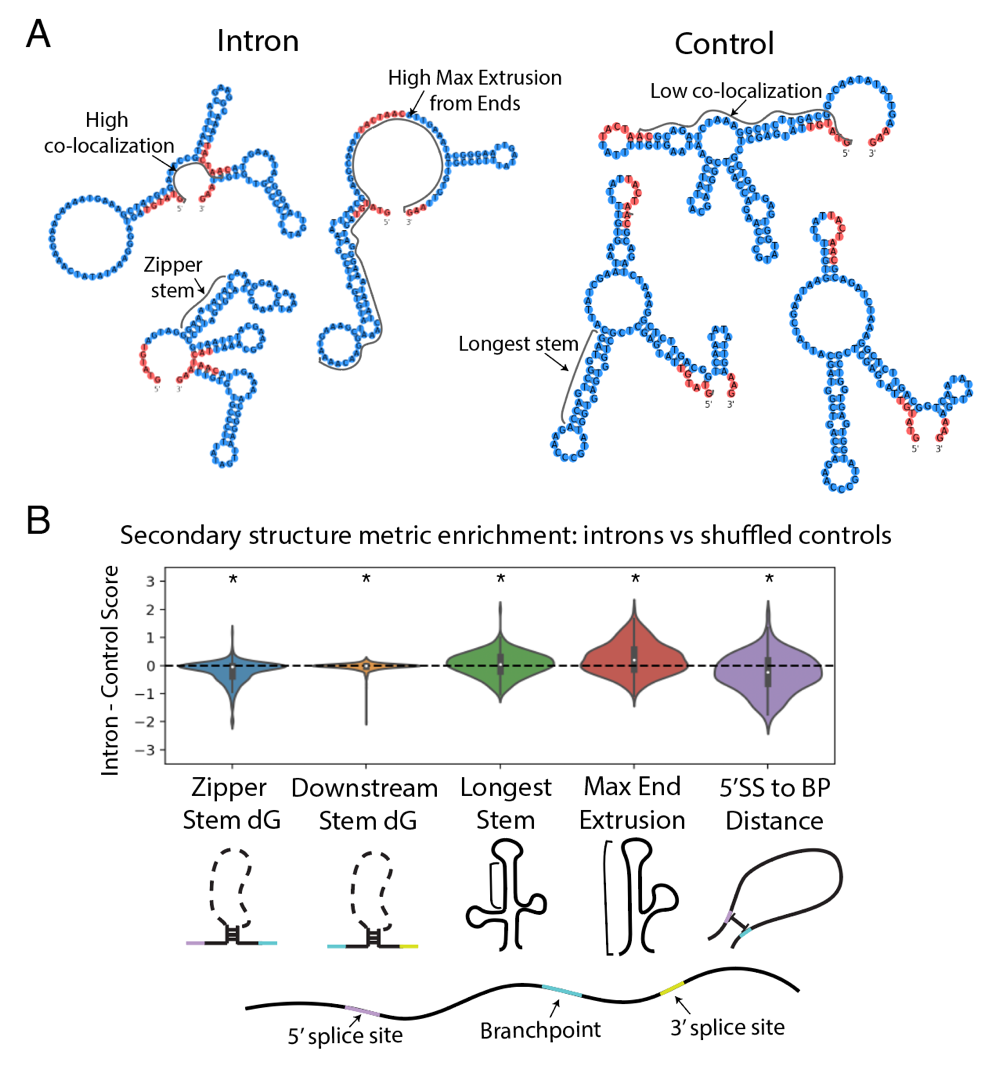

**Figure S24:** Schematics for *de novo* secondary structure feature prediction. A) Sample structures from an intron secondary structure ensemble and control secondary structure ensemble, with structural features annotated. B) Enrichment of secondary structure features comparing intron sequences to shuffled sequences using secondary structure ensembles predicted from Vienna 2.0. *p-value < 0.01 by Wilcoxon ranked-sum test.

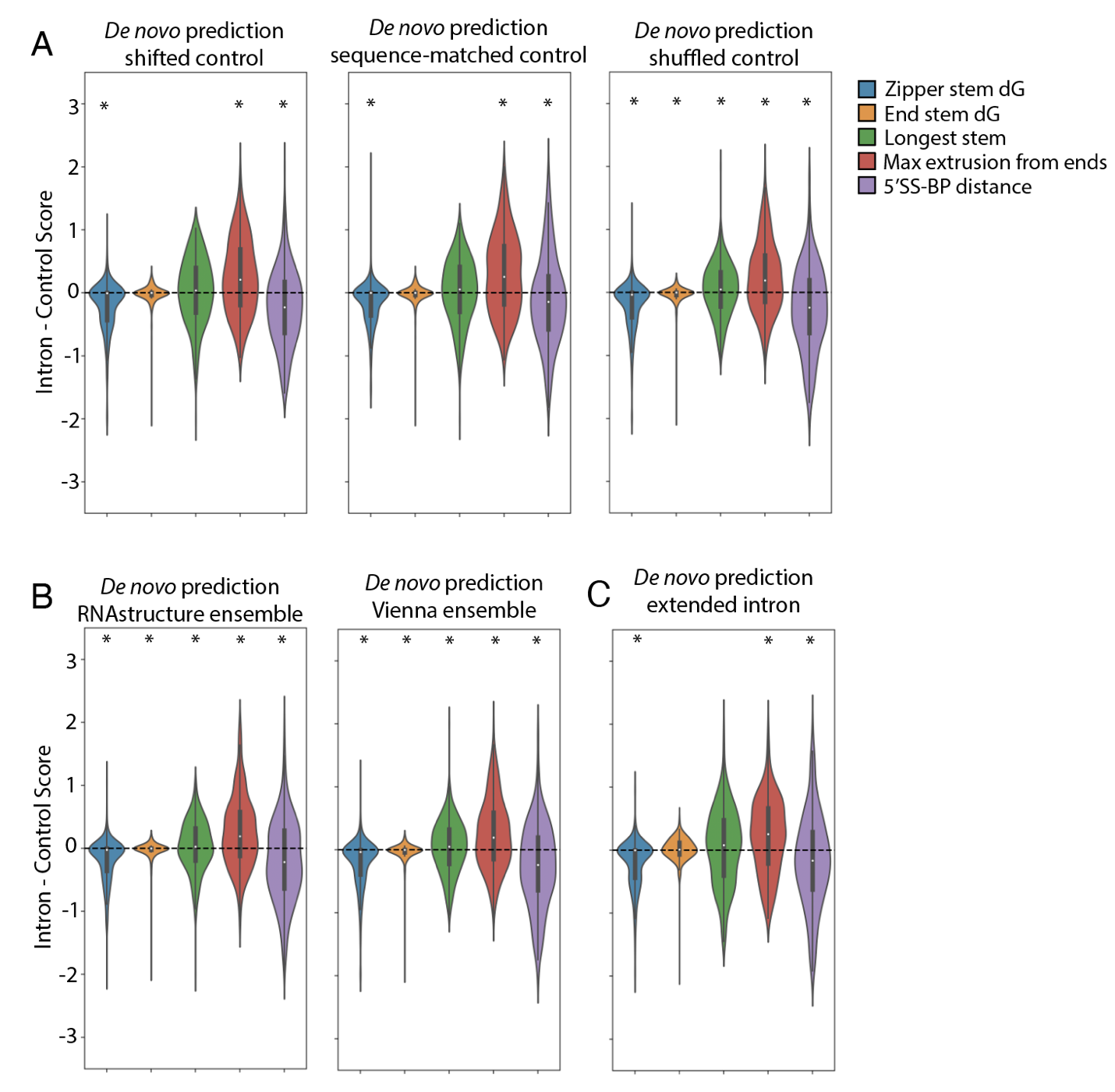

**Figure S25:** Computational prediction of secondary structure feature enrichment with various parameters. A) Feature enrichment when comparing intron sequences to a various control sequence sets. B) Feature enrichment when predicting secondary structure ensembles with RNAstructure vs Vienna, comparing introns to shuffled controls. C) Feature enrichment when including a 50 nucleotide sequence context upstream and downstream of the intron. *p-value < 0.01 from Wilcoxon ranked-sum test.

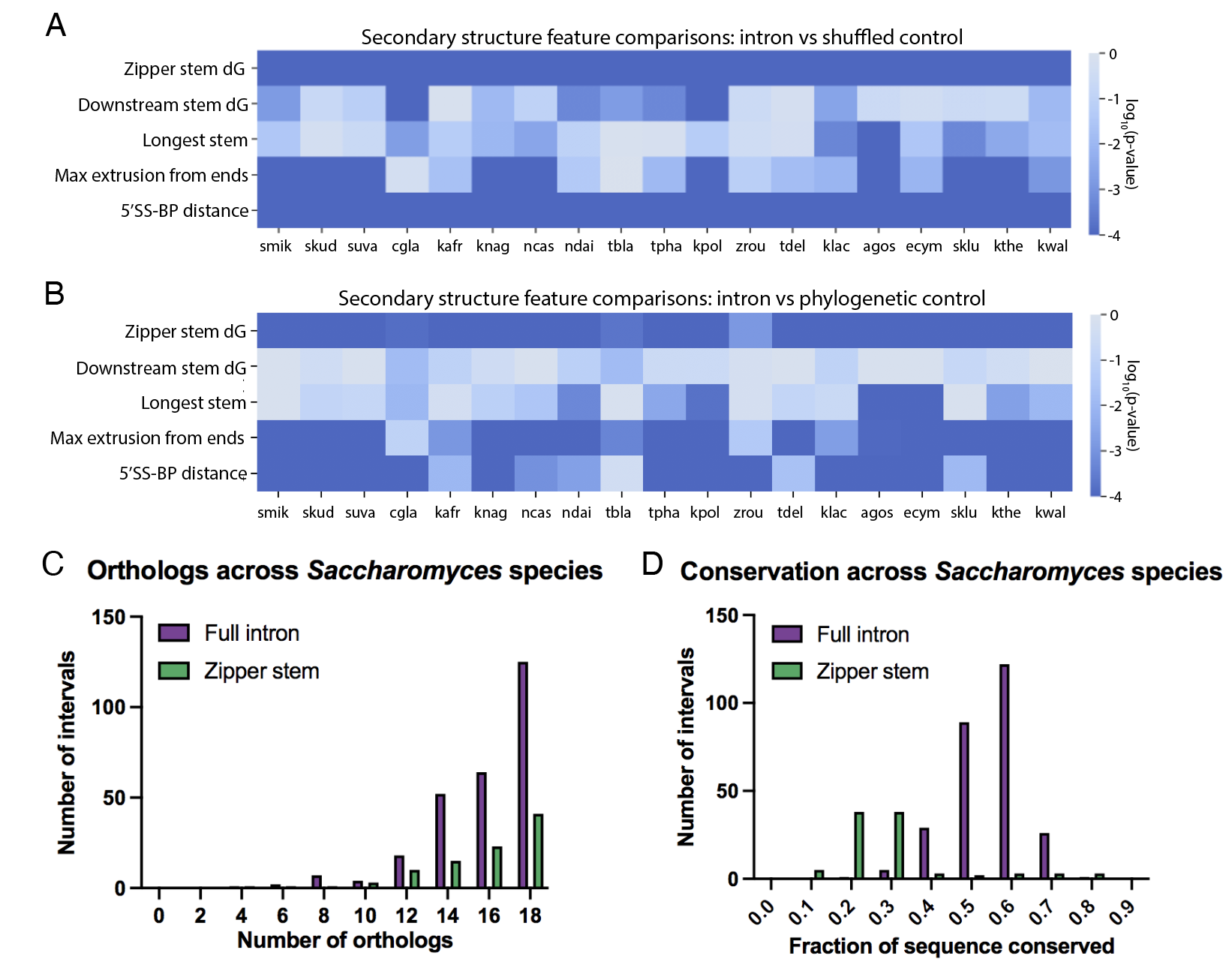

**Figure S26**: Intron secondary structure properties across *Saccharomyces* genus. A) P-values (log scale, base 10) as computed by Wilcoxon ranked sum test comparing secondary structure metrics for intron and shuffled sequence control ensembles across *Saccharomyces* yeast species. B) P-values computed by Wilcoxon ranked sum test for secondary structure metric comparisons between introns and phylogenetic control sequences (log scale, base 10). C) Statistics on the number of orthologs for each *S. cerevisiae* intron and zipper stem across the *Saccharomyces* genus. D) Statistics on sequence conservation for introns (full sequence and zipper stem sequences specifically) across the genus. The species analyzed in this figure are as follows, with label abbreviations noted: *E. gossypii* (agos), *C. glabrata* (cgla), *E. cymbalariae* (ecym), *K. africana* (kafr), *K. lactis* (klac), *K. naganishii* (knag), *V. polyspora* (kpol), *L. thermotolerans* (kthe), *L. waltii* (kwal), *N. castellii* (ncas), *N. dairenensis* (ndai), *S. kudiavzevii* (skud), *S. mikatae* (smik), *S. uvarum* (suva), *T. blattae* (tbla), *T. delbrueckii* (tdel), *T. phaffii* (tpha), *Z. rouxii* (zrou).

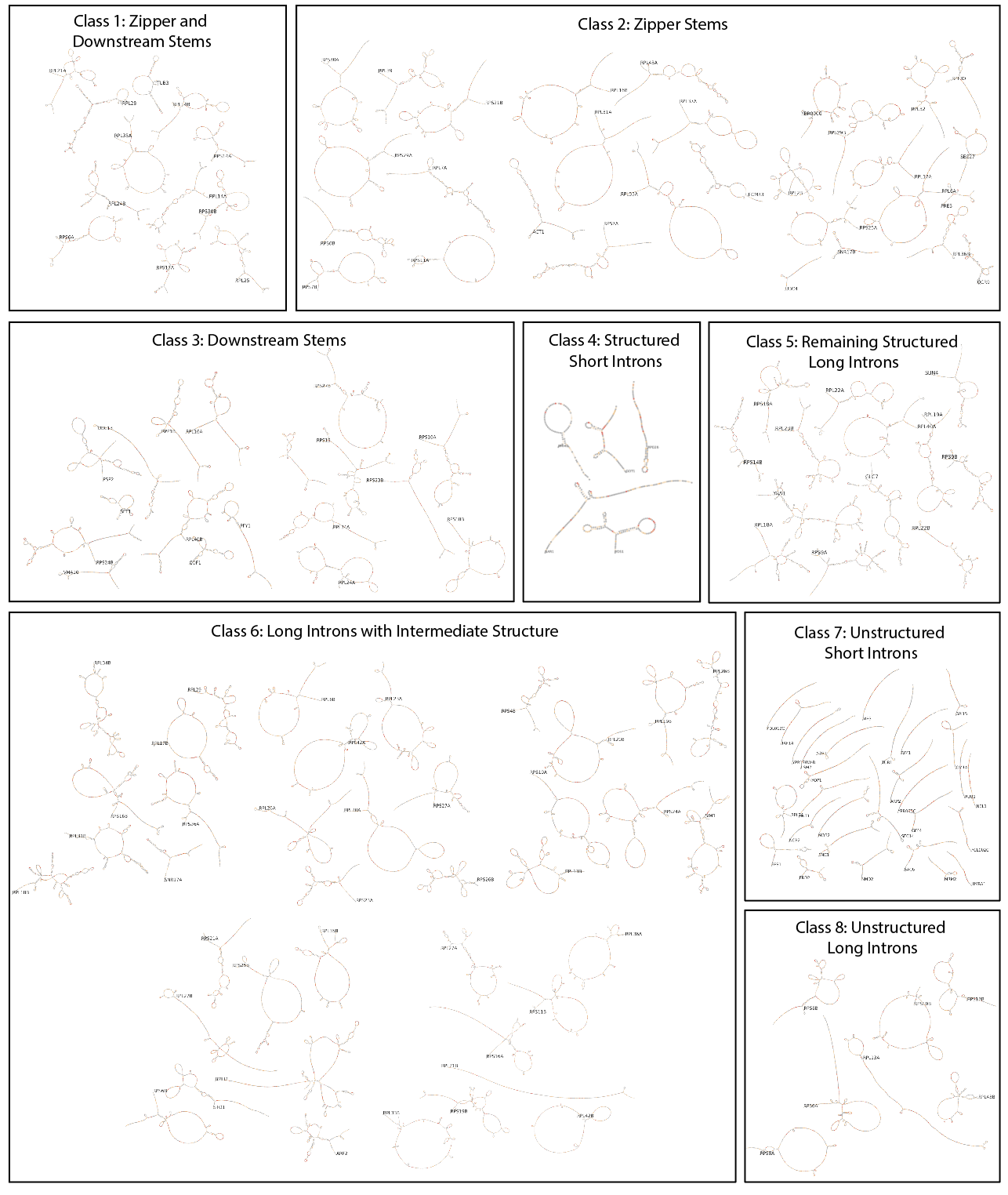

**Figure S27:** One-page overview of secondary structures and overlaid DMS reactivity profiles for all introns with sufficient coverage from DMS-MaPseq. Introns are grouped into classes from hierarchical clustering (Fig. 5). Secondary structures are depicted schematically using RiboGraphViz (<https://github.com/DasLab/RiboGraphViz>). Gene names are included for each intron.

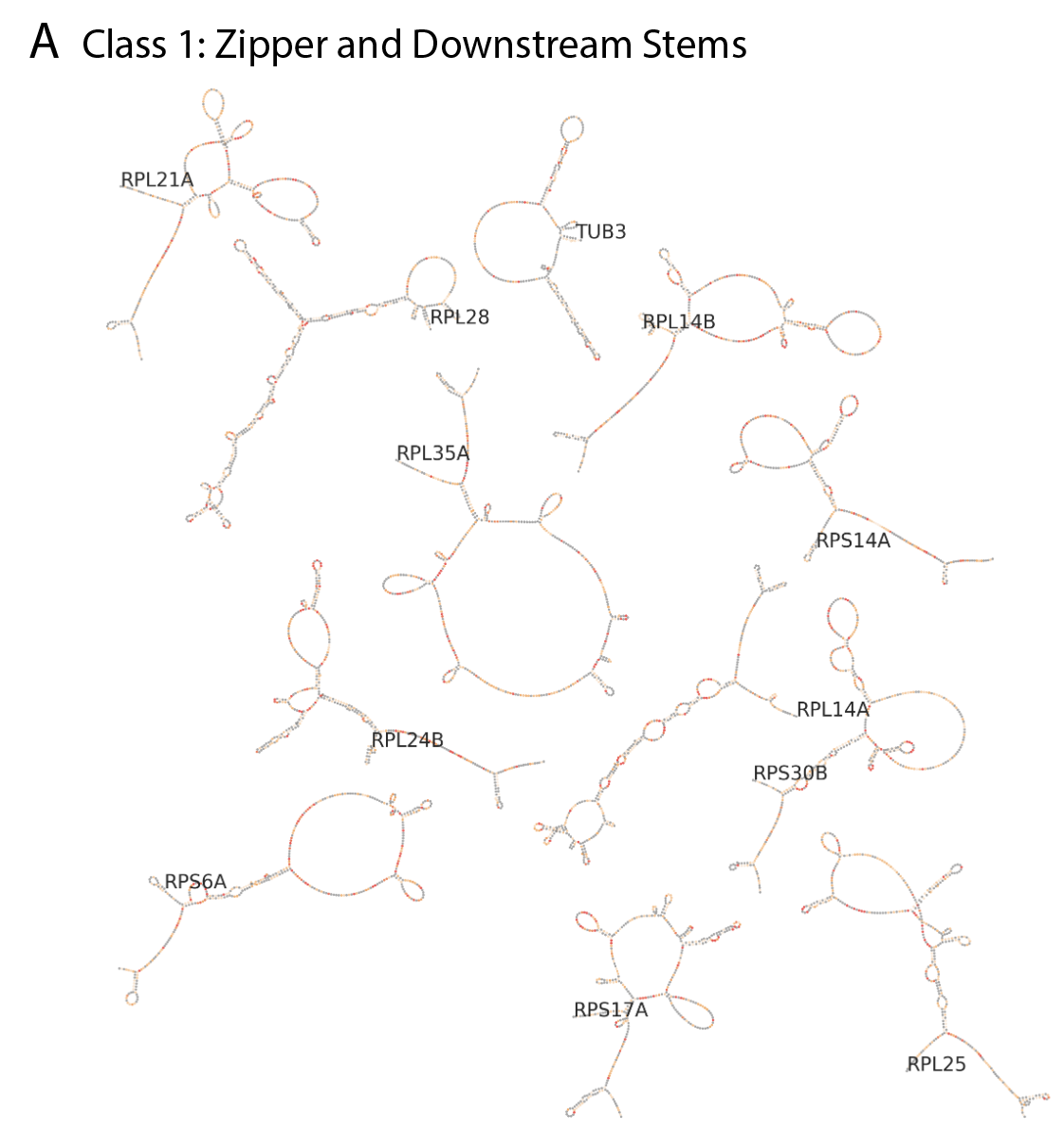

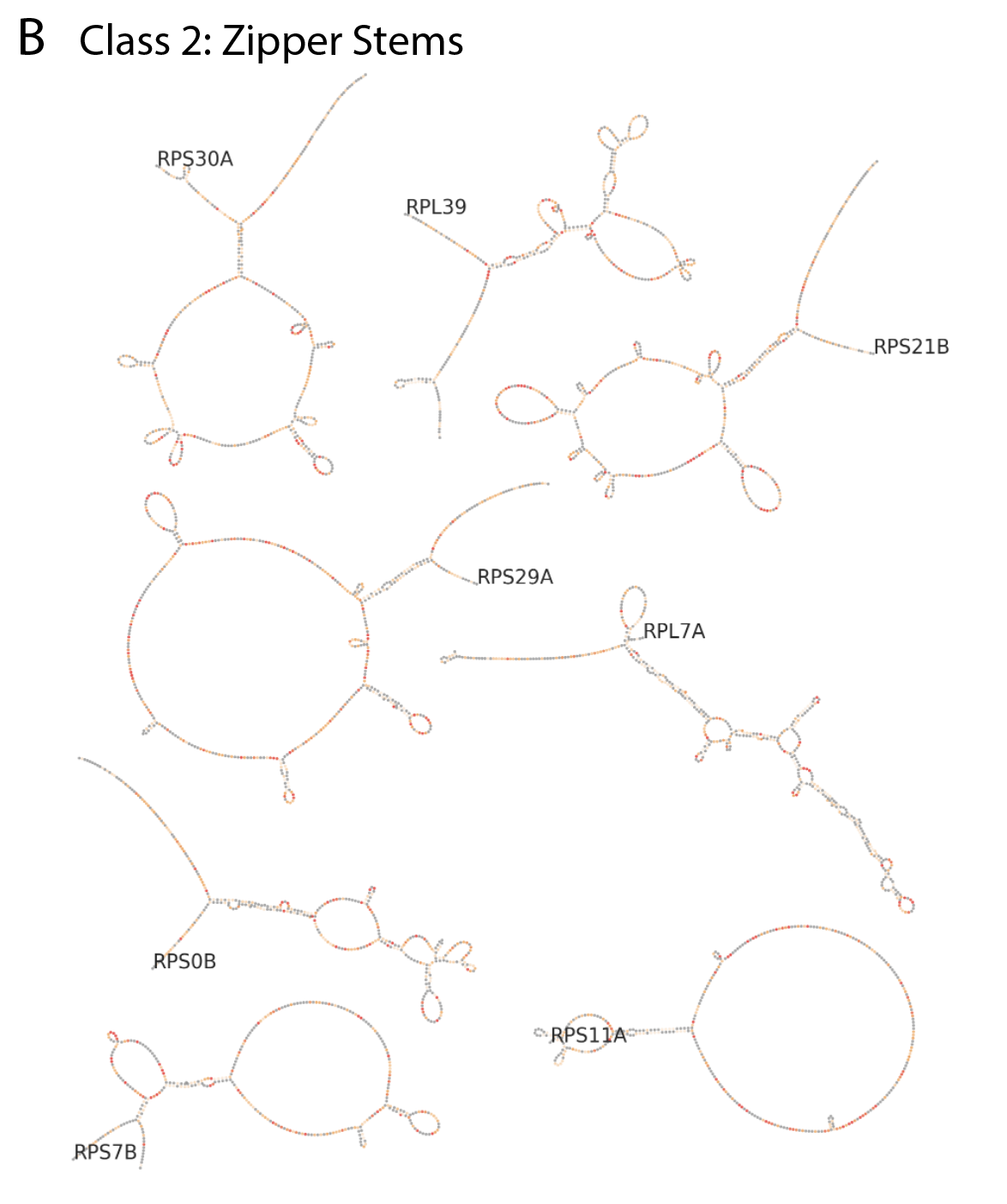

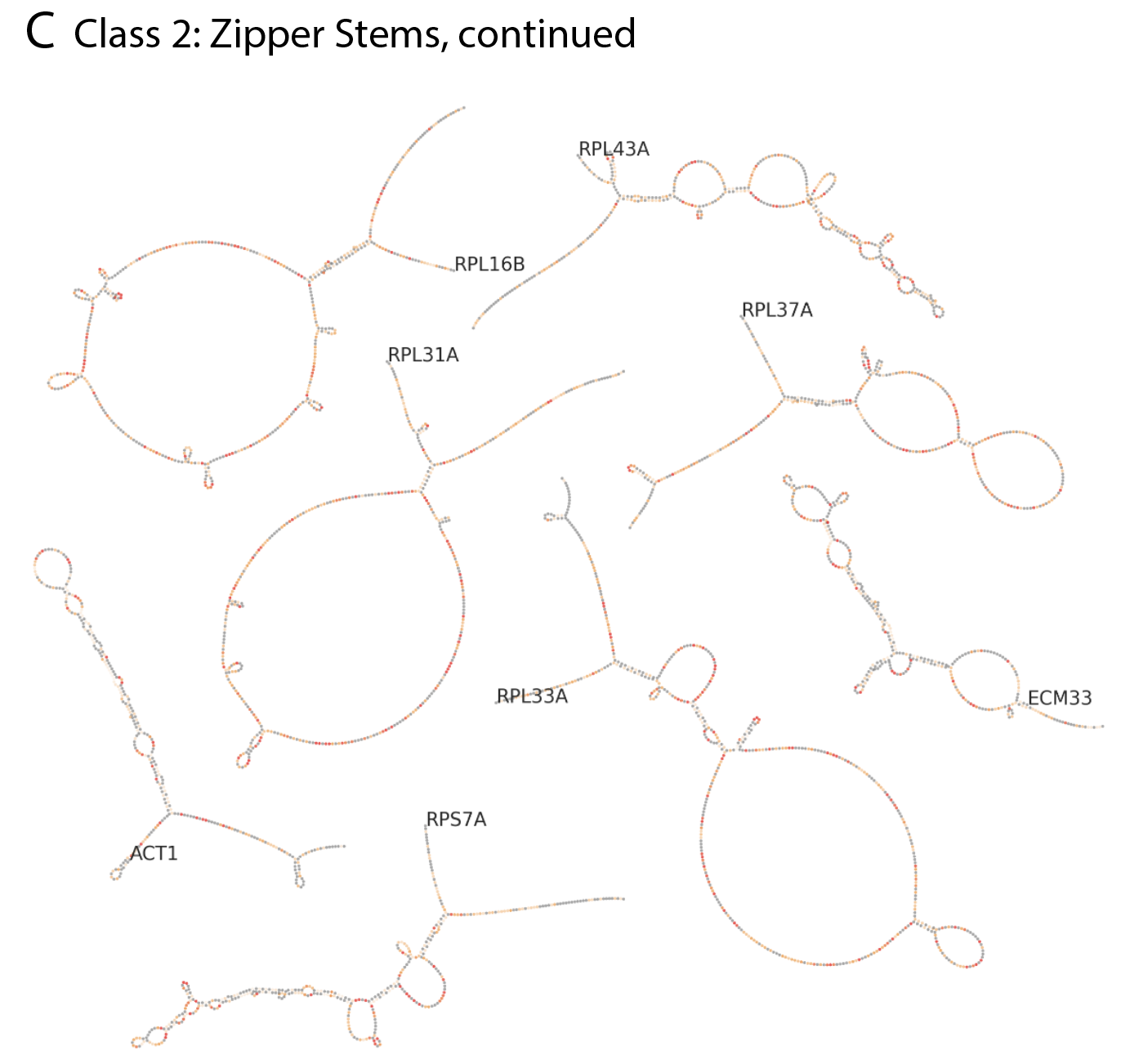

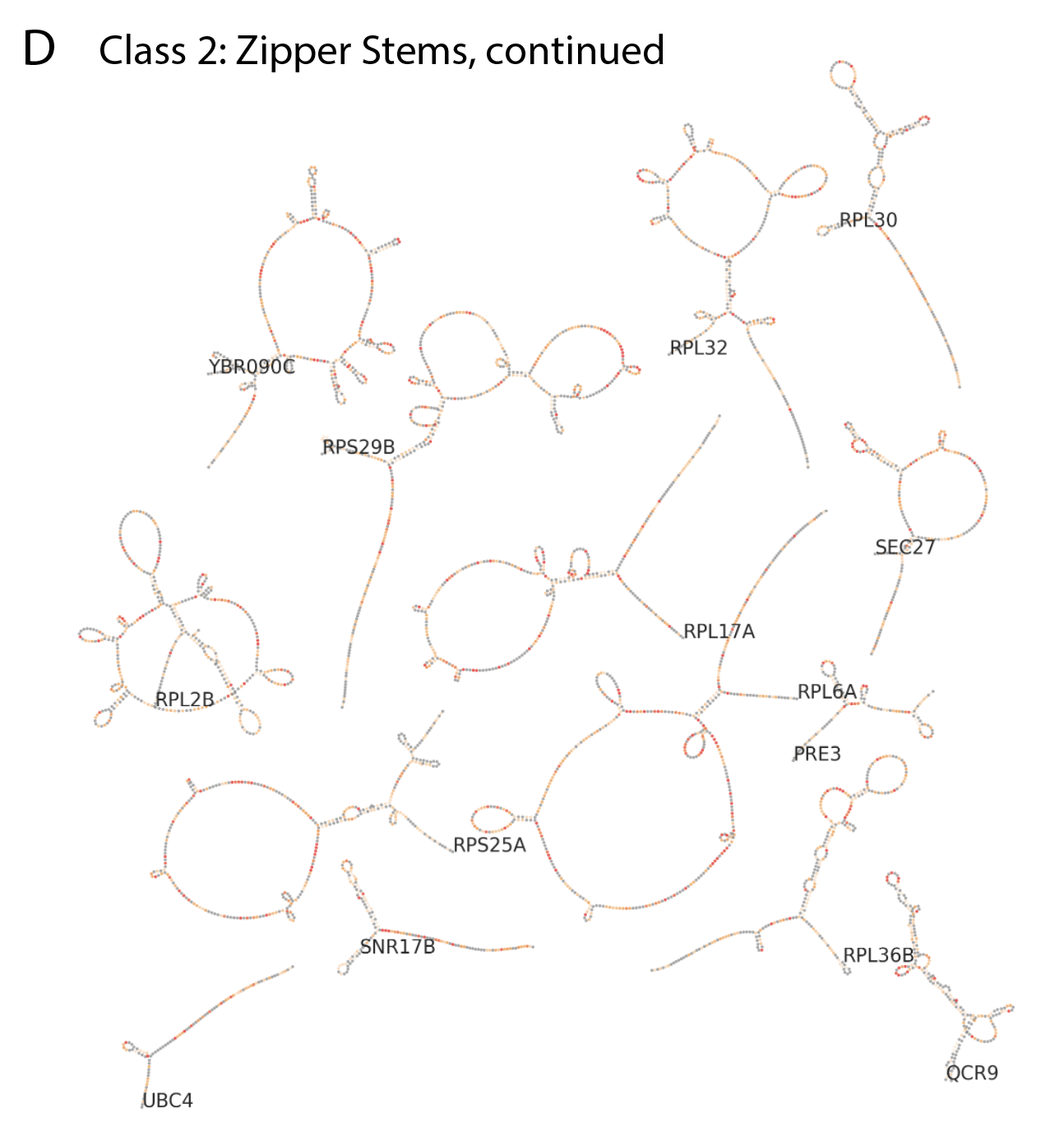

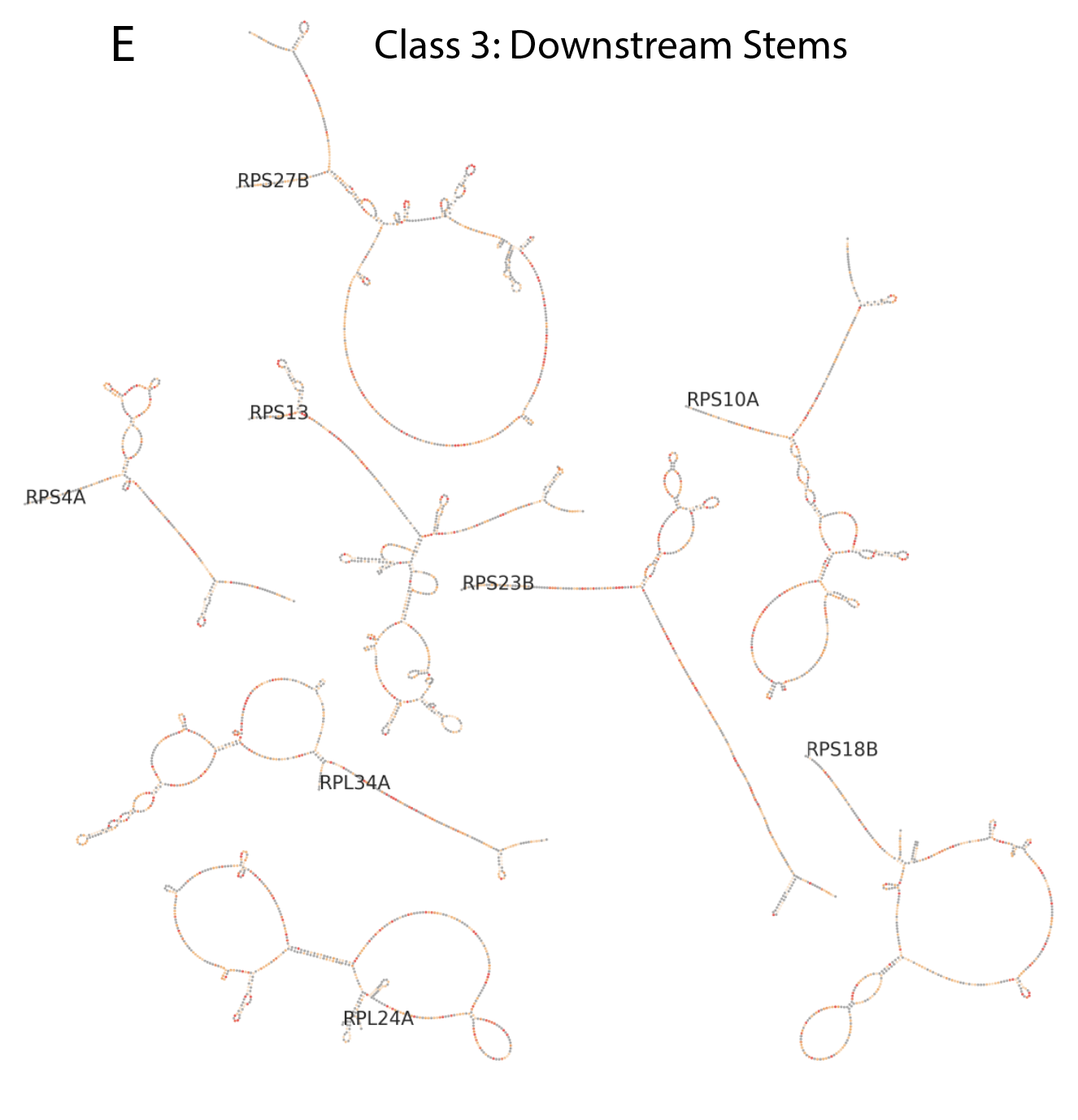

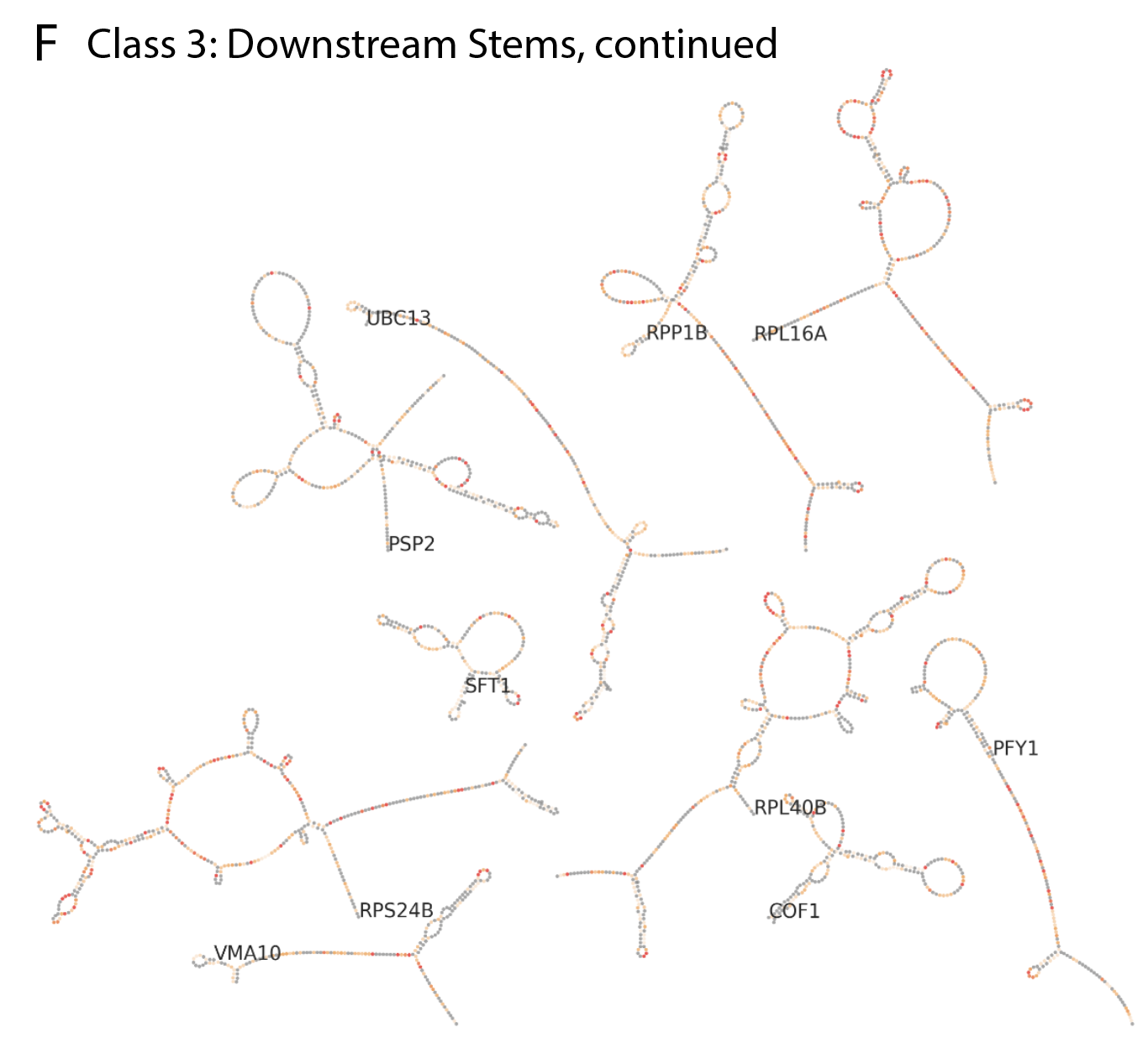

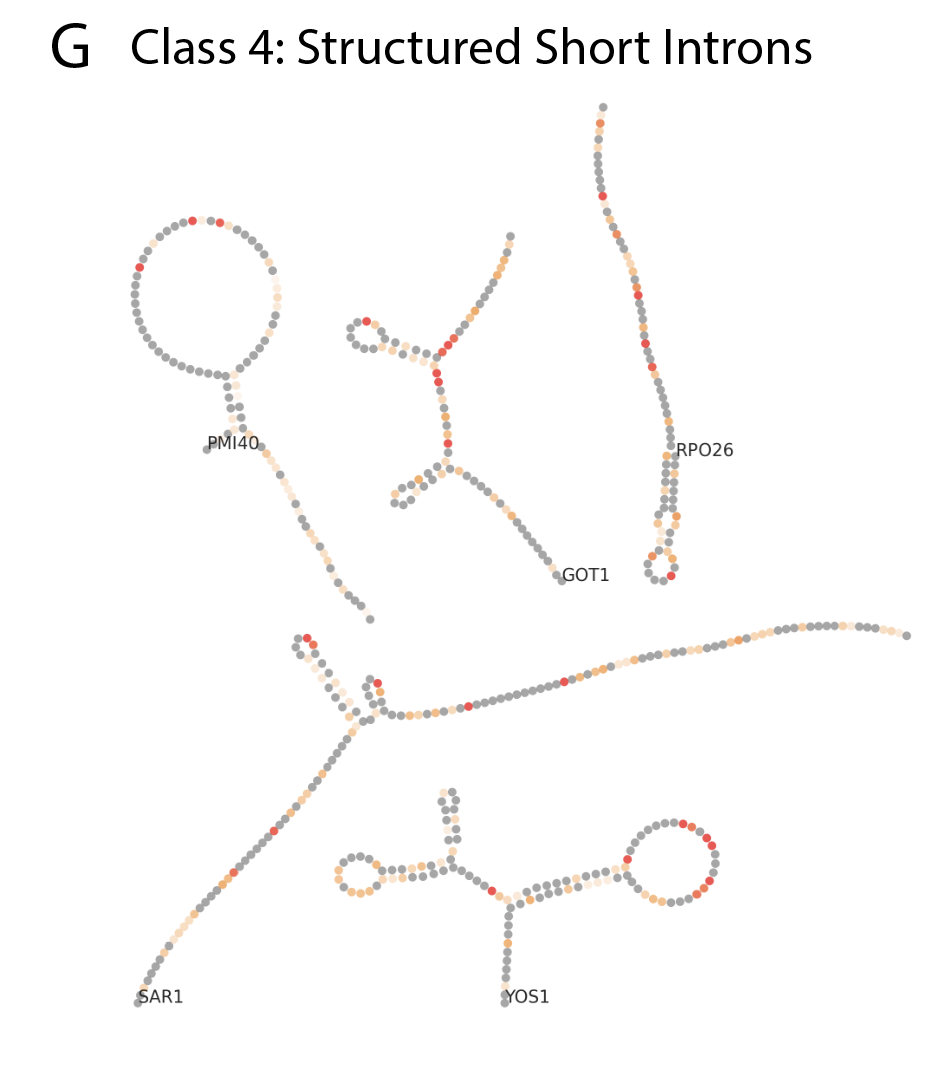

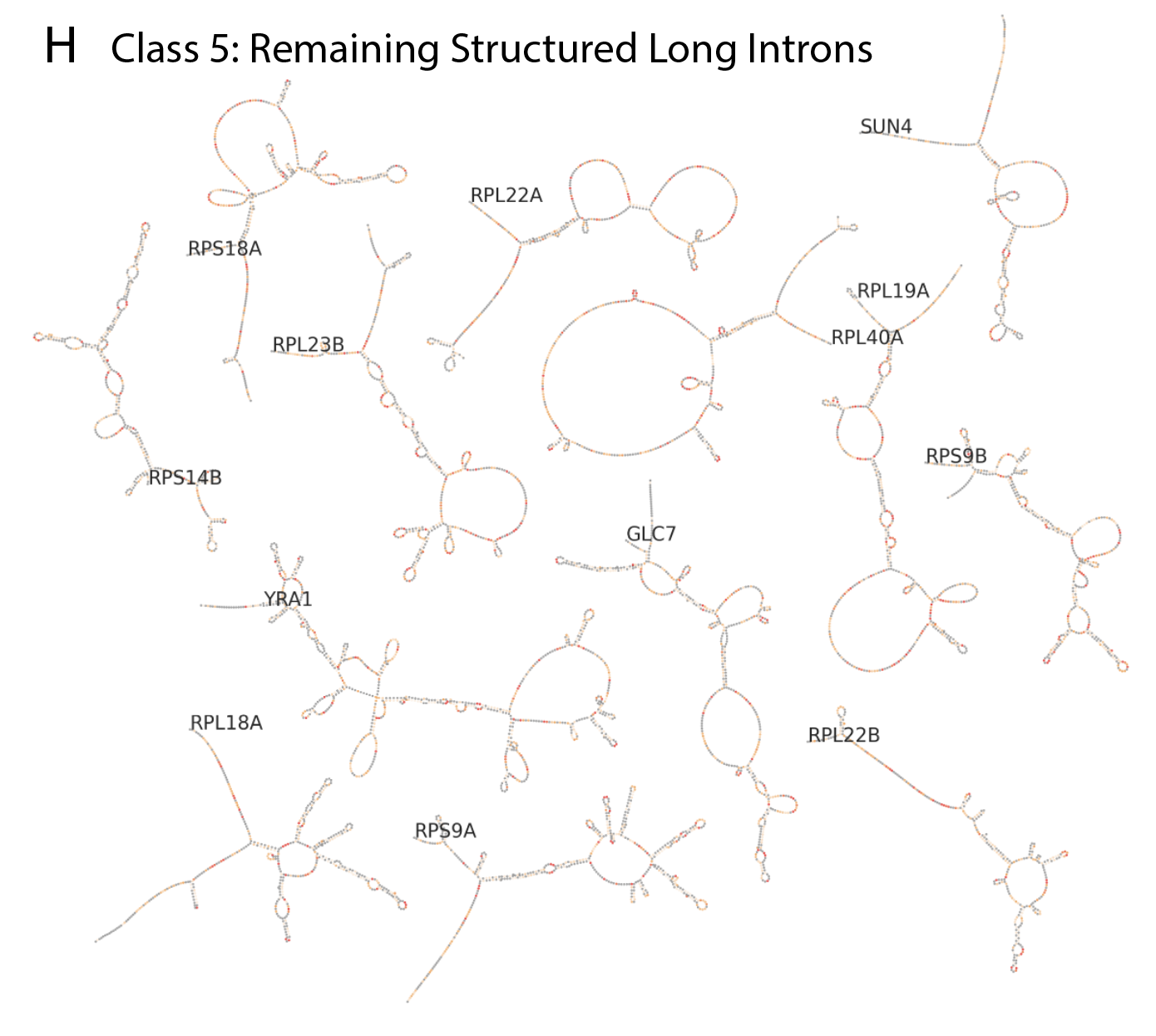

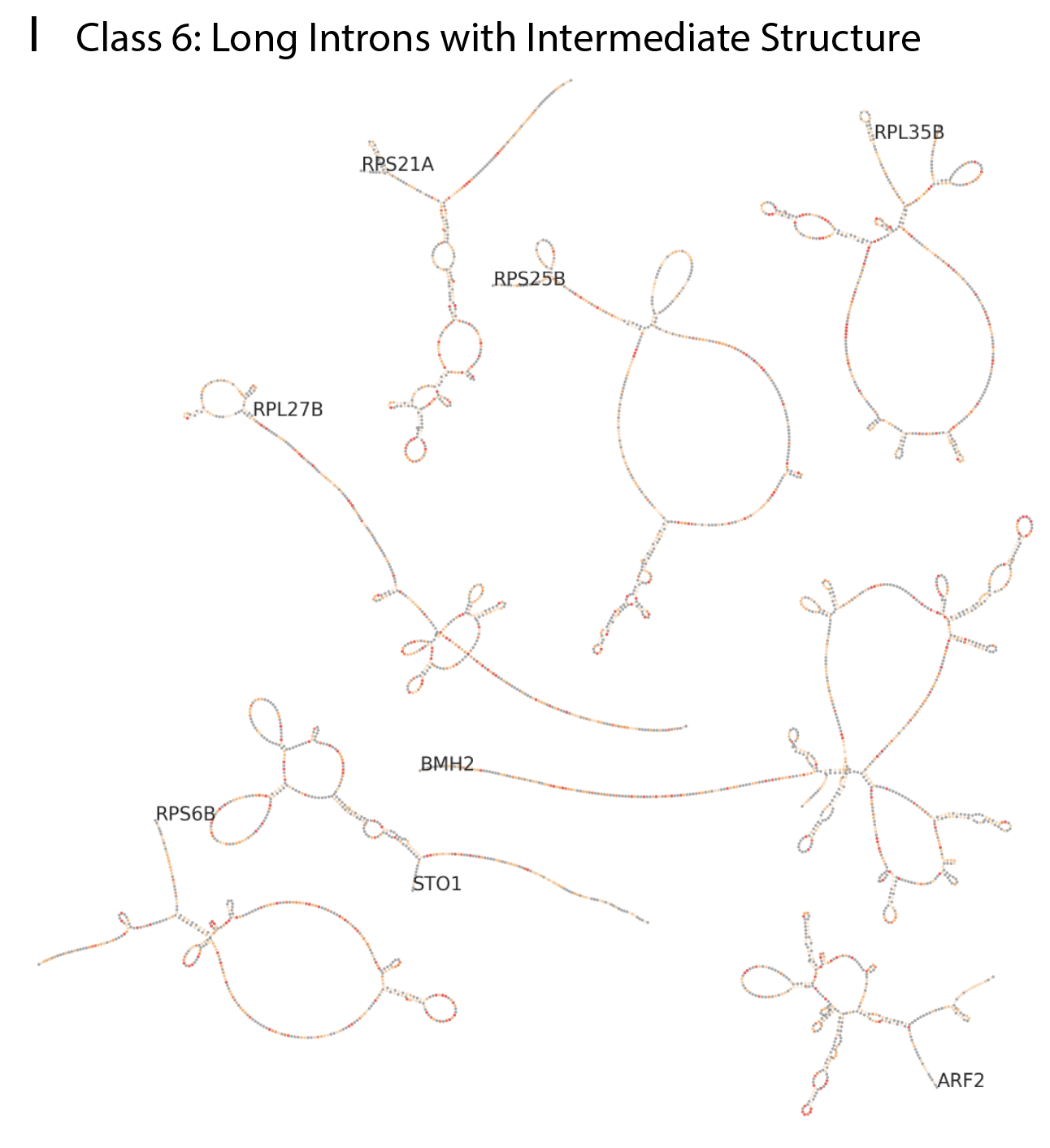

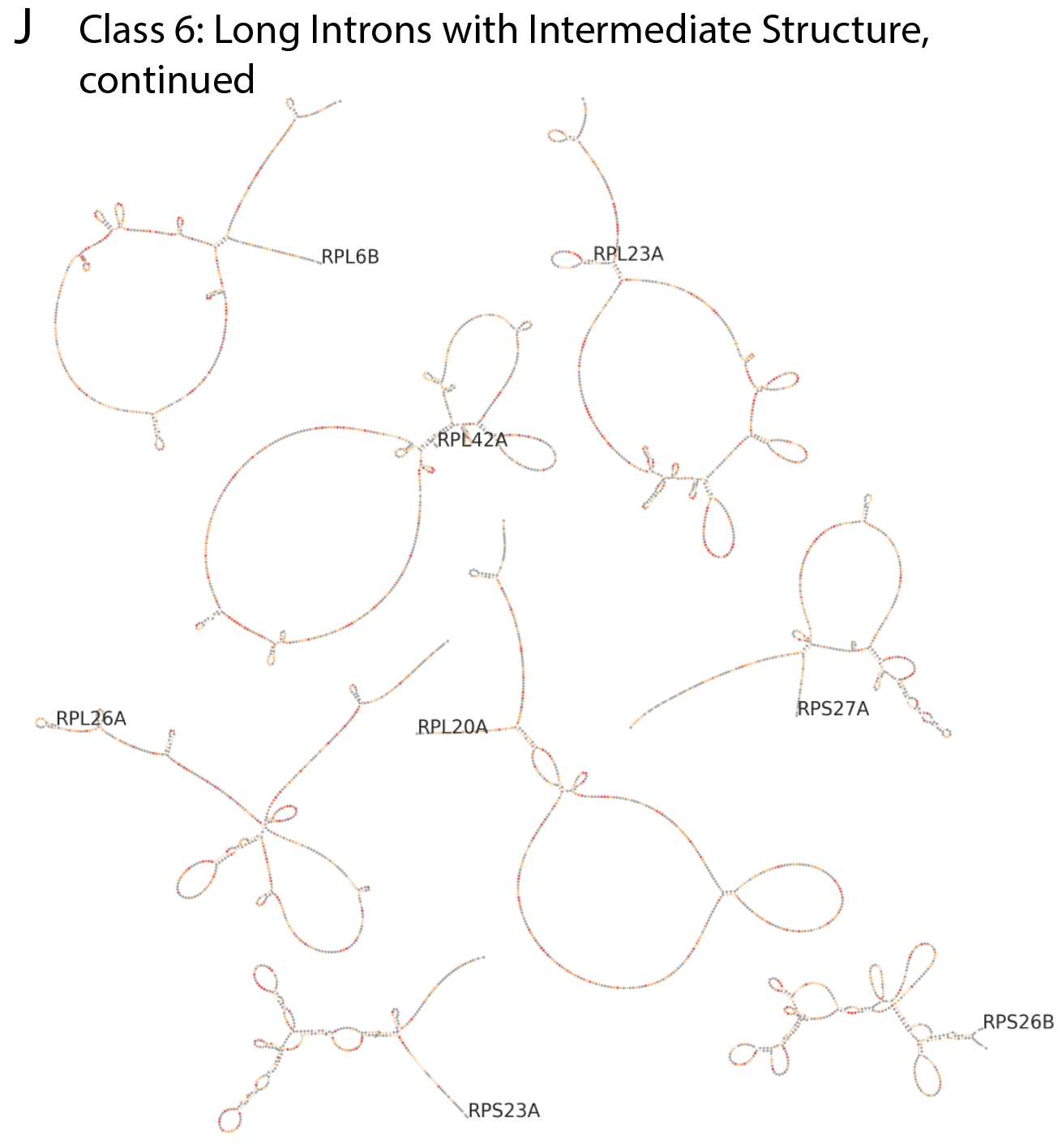

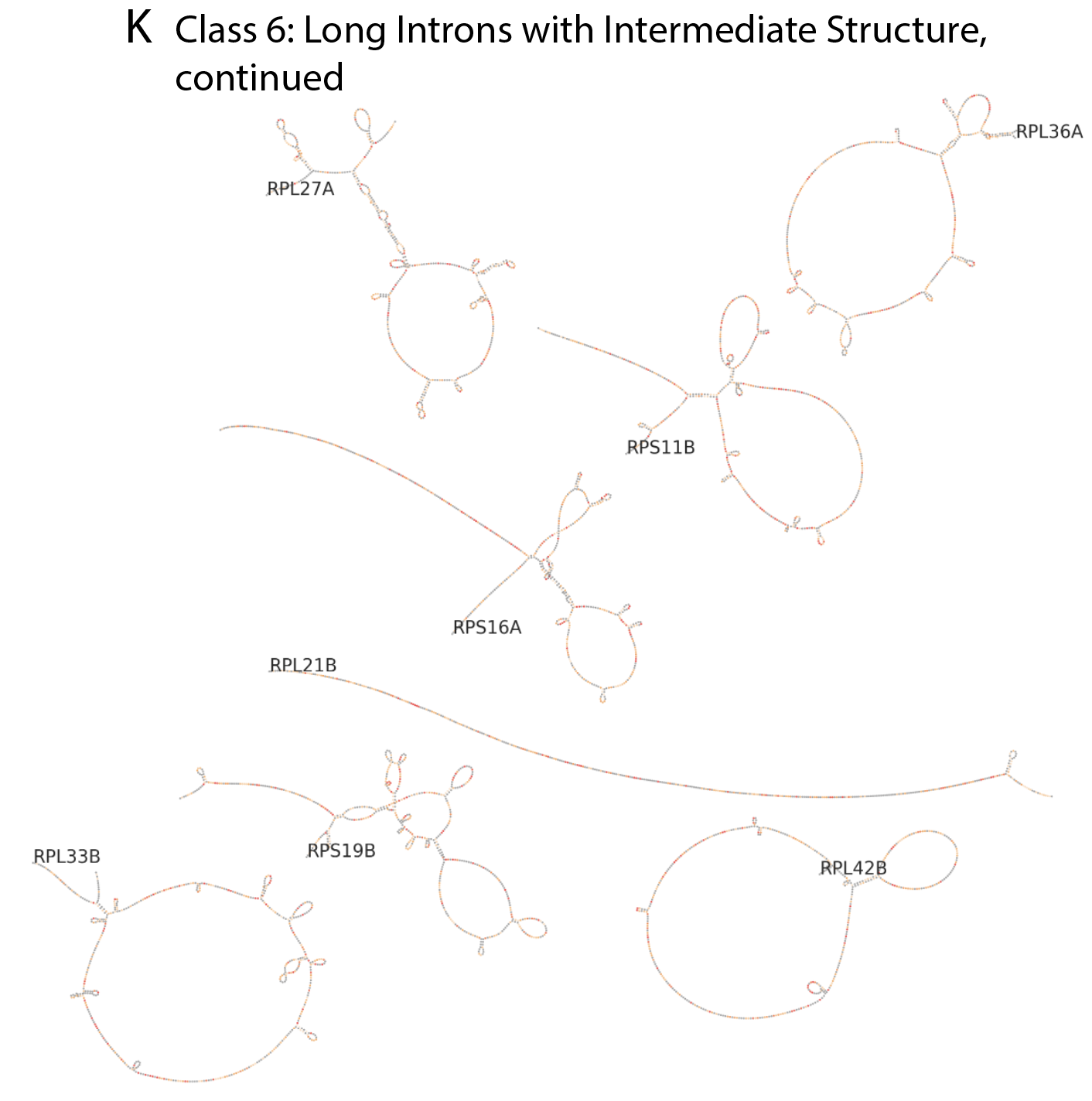

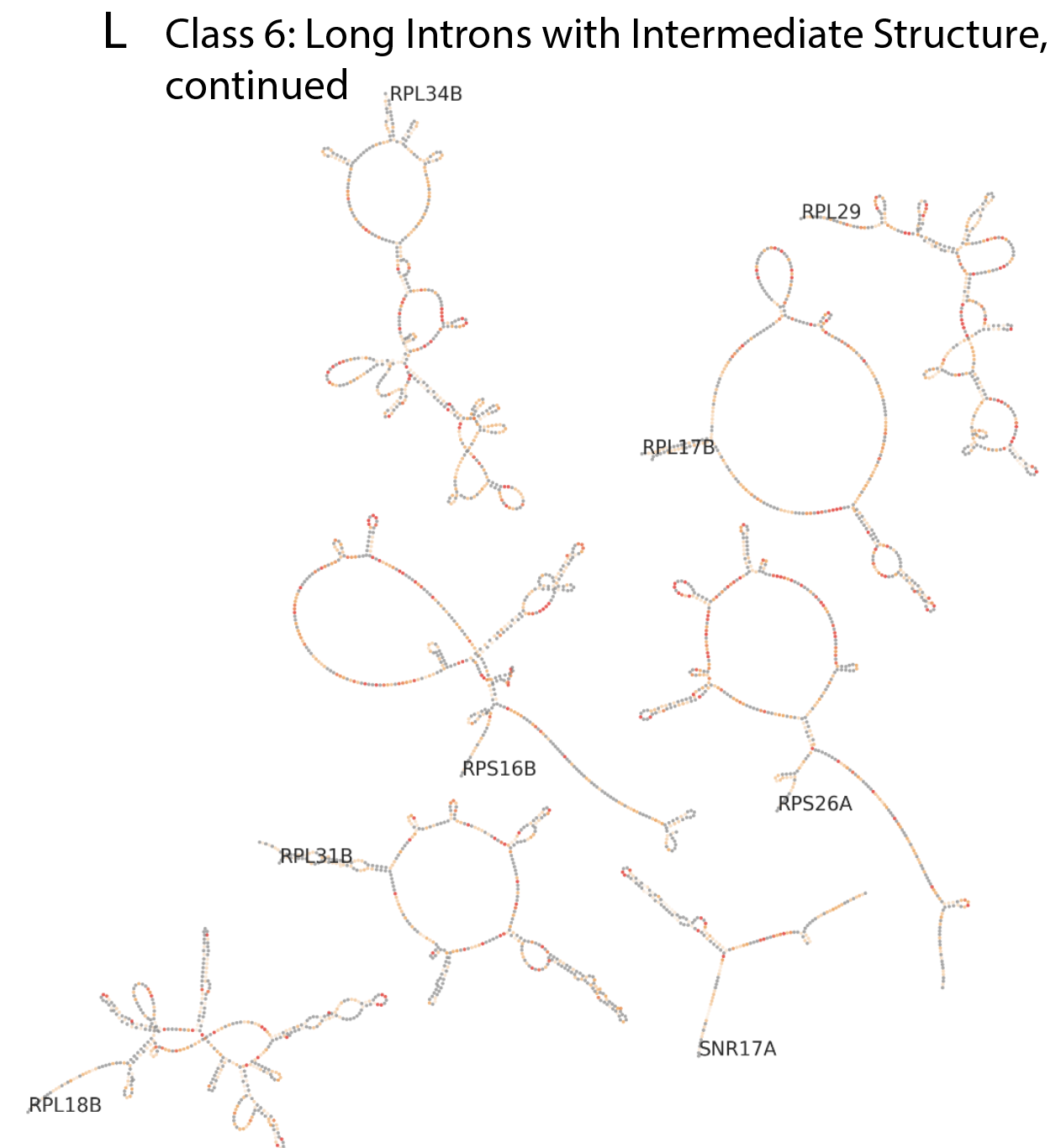

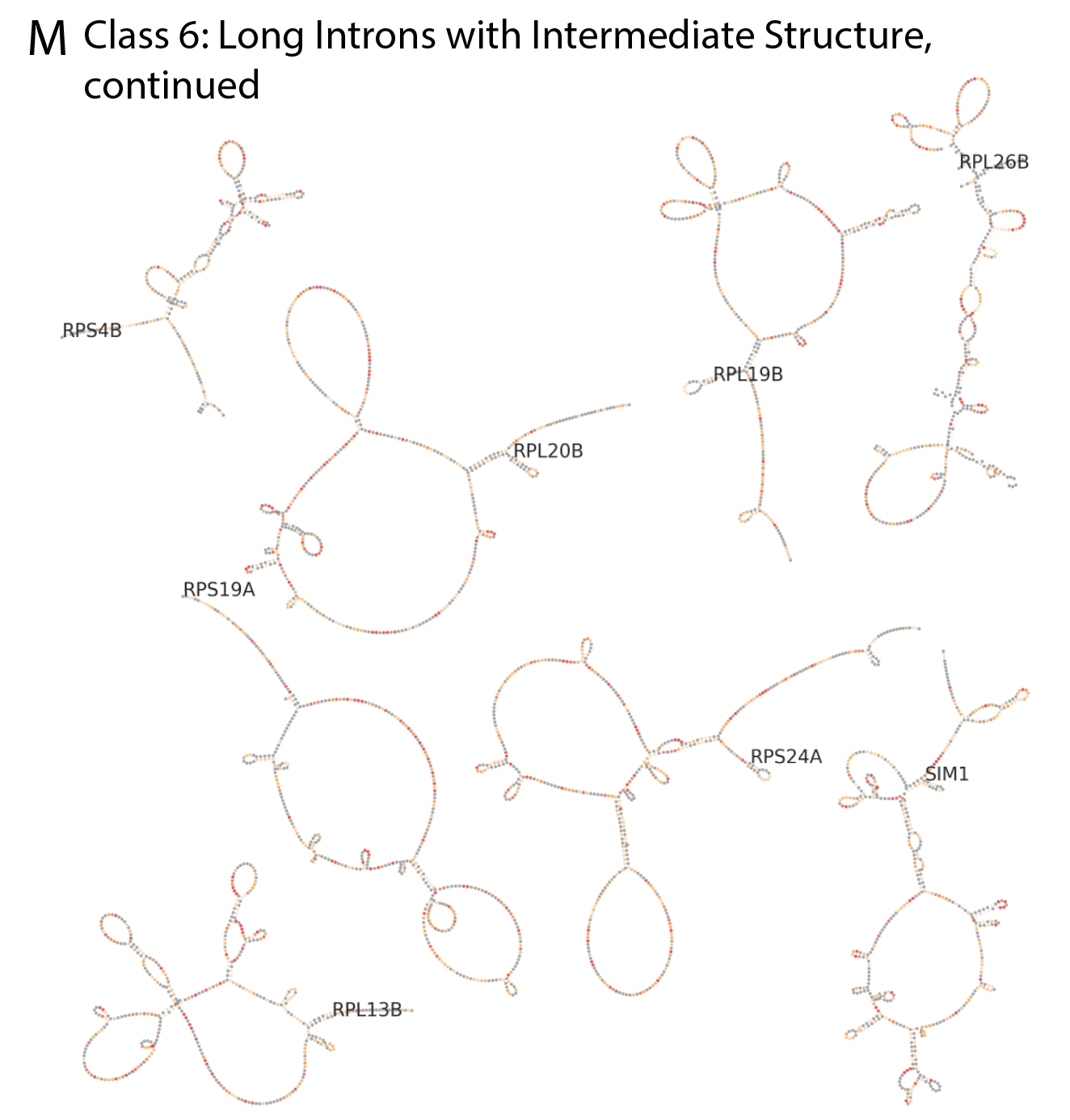

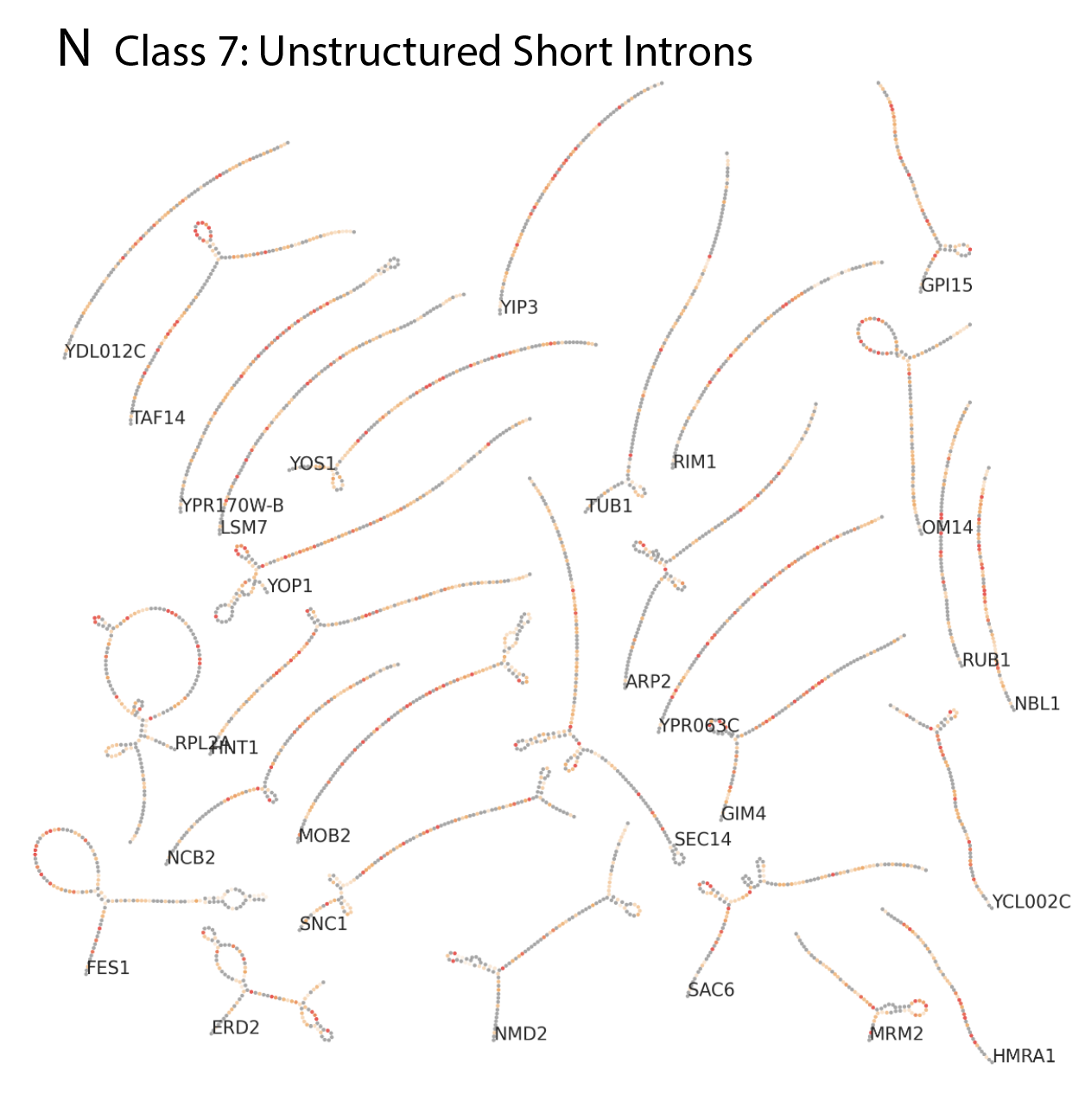

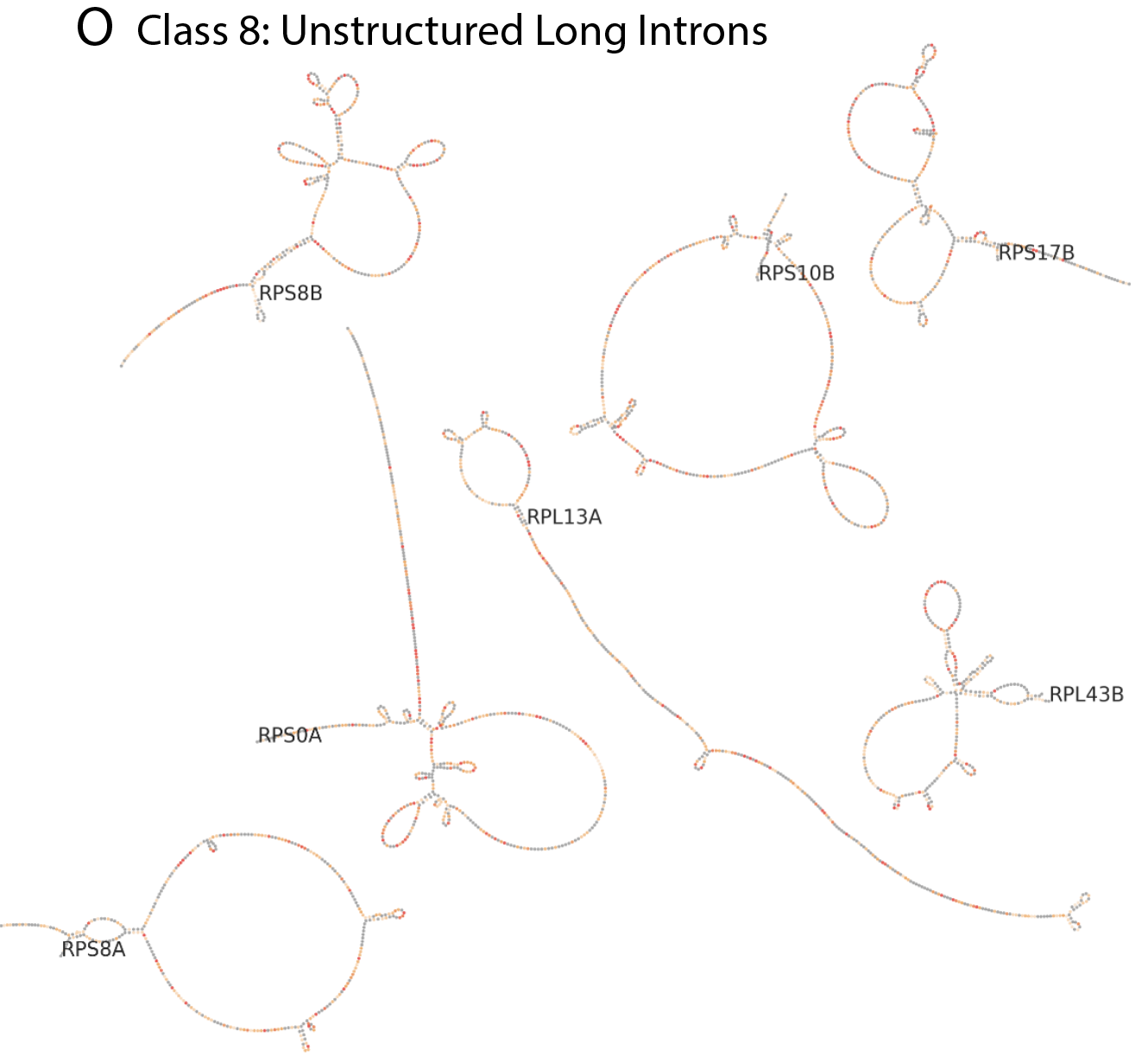

**Figure S28:** Secondary structures and overlaid DMS reactivity profiles for all introns with sufficient coverage from DMS-MaPseq. Introns are grouped into classes from hierarchical clustering (Fig. 5). Secondary structures are depicted schematically using RiboGraphViz (<https://github.com/DasLab/RiboGraphViz>). Gene names are included for each intron, with all labels at the 5’ ends of introns.

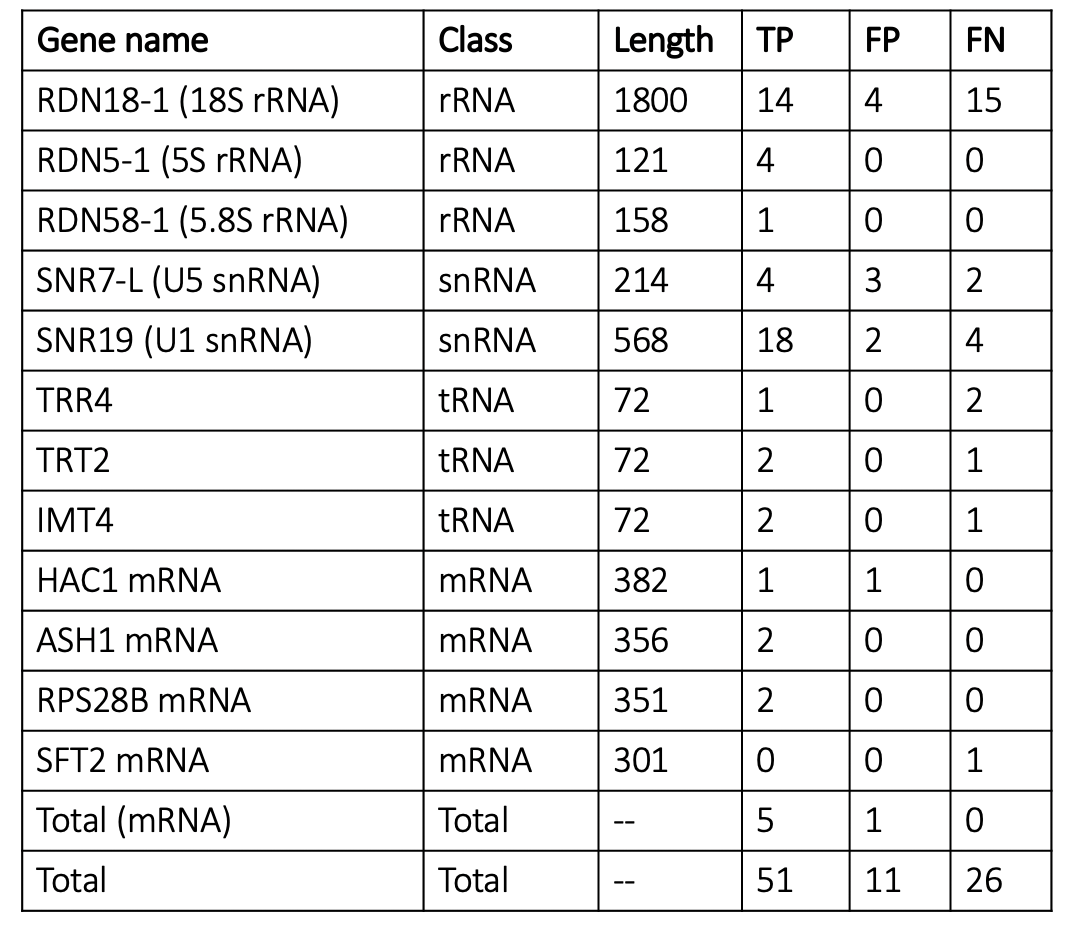

**Table S2:** True positive, false positive, and false negative stem predictions across a set of positive control structures including rRNAs, snRNAs, tRNAs, and mRNA segments. Secondary structure predictions are made using a 70% helix confidence estimate cutoff from bootstrapping DMS reactivity values. Stems are included if they include at least 5 base-pairs. The ground truth secondary structures for the 5S, 5.8S, and 18S rRNA were obtained from a high-resolution X-ray crystallography structure of the eukaryotic ribosome (PDB ID: 4V88)^21^. Structures for the U5 snRNA and U1 snRNA were obtained from Nguyen, et. al. (2016)^22^ and Li, et. al. (2017)^20^ respectively. Rfam-derived secondary structures^23^ served as ground truth structures for the four tRNA structures, and structures for mRNA segments in HAC1, ASH1, RPS28B, and SFT2 were obtained from Zubradt, et al. (2017)^24^.
